# TROP2-targeting chimeras (TRTACs) for tumor-selective membrane protein degradation and enhanced drug delivery

**DOI:** 10.64898/2026.08.19.745708

**Authors:** Luping Chen, Xinying Fu, Wenqian Dong, Xiaobei Deng, Simin Chen, Fujun Wang, Jian Zhao, Shiqun Shao, Liqiang Fan, Jingyu Zhang, Lixin Zhang

## Abstract

Extracellular targeted protein degradation (eTPD) systems typically utilize lysosome-targeting receptors (LTRs) to mediate internalization and lysosomal degradation of extracellular and membrane proteins. While multiple LTRs have been discovered, there remains a compelling need to seek for new LTRs, particularly those with clear clinical relevance, to expand the therapeutic potential of eTPD. Here we report trophoblast cell surface antigen-2 (TROP2), a clinically validated tumor-associated antigen, as a promising tumor-selective LTR. We engineer TROP2-targeting chimeras (TRTACs) by genetically fusing a TROP2-binding nanobody to nanobodies against specific target proteins. We show that TRTACs can induce tumor cell-selective degradation of diverse membrane proteins, including epithelial growth factor receptor (EGFR), human epithelial growth factor receptor 2 (HER2), and programmed death-ligand 1 (PD-L1). The EGFR-targeted TRTAC significantly inhibits tumor cell proliferation and shows potent antitumor activity *in vivo*. We further design TRTAC-drug conjugates (TRTAC-DCs) by attaching cytotoxic payloads to TRTACs, enabling targeted protein degradation together with enhanced drug delivery. TRTAC-DCs show significantly enhanced activity against HER2- and EGFR-positive tumors both *in vitro* and *in vivo*, with minimal toxicity observed in normal tissues. These findings establish TROP2 as a robust LTR and provide a versatile eTPD platform with profound translational potential for tumor treatment.

## 1. Introduction

Extracellular targeted protein degradation (eTPD) has emerged as a promising therapeutic modality that hijacks the cell’s endolysosomal pathway to degrade extracellular and membrane proteins ^[1, 2]^. This strategy typically uses heterobifunctional chimeras to bridge proteins of interest (POIs) with lysosome-targeting receptors (LTRs) or membrane-associated E3 ligases, thereby inducing internalization and lysosomal degradation of POIs. Since LTR engagement dictates cellular uptake, intracellular routing, recycling kinetics, tissue specificity, and overall degradation efficacy ^[3]^, identifying effective LTRs has become a major focus in the eTPD field. To date, ∼20 LTRs have been reported for eTPD development, including the cation-independent mannose-6-phosphate receptor (CI-M6PR)/insulin-like growth factor-2 receptor (IGF2R) ^[4–7]^, asialoglycoprotein receptor (ASGPR) ^[8–11]^, transferrin receptor (TfR) ^[12–14]^, among others ^[15–27]^. However, many of these receptors lack tissue or tumor selectivity and are susceptible to adaptive downregulation ^[28]^. Moreover, not all receptors mediate internalization and lysosomal trafficking with the same efficiency ^[12, 29]^, and not all POIs are amenable to existing degraders possibly due to incompatible binding kinetics for productive ternary complex formation ^[6]^. Expanding the repertoire of LTRs can increase the optionality of eTPD tools, unlocking greater opportunities for improved tissue-selectivity, enhanced degradation potency, a broader target scope, and strategic avenues to overcome resistance mechanisms.

Antibody-drug conjugates (ADCs) are an emerging class of biopharmaceuticals that leverage the targeting specificity of antibodies to selectively deliver cytotoxic drugs to diseased tissues ^[30]^. When binding to cell-surface antigens, ADCs are internalized through receptor-mediated endocytosis and trafficked to lysosomes, where the active payloads are released to kill cells ^[31–33]^. This process closely resembles that of most eTPD approaches (Figure 1A). We reasoned that these clinically validated ADC targets could be repurposed as tumor-selective LTRs for developing new eTPD systems. Moreover, these degraders can be further engineered into degrader-drug conjugates (DDCs) by conjugating cytotoxic drugs, which can enable tumor-targeted drug delivery while simultaneously inducing extracellular target degradation. Notably, current DDC systems are hampered by limited tumor specificity, as well as suboptimal tumor penetration due to the large size of antibody scaffolds ^[34–37]^. Although small-molecule DDCs can improve tissue penetration ^[38]^, their target scope is limited because only a small portion of membrane proteins have established small-molecule binders with sufficient affinity and specificity suitable for degrader engineering.

**Figure 1.**
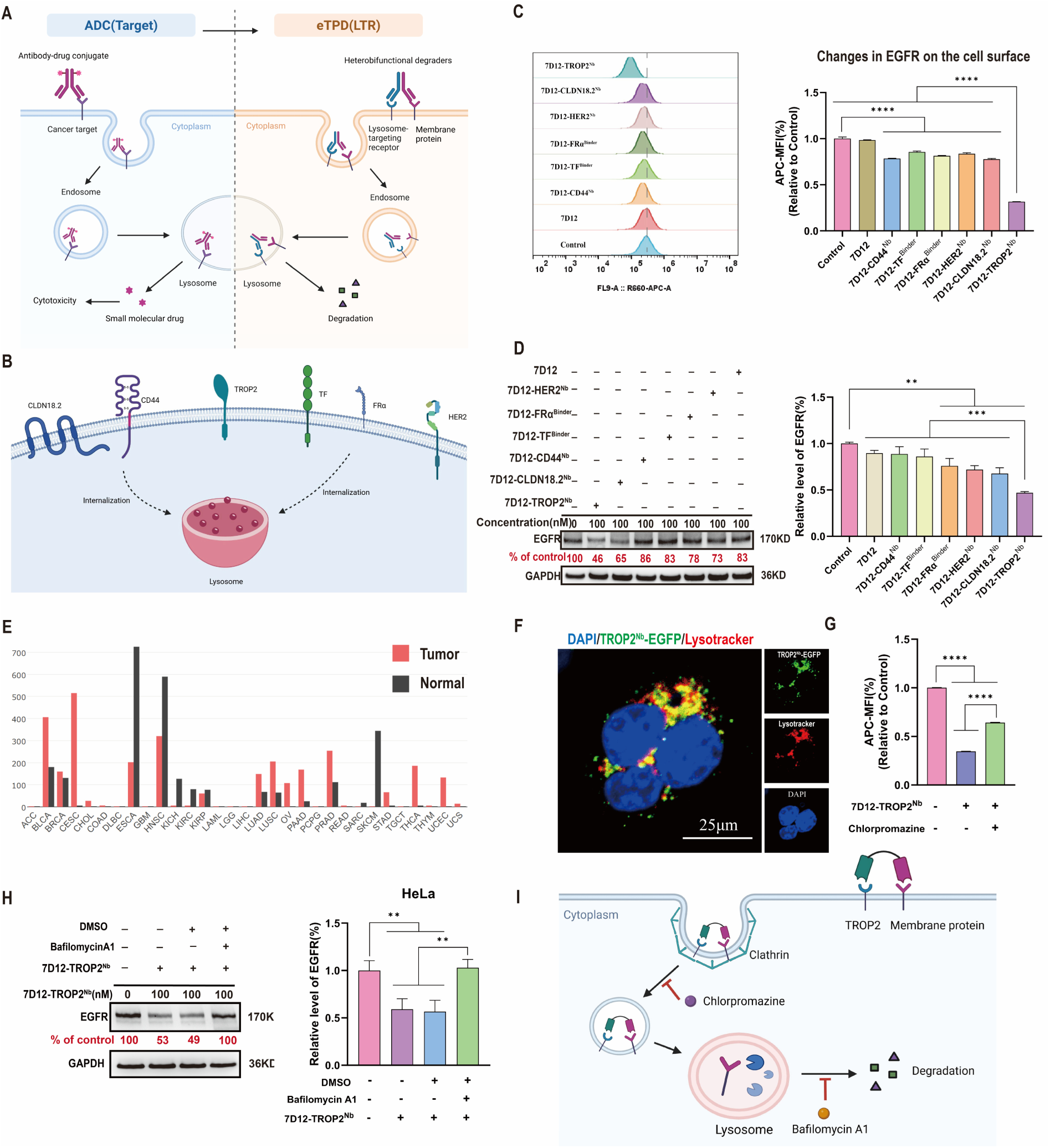
Design and mechanistic validation of TROP2-mediated EGFR internalization and lysosomal degradation. (**A**) Schematic illustration of ADC and eTPD. (**B**) Clinically relevant ADC targets screened for new LTR identification. (**C**) Flow cytometry analysis of EGFR levels in HeLa cells treated with 500 nM of 7D12, 7D12-CD44^Nb^, 7D12-TF^Binder^, 7D12-FRα^Binder^, 7D12-HER2^Nb^, 7D12-CLDN18.2^Nb^, or 7D12-TROP2^Nb^ for 24 h. (**D**) Western blot analysis of EGFR levels in HeLa cells treated with 100 nM of 7D12, 7D12-CD44^Nb^, 7D12-TF^Binder^, 7D12-FRα^Binder^, 7D12-HER2^Nb^, 7D12-CLDN18.2^Nb^, or 7D12-TROP2^Nb^ for 24 h. (**E**) GEPIA2 database analysis of TACSTD2 expression across multiple tumor tissues and matched normal tissues. (**F**) Live-cell confocal imaging of HeLa cells treated with 1 μM of TROP2^Nb^-EGFP for 6 h, then incubated with DAPI and LysoTracker Red for 30 min. Scale bar, 25 μm. (**G**) Flow cytometry analysis of EGFR levels in HeLa cells treated with 100 nM of 7D12-TROP2^Nb^ for 24 h in the presence or absence of 20 μM chlorpromazine, then stained with APC-conjugated anti-EGFR for 1 h. (**H**) Western blot analysis of EGFR protein levels in HeLa cells treated with 100 nM of 7D12-TROP2^Nb^ for 24 h in the presence or absence of 50 nM of bafilomycin A1. (**I**) Schematic illustration of TRTAC-mediated membrane protein degradation. Data are presented as mean ± standard deviation (SD) (n = 3 biologically independent replicates) when relevant. Data in **E** are shown as mean values. *P* values were determined by one-way analysis of variance (ANOVA) with Turkey’s *post hoc* test. \*\**P* < 0.01, \*\*\**P* < 0.001, \*\*\*\**P* < 0.0001.

Herein, we report an effective strategy to uncover new tumor-selective LTRs from clinically validated ADC targets. Through rigorous screening, we identified trophoblast cell surface antigen-2 (TROP2) as a promising candidate and developed TROP2-targeting chimeras (TRTACs) and TRTAC-drug conjugates (TRTAC-DCs). TRTACs are modularly produced by genetically fusing a TROP2-binding nanobody to POI-specific nanobodies. These nanobody-based degraders exhibited high tumor permeability and tumor cell-selective degradation of diverse cell-surface targets, including epithelial growth factor receptor (EGFR), human epidermal growth factor receptor 2 (HER2), and programmed death-ligand 1 (PD-L1). EGFR-specific TRTACs significantly inhibited tumor cell proliferation and showed strong antitumor activity *in vivo*. When conjugated with cytotoxic payloads, TRTAC-DCs demonstrated enhanced potency against HER2- and EGFR-positive tumors both *in vitro* and *in vivo*, with no systemic toxicity observed. This work provides a facile strategy for discovering tumor-selective LTRs and highlights TRTAC and TRTAC-DC platforms with promising therapeutic potentials.

## 2. Results

### 2.1. Identification of TROP2 as a promising tumor-selective LTR

To identify new LTRs, we selected six clinically successful ADC targets, including HER2 ^[39]^, TROP2 ^[40]^, FRα ^[41]^, tissue factor (TF) ^[42]^, claudin-18.2 (CLDN18.2) ^[43]^, and CD44 ^[44]^ (Figure 1B). Their capabilities in supporting target degradation were evaluated by using EGFR as a model substrate ^[4]^. We engineered heterobifunctional chimeric molecules by genetically fusing nanobodies or targeting peptides against each candidate receptor to the EGFR-binding nanobody 7D12 (Figure S1A). The six candidates were compared within the same cellular background to minimize cell line-associated variability. Because HeLa cells and SKBR3 cells express all six receptors (Figure S1B,C), both cell lines were selected to assess changes in cell-surface EGFR levels induced by these chimeras. Flow cytometry showed that, compared to the 7D12 control, 7D12-CLDN18.2^Nb^ and 7D12-TROP2^Nb^ consistently decreased EGFR levels in both cell lines, with 7D12-TROP2^Nb^ exhibiting the highest activity (Figure 1C and Figure S1D). While the other candidates induced mild EGFR reduction in HeLa cells, this activity was not observed in SKBR3 cells. These results were further corroborated by Western blot analysis (Figure 1D and Figure S1E). Notably, TROP2 expression levels in HeLa and SKBR3 cells were not uniquely higher than the other receptors (Figure S1B), suggesting that the superior degradation performance mediated by TROP2 was not simply due to higher receptor abundance. GEPIA2 database analysis revealed extensive overexpression of TROP2 across multiple tumor types, including cervical squamous cell carcinoma, endocervical adenocarcinoma, and pancreatic adenocarcinoma (Figure 1E). Furthermore, TROP2 overexpression correlates with poor prognosis in cervical cancer, pancreatic cancer, lung adenocarcinoma, and ovarian cancer (Figure S1F). Prior studies have also indicated that tumor cells exhibit stronger TROP2 internalization compared to normal tissues, and overexpressed TROP2 in tumor cells can be preferentially engaged ^[45]^. Collectively, these data prioritize TROP2 as a promising tumor-selective LTR candidate.

Next, we proceeded to validate TROP2 as an effective LTR. Flow cytometry revealed a time-dependent increase in cellular association and internalization of enhanced green fluorescence protein (EGFP)-fused TROP2^Nb^ (TROP2^Nb^-EGFP) (Figure S1G). Confocal microscopy confirmed that the internalized TROP2^Nb^-EGFP was primarily trafficked to lysosomes (Figure 1F). Pre-incubation of HeLa cells with chlorpromazine significantly blocked 7D12-TROP2^Nb^-mediated EGFR internalization, suggesting that TRTAC-induced internalization relies on clathrin-mediated endocytosis (Figure 1G). Inhibiting lysosomal acidification with bafilomycin A1 rescued intracellular EGFR levels (Figure 1H) without affecting cell-surface EGFR internalization (Figure S1H), confirming the lysosomal degradation of EGFR following TRTAC-induced internalization (Figure 1I). Together, these results validate TROP2 as an effective LTR for developing extracellular degraders.

#### Characterization of TRTAC-mediated EGFR degradation *in vitro*

Next, we detailed the characterization of TROP2-mediated EGFR degradation. An IGF2-fused counterpart (7D12-IGF2) was employed as a benchmark positive control) ^[5]^. Western blot and flow cytometry analyses revealed that 7D12-TROP2^Nb^ outperformed 7D12-IGF2 in EGFR degradation (Figure 2A and Figure S2A). Neither 7D12 alone nor the combination of free 7D12 and TROP2^Nb^ decreased EGFR levels, indicating that the chimeric architecture is indispensable for TRTAC-mediated degradation. We demonstrated that this degradation took place in a time- and concentration-dependent manner, with no hook effect observed at the tested concentration range (Figure 2B,C). Notably, incubation with 100 nM of 7D12-TROP2^Nb^ for 24 h decreased intracellular EGFR levels by 47∼59%, while higher concentrations led to greater degradation (Figure 2B,C and Figure S2B).

**Figure 2.**
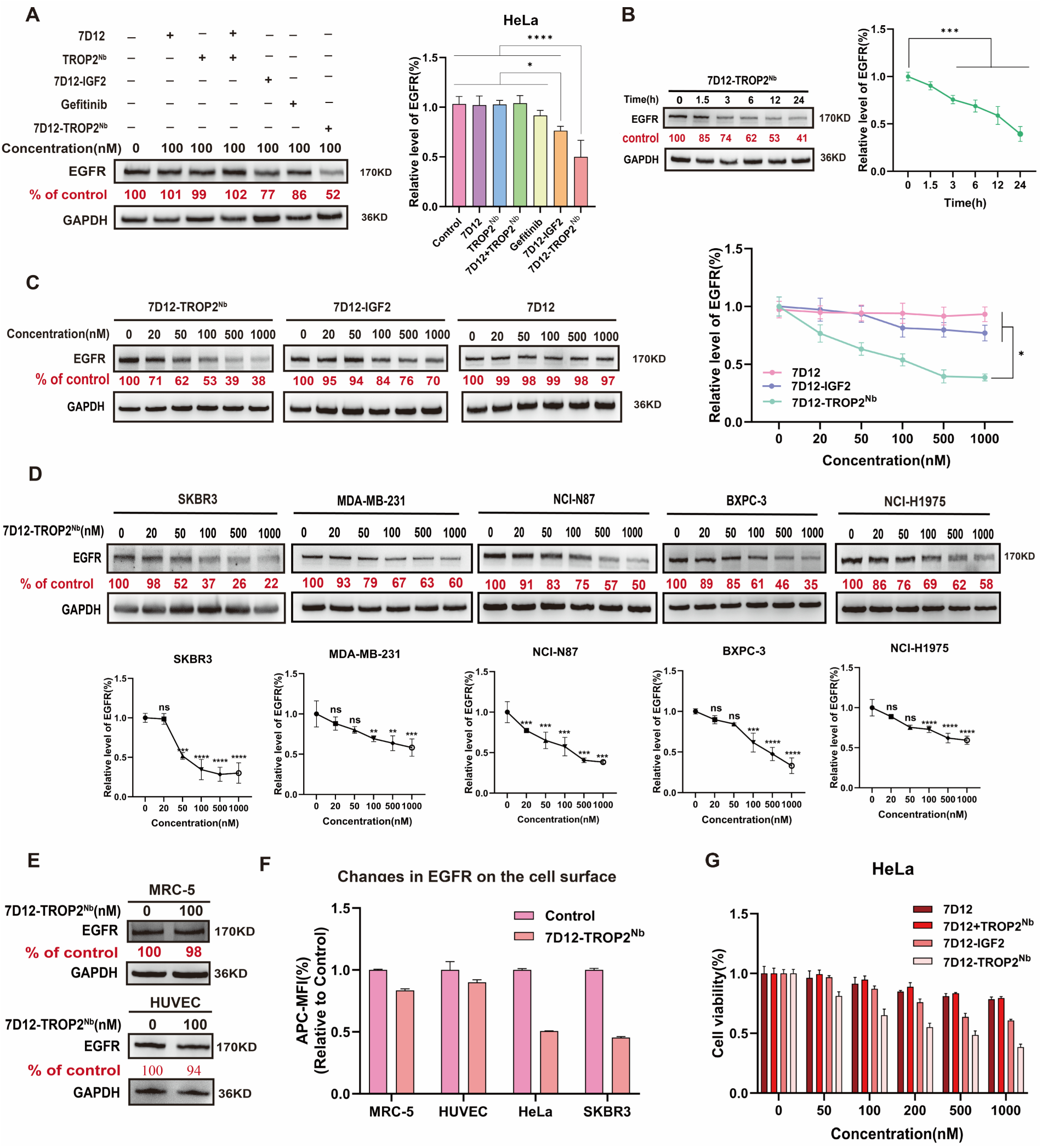
Characterization of 7D12-TROP2^Nb^-mediated EGFR degradation *in vitro*. (**A-C**) Western blot analysis of EGFR levels in HeLa cells treated for 24 h with 100 nM of 7D12, TROP2^Nb^, 7D12 + TROP2^Nb^, 7D12-IGF2, gefitinib, or 7D12-TROP2^Nb^ (**A**), or treated with 100 nM of 7D12-TROP2^Nb^ for 0, 1.5, 3, 6, 12, or 24 h (**B**), or treated for 24 h with different concentrations of 7D12-TROP2^Nb^, 7D12-IGF2, or 7D12 (**C**). (**D** and **E**) Western blot analysis of EGFR levels in SKBR3, MDA-MB-231, NCI-N87, BxPC-3, or NCI-H1975 cells treated with different concentrations of 7D12-TROP2^Nb^ for 24 h (**D**) or in MRC-5 and HUVEC cells treated with 100 nM of 7D12-TROP2^Nb^ for 24 h (**E**). (**F**) Flow cytometry analysis of MRC-5, HUVEC, HeLa, or SKBR3 cells treated with PBS or 100 nM of 7D12-TROP2^Nb^ for 24 h, then stained with APC-conjugated anti-EGFR for 1 h. (**G**) MTT assay of HeLa cells treated with serial concentrations of 7D12, 7D12 + TROP2^Nb^, 7D12-IGF2, or 7D12-TROP2^Nb^ for 7 d. Data are presented as mean ± SD (n = 3 biologically independent replicates) when relevant. *P* values were determined by one-way ANOVA with Turkey’s *post hoc* test. ns, no significance; \**P* < 0.05; \*\**P* < 0.01; \*\*\**P* < 0.001; \*\*\*\**P* < 0.0001.

We then evaluated the tumor selectivity of TRTACs. Six cancer cell lines (HeLa, SKBR3, MDA-MB-231, NCI-N87, BxPC-3, and H1975), and two normal cell lines, lung fibroblasts (MRC-5) and human umbilical vein endothelial cells (HUVECs), were included for examination. All cancer cell lines express both EGFR and TROP2, with SKBR3, NCI-N87, and BxPC-3 cells showing higher TROP2 levels (Figure S2C) ^[46]^. 7D12-TROP2^Nb^ induced EGFR degradation in all cancer cell lines, particularly in SKBR3, NCI-N87, and BxPC-3 cells (Figure 2D), suggesting that higher TROP2 abundance may enhance degradation efficiency. While MRC-5 and HUVEC cells also showed appreciable TROP2 expression (Figure S2D), 7D12-TROP2^Nb^ at 100 nM produced almost no detectable EGFR degradation in both normal cells (Figure 2E,F). MTT assays revealed that 7D12-TROP2^Nb^ significantly inhibited the proliferation of HeLa (Figure 2G), but showed no effects on normal cell lines (Figure S2E). These results confirm the tumor selectivity of TRTAC-mediated EGFR degradation.

To rule out the possibility that this activity was unique to this specific nanobody (TROP2^Nb^), we involved two additional TROP2 nanobodies (TROP2^NbV-5^ and TROP2^Nb108^) in TRTAC design. These nanobodies were similarly fused to 7D12 to generate EGFR degraders. The new constructs effectively reduced cell-surface EGFR levels (Figure S2F) and suppressed cell proliferation (Figure S2G), albeit with lower potencies than 7D12-TROP2^Nb^. We explored whether this variation was associated with differences in receptor affinity or epitope recognition. Molecular docking analysis suggested that these nanobodies recognized distinct predicted epitopes on TROP2, with TROP2^Nb^ showing the lowest predicted binding energy of −71.65 kcal mol^-1^ and hence the strongest predicted interaction (Figure S2H). To experimentally validate receptor affinity, we performed the enzyme-linked immunosorbent assay (ELISA) using EGFP as a detection tag and confirmed that TROP2^Nb^ had the highest affinity for TROP2 (EC_50_: 45.26 nM) (Figure S2I). These data indicate the robustness of TROP2 as an LTR and that increasing affinity for TROP2 may further enhance degradation efficiency.

#### EGFR-targeted TRTAC suppresses HeLa xenograft growth *in vivo*

Given that 3D tumor models more faithfully recapitulate solid tumors compared to conventional 2D cultures ^[47]^, we investigated the degradation activity of 7D12-TROP2^Nb^ in 3D tumor spheroids prior to *in vivo* evaluation. Western blot analysis showed that 7D12-TROP2^Nb^ efficiently degraded EGFR in spheroids derived from HeLa, NCI-H1975, and SKBR3 cells, suggesting a high tumor permeability (Figure S3A,B). Compared to previously reported antibody-based chimeras, 7D12-TROP2^Nb^ has a substantially smaller molecular mass of ∼30 kDa, which may favor deeper penetration into solid tumors.

We next moved on to evaluate the *in vivo* therapeutic efficacy of 7D12-TROP2^Nb^. HeLa tumor-bearing nude mice were intravenously administered PBS, 7D12-IGF2, or 7D12-TROP2^Nb^ every two days (Figure 3A). Compared to 7D12-IGF2, 7D12-TROP2^Nb^ treatment more significantly inhibited tumor growth (Figure 3B-D). Throughout the treatment, no significant body weight change was observed, and H&E staining of major organs revealed no obvious abnormalities (Figure 3E and Figure S3C), indicating favorable biosafety profiles for both EGFR degraders. Immunohistochemical analysis of tumor tissues confirmed a marked reduction in EGFR levels after 7D12-TROP2^Nb^ treatment (Figure 3F). These results showed that 7D12-TROP2^Nb^ can induce EGFR degradation *in vivo* and produce substantial tumor suppression with low apparent toxicity.

**Figure 3.**
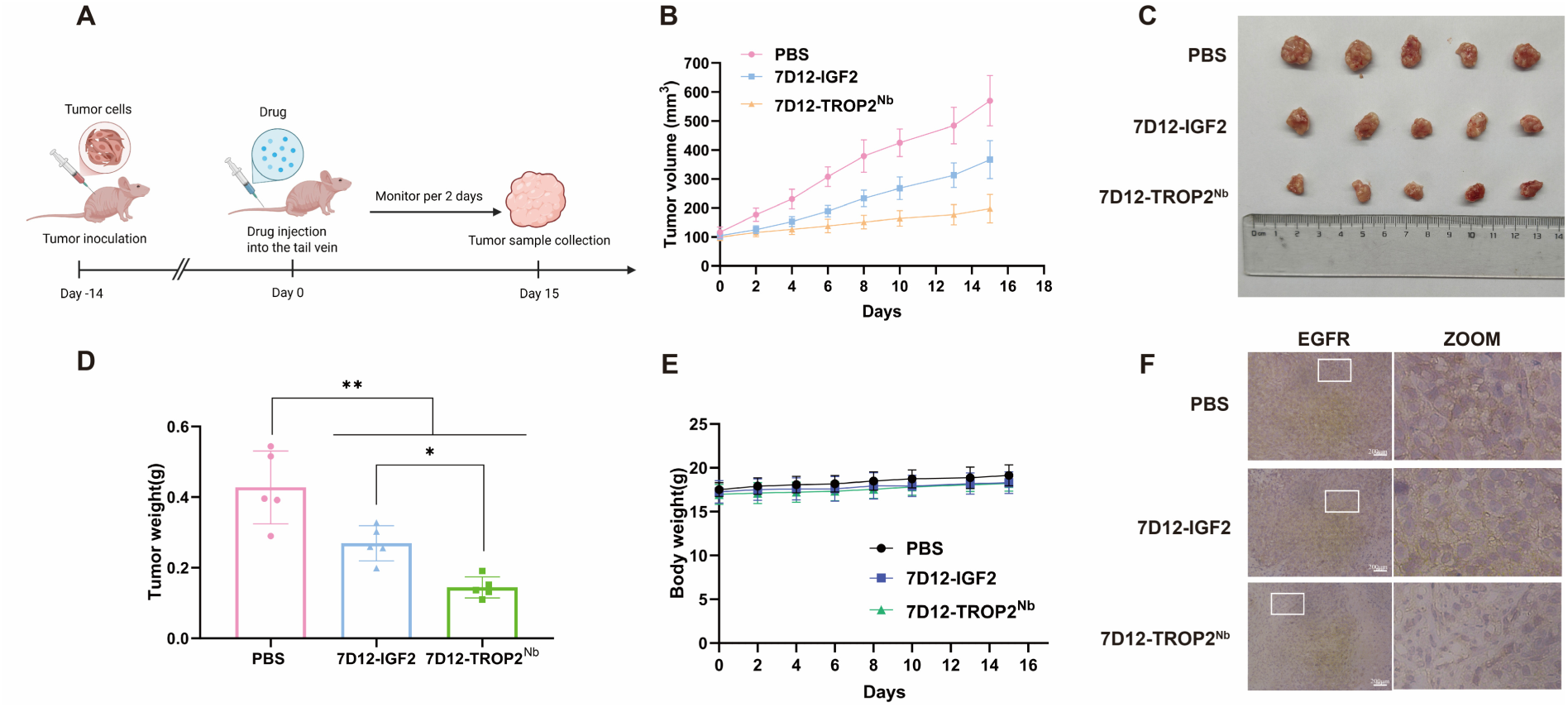
***In vivo* antitumor effects of 7D12-TROP2^Nb^.** (**A**) Schematic illustrating the experimental design to assess the antitumor efficacy of 7D12-TROP2^Nb^. HeLa tumor-bearing nude mice were *i.v.* administered PBS, 7D12-IGF2, or 7D12-TROP2^Nb^ at a 7D12-equiv. dose of 5 mg/kg every 2 days for 8 doses (n = 5 mice per group). (**B**) Tumor growth curves in each group. (**C**) Photograph of excised tumors from each group. (**D**) Average tumor weight of each group measured at the endpoint. (**E**) Body weight change of mice in each group. (**F**) Representative immunohistochemical staining of EGFR in tumor sections. Scale bar: 200 μm. Data are presented as mean ± SD. *P* values were determined by one-way ANOVA with Turkey’s *post hoc* test. \**P* < 0.05; \*\**P* < 0.01.

#### Expanding the target scope of TRTACs

Having demonstrated the target scope of TRTAC towards EGFR, we next proceeded to assess its efficacy in degrading other extracellular targets (Figure 4A). We first focused on soluble extracellular EGFP. The degrader was constructed by fusing an EGFP-binding nanobody (EGFP^Nb^) to TROP2^Nb^. Cells were co-incubated with EGFP and EGFP^Nb^-TROP2^Nb^ for 6 h and then chased in drug-free medium. Flow cytometry showed that SKBR3 cells efficiently internalized EGFP and subsequently degraded the internalized protein during the chase period, whereas the control group showed no appreciable uptake (Figure 4B). Pretreatment with bafilomycin A1 ablated EGFP degradation, confirming that this process is lysosome-dependent. Afterward, we turned to membrane proteins PD-L1 and HER2. The corresponding TRTACs were generated by fusing the PD-L1 nanobody (KN035) or the HER2 nanobody (11A4) to TROP2^Nb^. KN035-TROP2^Nb^ and 11A4-TROP2^Nb^ were evaluated in PD-L1-positive MDA-MB-231 cells and HER2-positive SKBR3 cells, respectively. Western blot and flow cytometry analyses revealed that KN035-TROP2^Nb^ or 11A4-TROP2^Nb^ degraded PD-L1 or HER2 more efficiently than IGF2-based controls, IGF2-KN035, or IGF2-11A4 (Figure 4C-J), in a dose-dependent manner (Figure 4K-N and Figure S4A,B). Notably, 11A4-TROP2^Nb^ triggered potent HER2 internalization and degradation in SKBR3 cells at a concentration as low as 4 nM, resulting in significantly arrested cell proliferation (Figure 4O). HER2 degradation was also observed in NCI-N87 cells after 11A4-TROP2^Nb^ treatment (Figure S4C). Again, bafilomycin A1 treatment blocked PD-L1 or HER2 degradation, substantiating the lysosome-dependence of TRTAC activity (Figure S4D,E). These results demonstrate the broad target scope of the TRTAC platform.

**Figure 4.**
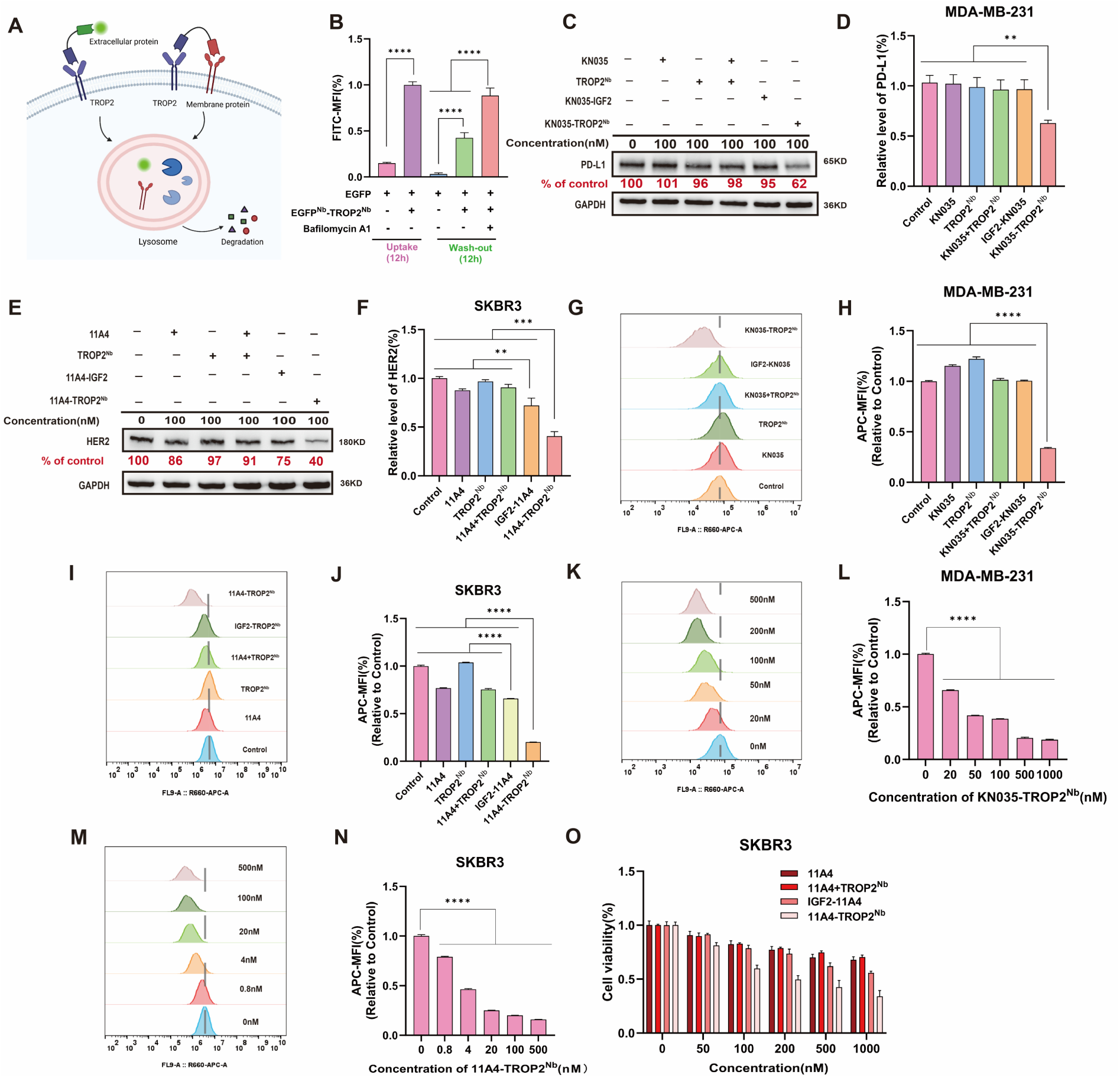
TRTAC-mediated degradation of multiple soluble and cell-surface targets. (**A**) Schematic illustration of TRTAC-mediated degradation of soluble and membrane-bound proteins. (**B**) Flow cytometry analysis of SKBR3 cells incubated for 12 h with soluble EGFP in the presence or absence of EGFP^Nb^-TROP2^Nb^ (100 nM) and bafilomycin A1 (50 nM). (**C** and **D**) Western blot analysis (**C**) and quantification (**D**) of PD-L1 levels in MDA-MB-231 cells treated for 24 h with 100 nM of KN035, TROP2^Nb^, IGF2-KN035, or KN035-TROP2^Nb^. (**E** and **F**) Western blot analysis (**E**) and quantification (**F**) of HER2 levels in SKBR3 cells treated for 24 h with 100 nM of 11A4, TROP2^Nb^, IGF2-11A4, or 11A4-TROP2^Nb^. (**G** and **H**) Flow cytometry analysis (**G**) and quantification (**H**) of PD-L1 levels in MDA-MB-231 cells treated for 24 h with 100 nM of KN035, TROP2^Nb^, IGF2-KN035, or KN035-TROP2^Nb^. (**I** and **J**) Flow cytometry analysis (**I**) and quantification (**J**) of HER2 levels in SKBR3 cells treated for 24 h with 100 nM of 11A4, TROP2^Nb^, IGF2-11A4, or 11A4-TROP2^Nb^. (**K** and **L**) Flow cytometry analysis (**K**) and quantification (**L**) of PD-L1 levels in MDA-MB-231 cells treated for 24 h with different concentrations of KN035-TROP2^Nb^. (**M** and **N**) Flow cytometry analysis (**M**) and quantification (**N**) of HER2 levels in SKBR3 cells treated for 24 h with different concentrations of 11A4-TROP2^Nb^. (**O**) MTT assay of SKBR3 cells treated with serial concentrations of 11A4, TROP2^Nb^, IGF2-11A4, or 11A4-TROP2^Nb^ for 7 d. Data are presented as mean ± SD (n = 3 biologically independent replicates) when relevant. *P* values were determined by one-way ANOVA with Turkey’s *post hoc* test. \*\**P* < 0.01; \*\*\**P* < 0.001; \*\*\*\**P* < 0.0001.

#### Construction and evaluation of TRTAC-DCs

Given the remarkable tumor-selectivity of TRTACs to mediate internalization and lysosomal delivery, we further engineered TRTACs into TRTAC-DCs (Figure 5A). For payload conjugation, we introduced a terminal cysteine handle into TRTAC sequences via a GSG linker. Reduction with tris(2-carboxyethyl)phosphine (TCEP) exposes the reactive thiol, which is then coupled with maleimide-functionalized valine-citrulline-linked monomethyl auristatin E (mal-vcMMAE) to yield TRTAC-DCs (Figure S5A) ^[48]^. We first synthesized a HER2-targeted TRTAC-DC (11A4-TROP2^Nb^-MMAE). MTT assays showed that 11A4-TROP2^Nb^-MMAE decreased SKBR3 cell viability by 50% at a concentration as low as 0.2 nM, whereas 11A4-MMAE and TROP2^Nb^-MMAE showed no significant activity at the same concentration, indicating that the potent cell -killing effect depends on the chimeric architecture (Figure 5B). In SKBR3 cell spheroids, 11A4-TROP2^Nb^-MMAE also exhibited stronger cytotoxicity compared to the non-chimeric controls (Figure 5C). Propidium iodide/Annexin V dual staining confirmed that 11A4-TROP2^Nb^-MMAE induced apoptosis in SKBR3 cells much more effectively than controls, in a dose-dependent manner (Figure 5D-G). Similar enhanced cytotoxicity was reproduced in HER2-overexpressing NCI-N87 cells (Figure 5H). In HER2-low HeLa cells and MRC-5 cells, however, moderate cytotoxicity was observed only at concentrations beyond 200 nM (Figure 5I and Figure S5B). No obvious cytotoxicity was detected in HUVECs even at 400 nM (Figure 5J). These findings underscore the favorable cell selectivity of 11A4-TROP2^Nb^-MMAE and its low toxicity toward normal cells.

**Figure 5.**
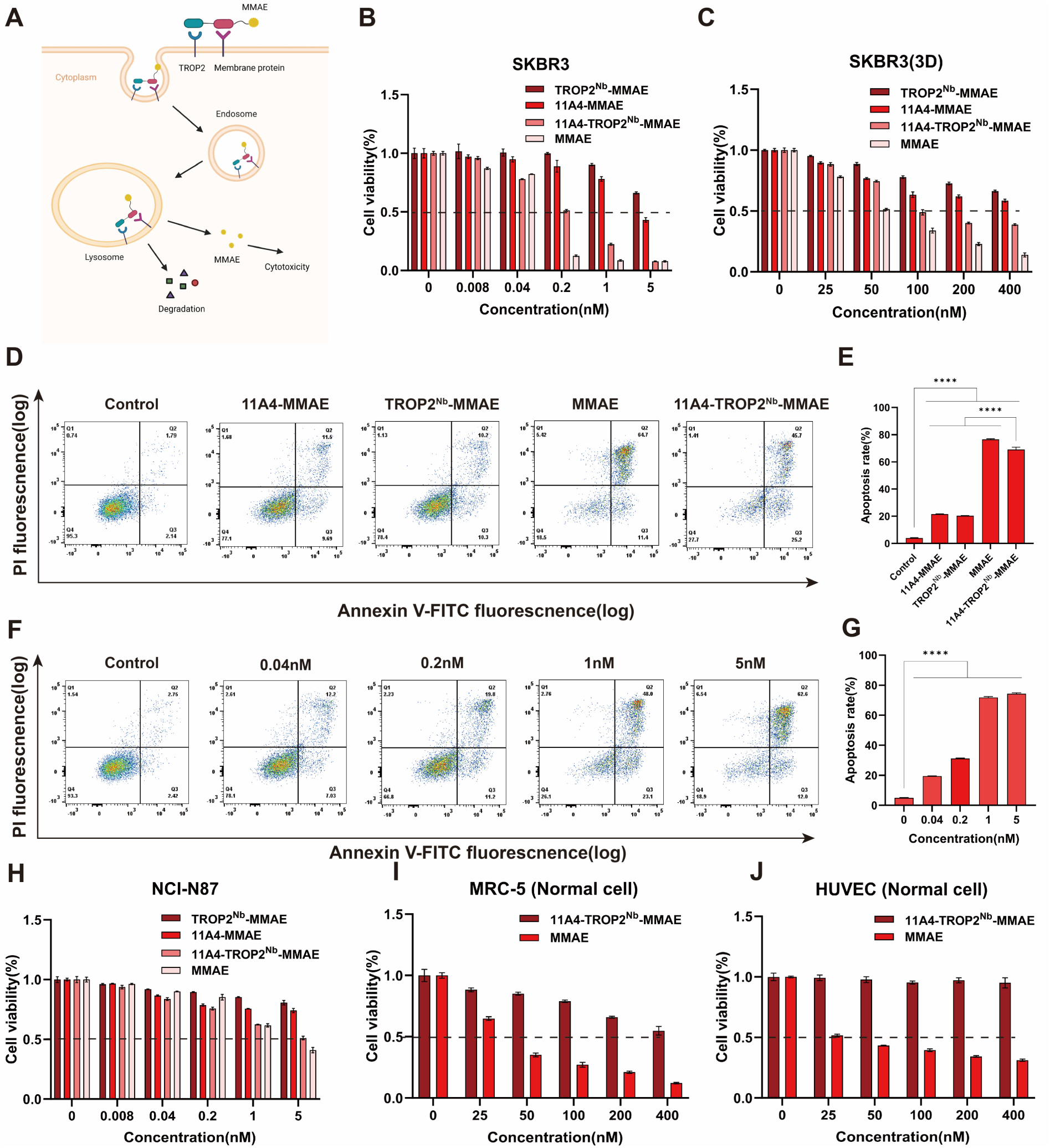
Construction of TRTAC-DC for HER2 degradation and enhanced drug delivery. (**A**) Schematic illustration of TRTAC-DC design for enhanced cytotoxin delivery while simultaneously inducing targeted membrane protein degradation. (**B**) MTT assay of SKBR3 cells treated with serial concentrations of 11A4-TROP2^Nb^-MMAE, 11A4-MMAE, TROP2^Nb^-MMAE, or free MMAE for 72 h. (**C**) Cell viability of SKBR3 tumor cell spheroids treated with serial concentrations of 11A4-TROP2^Nb^-MMAE, 11A4-MMAE, TROP2^Nb^-MMAE, or free MMAE for 72 h. (**D** and **E**) Flow cytometry analysis (**D**) and quantification (**E**) of apoptosis in SKBR3 cells after treatment with 1 nM of 11A4-TROP2^Nb^-MMAE, 11A4-MMAE, TROP2^Nb^-MMAE, or free MMAE for 48 h. (**F** and **G**) Flow cytometry analysis (**F**) and quantification (**G**) of apoptosis in SKBR3 cells after treatment with serial concentrations of 11A4-TROP2^Nb^-MMAE for 48 h. (**H-J**) MTT assay of NCI-N87 cells (**H**), MRC-5 cells (**I**), and HUVEC cells (**J**) treated with serial concentrations of 11A4-TROP2^Nb^-MMAE, 11A4-MMAE, TROP2^Nb^-MMAE, or free MMAE for 72 h. Data are presented as mean ± SD (n = 3 biologically independent replicates) when relevant. *P* values were determined by one-way ANOVA with Turkey’s *post hoc* test. \*\*\*\**P* < 0.0001.

Apart from the HER2-targeted TRTAC-DC, we next investigated the efficacy of the EGFR-targeted TRTAC-DC (7D12-TROP2^Nb^-MMAE). Cell viability assays showed that 7D12-TROP2^Nb^-MMAE killed HeLa cells more effectively than TROP2^Nb^-MMAE and 7D12-MMAE controls (Figure S5C). Moreover, 7D12-TROP2^Nb^-MMAE was highly cytotoxic to EGFR-high A431, MDA-MB-231, and SKBR3 cells while sparing HUVECs (Figure S5D-G), further substantiating the tumor cell selectivity of TRTAC-DCs. Previous studies have suggested that targeted EGFR degradation may help overcome resistance to tyrosine kinase inhibitors ^[12]^. We established a gefitinib-resistant PC-9/GR strain from gefitinib-sensitive PC-9 cells by stepwise *in vitro* selection with gefitinib (Figure S5H). The established PC-9/GR cells showed multi-drug resistance not only to gefitinib but also to MMAE (Figure S5I,J). At high concentrations, 7D12-TROP2^Nb^-MMAE was less cytotoxic to PC-9/GR cells than to parental PC-9 cells (Figure S5I,J). However, at 12.5 and 25 nM, 7D12-TROP2^Nb^-MMAE showed relatively greater cytotoxicity toward PC-9/GR cells, implying that EGFR degradation may contribute to overcoming resistance.

In addition to MMAE, we also explored the delivery of other cytotoxic payloads. A HER2-targeted TRTAC-doxorubicin conjugate (11A4-TROP2^Nb^-DOX) was synthesized similarly (Figure S5K) ^[49]^. This conjugate exhibited stronger cytotoxicity against SKBR3 cells than the non-chimeric counterparts and even free DOX (Figure S5L). In Hela cells, 11A4-TROP2^Nb^-DOX showed weaker activity with no advantage over free DOX (Figure S5M). Collectively, these findings demonstrate that TRTAC-DC displays promising activity *in vitro* and warrants further investigation as a potential antitumor therapeutic modality.

#### *In vivo* antitumor activity of TRTAC-DCs

Lastly, we assessed the *in vivo* antitumor effect of TRTAC-DCs. The HER2-targeted conjugate 11A4-TROP2^Nb^-MMAE was tested in mice bearing NCI-N87 xenografts. Mice were treated every two days with PBS, or 1 mg kg^-1^ of 11A4-MMAE, TROP2^Nb^-MMAE, or 11A4-TROP2^Nb^-MMAE via intravenous injection (Figure 6A). Tumor growth was more significantly suppressed with 11A4-TROP2^Nb^-MMAE treatment compared to the other treatments (Figure 6B-D). TUNEL (terminal deoxynucleotidyl transferase dUTP nick end labeling) staining revealed abundant apoptotic cells following 11A4-TROP2^Nb^-MMAE treatment, confirming the higher antitumor activity of 11A4-TROP2^Nb^-MMAE relative to 11A4-MMAE or TROP2^Nb^-MMAE (Figure 6E). Immunohistochemical staining showed that 11A4-TROP2^Nb^-MMAE markedly reduced HER2 expression in tumor tissues (Figure 6E). In addition, no significant body weight loss was observed throughout the experiment (Figure 6F). H&E staining of major organs showed no obvious abnormalities compared with controls, indicating a favorable *in vivo* safety profile (Figure S6). Similarly, we next evaluated the efficacy of the EGFR-targeted TRTAC-DC in HeLa tumor-bearing mice. Mice were intravenously administered PBS, or 2.5 mg kg^-1^ of 7D12-MMAE, TROP2^Nb^-MMAE, or 7D12-TROP2^Nb^-MMAE every two days (Figure 6G). Consistent with the results for 11A4-TROP2^Nb^-MMAE, this TRTAC-DC also exhibited superior antitumor activity than control treatments (Figure 6H-K), with no obvious systemic toxicity observed (Figure 6L). Together, these results confirm that TRTAC-DCs, which degrades cell-surface target proteins while promoting intracellular drug delivery, have promising safety and antitumor efficacy in an *in vivo* setting.

**Figure 6.**
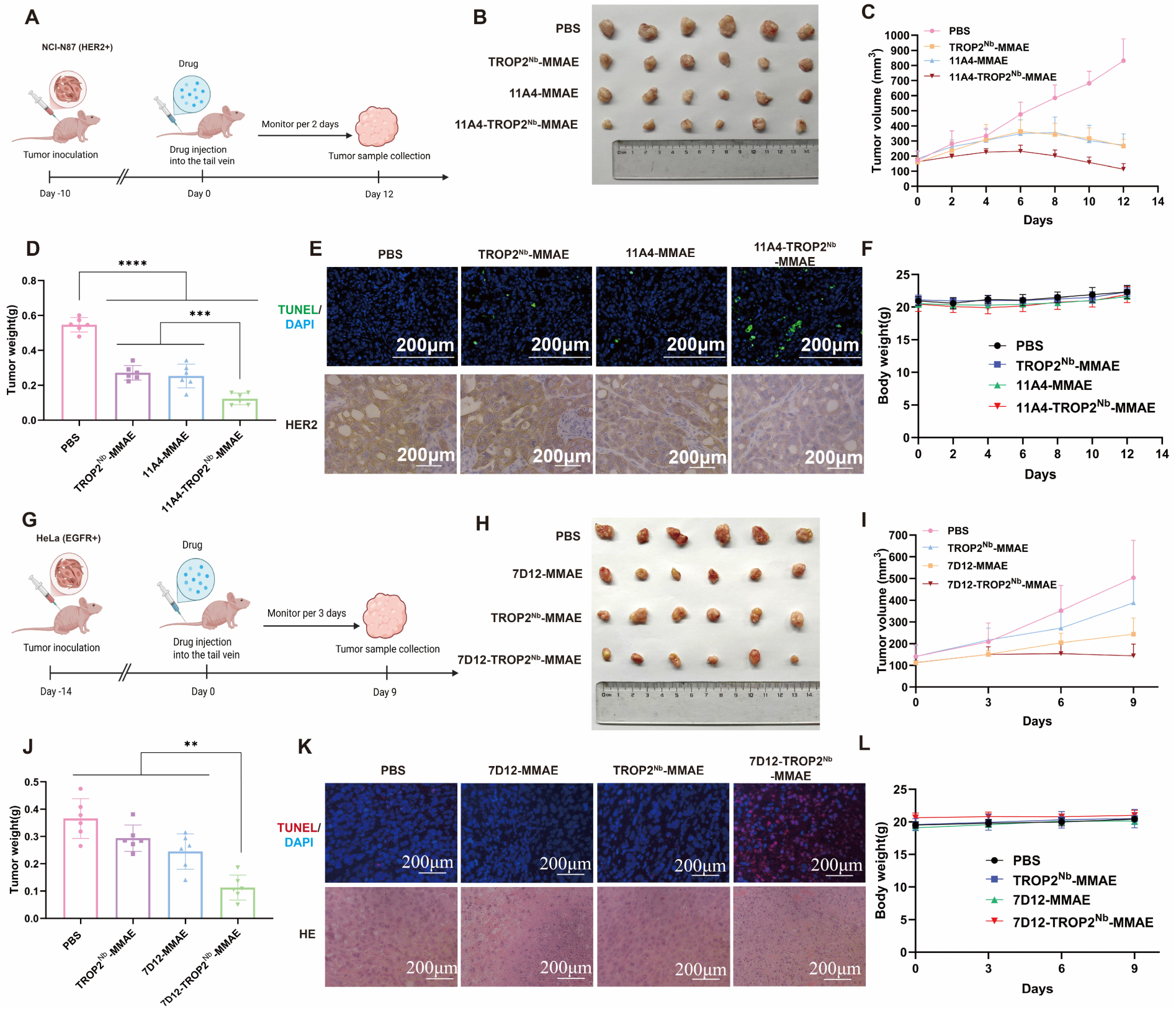
***In vivo* antitumor activity of TRTAC-DCs.** (**A-F**) Therapeutic efficacy of 11A4-TROP2^Nb^-MMAE in NCI-N87 xenografts. (**A**) Schematic illustrating the experimental design to assess the antitumor efficacy of 11A4-TROP2^Nb^-MMAE. NCI-N87 tumor-bearing nude mice were *i.v.* administered PBS, 11A4-MMAE, TROP2^Nb^-MMAE, or 11A4-TROP2^Nb^-MMAE at a 11A4-equiv. dose of 1 mg/kg every 2 days for 7 doses. (**B**) Photograph of excised tumors from each group. (**C**) Tumor growth curves in each group. (**D**) Average tumor weight of each group measured at the endpoint. (**E**) TUNEL staining and HER2 immunostaining of tumor sections collected from each group. Scale bar: 200 μm. (**F**) Body weight change of mice in each group. (**G-L**) Therapeutic efficacy of 7D12-TROP^Nb^-MMAE in HeLa xenografts. (**G**) Schematic illustrating the experimental design to assess the antitumor efficacy of 7D12-TROP^Nb^-MMAE. HeLa tumor-bearing nude mice were *i.v.* administered PBS, 7D12-MMAE, TROP2^Nb^-MMAE, or 7D12-TROP^Nb^-MMAE at a 7D12-equiv. dose of 2.5 mg/kg every 2 days for 5 doses. (**H**) Photograph of excised tumors from each group. (**I**) Tumor growth curves in each group. (**J**) Average tumor weight of each group measured at the endpoint. (**K**) TUNEL and H&E staining of tumor sections collected from each group. Scale bar: 200 μm. (**L**) Body weight change of mice in each group. Data are presented as mean ± SD (n = 6 mice per group). *P* values were determined by one-way ANOVA with Turkey’s *post hoc* test. \*\**P* < 0.01; \*\*\**P* < 0.001; \*\*\*\**P* < 0.0001.

## 3. Conclusion

This study has identified TROP2 from clinically validated ADC targets as an efficient tumor-selective LTR and established the TRTAC platform for targeted degradation of membrane and extracellular proteins. We show that TRTACs efficiently degraded oncogenic proteins including EGFR, HER2, and PD-L1, outperforming IGF2-based benchmark controls, in a tumor-selective manner. We further developed the TRTAC-DC format, which integrates eTPD with cytotoxic drug delivery. This bifunctional design exhibited potent antitumor effects in mouse xenograft models at doses where the corresponding nonchimeric conjugates showed only limited activity. By harnessing the efficient endocytosis and tumor-enriched expression pattern of TROP2, the TRTAC platform provides a versatile and translationally relevant strategy for converting existing binding modules into degraders and for enhancing the efficacy of conventional cytotoxic payloads.

These findings align with contemporary efforts to expand the scope of lysosomal targeting and intracellular delivery. Several studies have explored alternative LTRs with different degrees of tissue specificity ^[2, 9]^. Our results, together with previous reports, suggest that TROP2 trafficking to lysosomes is not merely constitutive but may be more effectively engaged in malignant cells, potentially because of altered membrane dynamics or endosomal sorting ^[50]^. This observation points to a broader opportunity for platform expansion. Replacing nonspecific internalization motifs with a TROP2-targeting module may enable the development of tumor-selective delivery systems for neoantigens or immune adjuvants. Such an approach may help convert tumors into *in situ* vaccines and induce localized antitumor immune responses while limiting toxicity in normal tissues. More broadly, the modular nature of the TRTAC scaffold should allow the incorporation of a wider range of payloads beyond tubulin inhibitors or anthracyclines. For instance, this platform may also be adapted for the delivery of PROTAC-like cargos, potentially enabling coordinated targeting of extracellular and intracellular disease associated proteins.

A further point of significant interest is the potential link between TROP2 and therapeutic resistance. Accumulating evidence suggests that TROP2 may contribute to the development of drug resistance, and that this effect can be at least partly alleviated by TROP2 inhibition ^[51, 52]^. In parallel, chemotherapy itself may increase TROP2 expression in certain settings. For example, tamoxifen treatment has been reported to markedly increase TROP2 expression in luminal breast cancer cell lines ^[53]^. TROP2 expression has also been associated with stemness related features in prostate cancer ^[54]^. In this context, the clinical success of anti-TROP2/SN-38 conjugates in patients who failed conventional chemotherapy may represent one of the clearest proofs of concept that TROP2 is functionally linked to chemoresistance. These observations raise an interesting question for future investigation that TRTAC and TRTAC-DC systems may not only serve as tumor-selective degradation and delivery platforms, but may also offer a strategy to overcome resistance in refractory tumors. This possibility is especially worth exploring in disease settings in which TROP2 is upregulated after treatment or enriched in drug tolerant cell populations.

Despite the promising therapeutic activity observed in this study, several limitations should be acknowledged. First, the use of small nanobody fusion proteins may improve tumor penetration relative to full length antibodies, but it also leads to rapid renal clearance and a short serum half-life ^[55]^. Further optimization will therefore require half-life extension strategies, such as fusion to albumin binding domains or other pharmacokinetic enhancement modules, to improve systemic exposure and reduce dosing frequency. Second, although TROP2 is broadly overexpressed in many epithelial cancers, its expression is not universal. Certain tumor types, including subsets of colorectal cancer and renal cell carcinoma, may express relatively low levels of TROP2, which could limit the applicability of this receptor without appropriate patient stratification. Third, the *in vivo* validation presented here was mainly based on conventional cell line derived xenograft models. Although these models clearly support target engagement and therapeutic efficacy, they do not fully capture the stromal complexity, immune context, or intratumoral heterogeneity of human tumors. Future studies in patient derived xenografts, organoid based systems, and more physiologically relevant tumor models will be important for evaluating clinical response and for clarifying how TROP2 heterogeneity influences therapeutic outcome.

## 4. Experimental Section

*Reagents and antibodies:* Kanamycin, 3-[4,5-dimethylthiazol-2-yl]-2,5 diphenyl tetrazolium bromide (MTT), RIPA lysis buffer, protease/phosphatase inhibitor cocktail, chlorpromazine, bafilomycin A1, LysoTracker Red, recombinant human TROP2 protein (HY-P70457), maleimide-functionalized valine-citrulline-linked monomethyl auristatin E (mal-vcMMAE; HY-15575), (6-maleimidocaproyl)hydrazone doxorubicin (DOXO-EMCH; HY-16261), and tris(2-carboxyethyl)phosphine (TCEP) were purchased from MedChemExpress. Antibodies used for immunoblotting included anti-EGFR (1:5000, Proteintech, 66455-1-Ig), anti-HER2 (1:5000, Proteintech, 18299-1-AP), anti-GAPDH (1:5000, Proteintech, 10494-1-AP), anti-PD-L1 (1:1000, Cell Signaling Technology, 13684T), HRP-conjugated goat anti-rabbit IgG (1:5000, Proteintech, SA00001-2), and goat anti-mouse IgG (1:5000, Proteintech, SA00001-1). Antibodies used for flow cytometry included APC-conjugated anti-EGFR (1:40, BioLegend, 352906), APC-conjugated anti-PD-L1 (1:40, BioLegend, 329708), and APC-conjugated anti-HER2 (1:40, BioLegend, 324408). HRP-conjugated anti-GFP nanobody (1:4000, AlpalifeBio, KTSM1311) was used for ELISA.

*Cell lines:* HeLa, MDA-MB-231, SKBR3, NCI-N87, BXPC-3, H1975, MRC-5, HUVEC, and A431 cells were obtained from the Institute of Biochemistry and Cell Biology, Chinese Academy of Sciences. MDA-MB-231, SKBR3, BXPC-3, H1975, MRC-5, and A431 cells were cultured in DMEM (Corning). HeLa, NCI-N87, and HUVEC cells were cultured in RPMI-1640 (Corning). All media were supplemented with 10% Fetal bovine serum (FBS) (LONSERA) and 1% penicillin-streptomycin (Coolaber). Cells were maintained at 37 °C in a humidified incubator with 5% CO_2_.

*Tumor cell spheroid formation:* HeLa, H1975, or SKBR3 cells were seeded into ultra-low attachment plates and cultured at 37 °C for 2–4 days until reaching a relatively stable size. Spheroids were treated as indicated.

*Plasmid construction:* Genes encoding 11A4, CD44^Nb^, CLDN18.2^Nb^, Frα^binder^, TF^binder^, TROP2^Nb^, TROP2^Nb108^, TROP2^NbV-5^, and IGF2 were synthesized by Tsingke Biotechnology. Fusion protein constructs were generated by *in vitro* recombination using laboratory stock vectors, including 7D12-LZ-8-pET28b, EGFP-LZ8-pET28b and EGFP-pET28a. Plasmids carrying inserts and corresponding vector backbones were first amplified in LB medium containing kanamycin and purified for use as templates. Target genes were amplified by PCR, recovered from agarose gels, and subjected to double digestion with matched restriction enzymes together with the corresponding vectors. After purification, digested products were ligated with T4 DNA ligase at an insert/vector ratio of 10:1. Ligation products were transformed into competent *E. coli*. Single colonies were expanded and verified by Sanger sequencing. Sequence-confirmed clones were used for protein expression.

*Recombinant protein expression and purification:* Sequence-verified expression plasmids were transformed into *E. coli* BL21(DE3) competent cells. Single colonies were inoculated into LB medium containing kanamycin and grown overnight at 37 °C with shaking. Cultures were then transferred into fresh medium and grown to an OD_600_ of 0.6–0.8. Protein expression was induced with 0.5 mM IPTG at 16 °C for 15 h. Bacteria pellets were collected by centrifugation, resuspended in PBS, and disrupted by sonication. Lysates were centrifuged at 4 °C and the supernatants were collected for purification. All recombinant proteins were loaded onto a pre-equilibrated nickel column, washed sequentially with PBS, 20 mM imidazole, and 40 mM imidazole, and then eluted with 200 mM imidazole. Eluates were dialyzed against PBS, filtered through a 0.22-μm filter, aliquoted and stored at −80 °C. Protein concentrations were determined using the BCA Protein Assay Kit (Beyotime, P0012).

*Flow cytometry:* Cells were seeded in 24-well plates (1×10^6^ cells per well) one day before the experiments. For receptor screening, cells were incubated with 500 nM of 7D12, 7D12-CD44^Nb^, 7D12-TF^Binder^, 7D12-FRα^Binder^, 7D12-HER2^Nb^, 7D12-CLDN18.2^Nb^, or 7D12-TROP2^Nb^ for 24 h, treated with trypan blue to quench cell surface fluorescence, washed with PBS, and then analyzed using flow cytometry. For membrane protein detection, cells were incubated with specific antibodies at 4 °C for 60 min, washed with PBS, and then analyzed using flow cytometry. For cell apoptosis analysis, cells were treated as indicated, washed with PBS, stained with the Annexin V-FITC Apoptosis Detection Kit (Beyotime, C1062L) according to the manufacturer’s instructions, and analyzed using flow cytometry. Flow cytometry was performed using a Beckman Coulter CytoFLEX LX Flow Cytometer. Data were acquired with CytExpert v2.3.0.84 and analyzed with FlowJo v10.0.7.

*Confocal imaging for lysosomal colocalization analysis:* HeLa cells were seeded in 24-well plates (5–8×10^5^ cells per well). After 24 h of culture, cells were incubated with TROP2^Nb^-EGFP for 6 h, washed with PBS, stained with DAPI and LysoTracker Red for 30 min. Cells were washed with PBS and imaged using a Nikon A1R confocal microscope.

*Western blot analysis:* Protein samples were prepared by lysing cells in ice-cold RIPA buffer supplemented with 1× protease/phosphatase inhibitor cocktail. Lysates were centrifuged at 4 °C and supernatants were collected. Protein concentrations were determined by the BCA assay. Protein samples were mixed with loading buffer. For detection of EGFR or HER2, samples were denatured at 37 °C for 30 min; all other samples were denatured at 95 °C for 10 min. After cooling, samples were separated by 10% SDS–PAGE and transferred to 0.45 μm polyvinylidene fluoride (PVDF) membranes (Cytiva, 10600029). Blots were blocked with protein-free rapid blocking solution for 30 min at room temperature, incubated with primary antibodies overnight at 4 °C, washed three times with TBST for 10 min each, and then incubated with secondary antibodies for 50 min at room temperature. After three washes with TBST, blots were developed with BeyoECL Star (Beyotime) for 2 min and imaged using a Tanon 5200 Chemiluminescent Imaging System. Band intensity was quantified using ImageJ.

*Cell viability assays:* Cells were seeded in 96-well plates (1.0×10^4^ cells per well). After 3-7 d of culture, cells were treated as indicated. For 2D cell cultures, MTT was added to each well and cells were incubated at 37 °C for 2 h. Supernatants were then removed, formazan crystals were dissolved in DMSO, and absorbance at 570 nm was measured with a microplate reader. Cell viability was calculated as the relative OD_570_ value of treated cells compared with control cells. For 3D spheroids, cell viability was assessed using the CellTiter-Lumi™ Plus II Luminescent Cell Viability Assay Kit (Beyotime, C0057S) according to the manufacturer’s instructions.

*ELISA:* Recombinant human TROP2 protein was diluted in coating buffer (10 µg/mL) and added to ELISA plates at 100 μL per well. After incubation at 4 °C overnight, the coating solution was removed and plates were blocked with PBST containing 5% BSA for 1 h at room temperature. Serial concentrations of TROP2^Nb^-EGFP, TROP2^Nb108^-EGFP, TROP2^NbV-5^-EGFP, or EGFP were then added and incubated for 1 h at room temperature. After washing three times with PBST, HRP-conjugated anti-EGFP was added and incubated for 45min at room temperature. After washing five times with PBST, 150 μL of TMB substrate solution (Beyotime, P0209) was added, and reactions were stopped with 50 μL of 10% H_2_SO_4_. Absorbance was measured at 450 nm using a Thermo Fisher Scientific Labserv-K3 multiplate reader. Binding curves were fitted using GraphPad Prism and apparent EC_50_ values were calculated.

*Molecular docking and binding free energy analysis:* Structural models of TROP2, TROP2^Nb^, TROP2^Nb108^, and TROP2^NbV-5^ were obtained from the AlphaFold Protein Structure Database. Protein docking was carried out using the HDOCK server. Binding free energy was calculated using the MM/GBSA mode of the HawkDock platform. The best model was selected from the docking outputs according to docking score, and contributions of interface residues were further analyzed. Final structures were visualized with PyMOL 2.4.

*Preparation of TRTAC-DCs:* To construct TRTAC-DCs, a cysteine residue was introduced at the C-terminus of fusion proteins. For payload conjugation, proteins were first reduced with TCEP, followed by incubation with either mal-vcMMAE or DOXO-EMCH. Each treatment was performed at a 2:1 molar ratio in pH 6.5–7.5 buffer at 37 °C for 1 h. TRTAC-DCs were purified by dialysis against PBS for 12 h.

*Animal ethics statement:* Female BALB/c nude mice (6-week-old) were purchased from Shanghai SLAC Laboratory Animal Co., Ltd. and housed under specific pathogen-free conditions with free access to food and water under a 12-h light-dark cycle. All animal experiments were approved by the Bioethics Committee of East China University of Science and Technology under an approval number ECUST-2025-107 and were performed in accordance with relevant institutional guidelines for animal care and use. Animal suffering was minimized throughout the study, and experiments were terminated according to predefined humane endpoint criteria.

*In vivo antitumor experiments:* Tumor cells suspended in PBS were injected subcutaneously into the right dorsal forelimb region of mice at a density of 2×10^6^ cells per mouse. Tumors were allowed to grow for 2–3 weeks until the average tumor volume reached 150 mm^3^. Mice were randomly assigned to different groups and treated as indicated by tail vein injection. Body weight and tumor volume were monitored every 2–3 d. Tumor volume was calculated following the formula: tumor volume = Length (*L*) × Width^2^ (*W*^2^)/2. Mice were euthanized when tumors reached approximately 600–700 mm^3^. Tumors and major organs were excised for histopathological analysis.

*H&E staining, immunohistochemistry and TUNEL staining:* Excised tumor and organ tissues were fixed in 4% paraformaldehyde, embedded in paraffin, and sectioned using standard procedures. For histological analysis, sections were stained with hematoxylin and eosin (H&E) and examined under an optical microscope. For immunohistochemical staining, sections were deparaffinized, rehydrated, subjected to antigen retrieval, and then treated to block endogenous peroxidase activity and nonspecific binding. Sections were then incubated with primary antibodies at 4 °C overnight, washed five times with TBST, incubated with HRP-conjugated secondary antibodies, followed by TBST washes, DAB development, hematoxylin counterstaining, and mounting. Images were acquired using an optical microscopy. For detection of apoptotic cells, sections were analyzed using the TUNEL Cell Apoptosis Detection Kit (Servicebio, G1502) following the manufacturer’s instructions. Sections were deparaffinized, rehydrated, treated with Proteinase K (Servicebio, G1237) at 37 °C for 20 min, followed by incubation with Permeabilization Solution (Servicebio, G1204). Sections were then incubated with Reaction Solution containing recombinant TdT enzyme and dUTP in a humidified chamber at 37 °C for 1 h. After counterstaining with DAPI (Servicebio, G1012), sections were mounted with antifade mounting medium (Servicebio, G1401). Images were acquired using a NIKON ECLIPSE C1 fluorescence microscope.

*Statistical analysis:* All data are presented as mean ± SD. *In vitro* experiments were generally performed with three independent biological replicates, and animal experiments used a sample size of n = 5-6 per group. Statistical significance between two independent groups was determined using two-tailed, unpaired Student’s *t*-test. Multiple groups were compared using one-way ANOVA with Dunnett’s or Tukey’s *post hoc* test accordingly. *P* values of <0.05 were considered statistically significant. ns, no significance; \**P* < 0.05; \*\**P* < 0.01; \*\*\**P* < 0.001; \*\*\*\**P* < 0.0001. All statistical analyses were performed using GraphPad Prism 10.1.2.

## Acknowledgements

This work was financially supported by the National Natural Science Foundation of China (32121005, 32571667, 52473153, and 32327801), Natural Science Foundation of Shanghai (24ZR1417000), the Shanghai Excellent Young Academic Leader Program of Eastern Talent Plan (QNKJ2025047), the Zhejiang Provincial Natural Science Foundation of China (LR25E030002), Key Research and Development Program of Zhejiang Province (2025E10069), the Shanghai Sci-Tech Inno Center for Infection & Immunity (SSIII-2024A0302), the International Cooperation and Exchange of the National Natural Science Foundation of China (W2412094), the National Science and Technology Major Project of China (2023ZD0508000), the Fundamental Research Funds for the Central Universities (226-2025-00196), and Zhejiang Fonow Medicine Co., Ltd. (F100-42106).

## Data Availability Statement

All data needed to evaluate and reproduce the results in the paper are present in the paper and the Supporting Information.

## Funding

National Natural Science Foundation of China (32121005, 32571667, 52473153, 32327801) Natural Science Foundation of Shanghai (24ZR1417000)

Shanghai Excellent Young Academic Leader Program of Eastern Talent Plan (QNKJ2025047) Zhejiang Provincial Natural Science Foundation of China (LR25E030002)

Key Research and Development Program of Zhejiang Province (2025E10069) Shanghai Sci-Tech Inno Center for Infection & Immunity (SSIII-2024A0302) International Cooperation and Exchange Program of NSFC (W2412094)

National Science and Technology Major Project of China (2023ZD0508000) Fundamental Research Funds for the Central Universities (226-2025-00196) Zhejiang Fonow Medicine Co., Ltd. (F100-42106)

## Supporting Information

Supporting Information is available from the Wiley Online Library or from the author.

TROP2 is identified as a novel tumor-selective lysosomal-targeting receptor with established clinical relevance. TROP2-targeting chimeras (TRTACs) are developed by genetically fusing a TROP2-binding nanobody to nanobodies against specific target proteins. TRTACs can induce tumor-selective degradation of diverse membrane proteins and enhance drug delivery.

TROP2-targeting chimeras (TRTACs) for tumor-selective membrane protein degradation and enhanced drug delivery

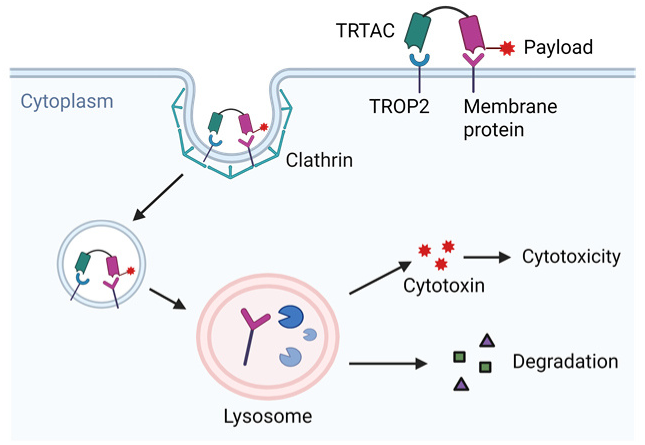

