## Supplementary Material for "TROP2-targeting chimeras (TRTACs) for tumor-selective membrane protein degradation and enhanced drug delivery"

F. Wang
New Drug R&D Center, Zhejiang Fonow Medicine Co. Ltd., Dongyang, Zhejiang 322100, China

F. Wang
*Institute of Chinese Materia Medica, Shanghai University of Traditional Chinese Medicine, Shanghai, 201203, China*

**Supporting Information text**

Protein sequence

11A4

MGEVQLVESGGGLVQAGGSLRLSCATSGITFMRYALGWYRQSPGKQREMVASINSG
GTTNYADSVKGRFTISRDNAKNTVYLQMNSLKPEDTAVYYCNARWVKPQFIDNNY
WGQGTQVTVSSLELEHHHHHH

11A4-cys

MGSSHHHHHHSSGLVPRGSHMEVQLVESGGGLVQAGGSLRLSCATSGITFMRYALG
WYRQSPGKQREMVASINSGGTTNYADSVKGRFTISRDNAKNTVYLQMNSLKPEDTA
VYYCNARWVKPQFIDNNYWGQGTQVTVSSLEGSGC

7D12-cys

MGSSHHHHHHSSGLVPRGSHMQVKLEESGGGSVQTGGSLRLTCAASGRTSRSYGMG
WFRQAPGKEREFVSGISWRGDSTGYADSVKGRFTISRDNAKNTVDLQMNSLKPEDT
AIYYCAAAAGSAWYGTLYEYDYWGQGTQVTVSSGSGC

7D12-11A4

MAMQVKLEESGGGSVQTGGSLRLTCAASGRTSRSYGMGWFRQAPGKEREFVSGISW
RGDSTGYADSVKGRFTISRDNAKNTVDLQMNSLKPEDTAIYYCAAAAGSAWYGTLY
EYDYWGQGTQVTVSSGSGGGGSGGGGSEVQLVESGGGLVQAGGSLRLSCATSGITF
MRYALGWYRQSPGKQREMVASINSGGTTNYADSVKGRFTISRDNAKNTVYLQMNS
LKPEDTAVYYCNARWVKPQFIDNNYWGQGTQVTVSSLELEHHHHHH

RLM(TF)-EGFP

MGRLMTQDCLQQSRKGGGGSGGGGSGGGGSRLMTQDCLQQSRKGGGGSGGGGSG
GGGSRLMTQDCLQQSRKGGGGSGGGGSGSGGGGSVSKGEELFTGVVPILVELDGDV
NGHKFSVSGEGEGDATYGKLTLKFICTTGKLPVPWPTLVTTLTYGVQCFSRYPDHMK
QHDFFKSAMPEGYVQERTIFFKDDGNYKTRAEVKFEGDTLVNRIELKGIDFKEDGNIL
GHKLEYNYNSHNVYIMADKQKNGIKVNFKIRHNIEDGSVQLADHYQQNTPIGDGPV
LLPDNHYLSTQSALSKDPNEKRDHMVLLEFVTAAGITLGMDELYKLEHHHHHH

Trop2$^{Nb}$-EGFP

MGQVQLQESGGGLVQPGGSLRLSCAASGFTLDSYAIAWFRQAPGKEREGVSCIRSKD GSTYYADSVKGRFTISRDNAKNTVYLQMNSLKPEDTAVYYCAACDDDDEGGIIKMR SSSTVVWYYYYWGKGTQVTVSSGSGGGGSVSKGEELFTGVVPILVELDGDVNGHKF SVSGEGEGDATYGKLTLKFICTTGKLPVPWPTLVTTLTYGVQCFSRYPDHMKQHDFF KSAMPEGYVQERTIFFKDDGNYKTRAEVKFEGDTLVNRIELKGIDFKEDGNILGHKLE YNYNSHNVYIMADKQKNGIKVNFKIRHNIEDGSVQLADHYQQNTPIGDGPVLLPDNH YLSTQSALSKDPNEKRDHMVLLEFVTAAGITLGMDELYKLEHHHHHH

$EGFP^{Nb}$-$Trop2^{Nb}$

MGQVQLVESGGALVQPGGSLRLSCAASGFPVNRYSMRWYRQADTNNDGWIEGDEL KEREWVAGMSSAGDRSSYEDSVKGRFTISRDDARNTVYLQMNSLKPEDTAVYYCNV NVGFEYWGQGTQVTVSSGSGGGGSGGGGSGGGGSQVQLQESGGGLVQPGGSLRLS CAASGFTLDSYAIAWFRQAPGKEREGVSCIRSKDGSTYYADSVKGRFTISRDNAKNT VYLQMNSLKPEDTAVYYCAACDDDDEGGIIKMRSSSTVVWYYYYWGKGTQVTVSS LEHHHHHH

11A4-EGFP

MGEVQLVESGGGLVQAGGSLRLSCATSGITFMRYALGWYRQSPGKQREMVASINSG GTTNYADSVKGRFTISRDNAKNTVYLQMNSLKPEDTAVYYCNARWVKPQFIDNNY WGQGTQVTVSSLEGGGGSGGGGSGSGGGGSVSKGEELFTGVVPILVELDGDVNGHK FSVSGEGEGDATYGKLTLKFICTTGKLPVPWPTLVTTLTYGVQCFSRYPDHMKQHDF FKSAMPEGYVQERTIFFKDDGNYKTRAEVKFEGDTLVNRIELKGIDFKEDGNILGHKL EYNYNSHNVYIMADKQKNGIKVNFKIRHNIEDGSVQLADHYQQNTPIGDGPVLLPDN HYLSTQSALSKDPNEKRDHMVLLEFVTAAGITLGMDELYKLEHHHHHH

7D12-RLM(TF)

MAMQVKLEESGGGSVQTGGSLRLTCAASGRTSRSYGMGWFRQAPGKEREFVSGISW RGDSTGYADSVKGRFTISRDNAKNTVDLQMNSLKPEDTAIYYCAAAAGSAWYGTLY EYDYWGQGTQVTVSSGSGGGGSGGGGSRLMTQDCLQQSRKGGGGSGGGGSGGGGS RLMTQDCLQQSRKGGGGSGGGGSGGGGSRLMTQDCLQQSRKLEHHHHHH

11A4-$Trop2^{Nb}$-cys

MGSSHHHHHHSSGLVPRGSHMEVQLVESGGGLVQAGGSLRLSCATSGITFMRYALG WYRQSPGKQREMVASINSGGTTNYADSVKGRFTISRDNAKNTVYLQMNSLKPEDTA

VYYCNARWVKPQFIDNNYWGQGTQVTVSSLEGSGGGGSGGGGSQVQLQESGGGLV
QPGGSLRLSCAASGFTLDSYAIAWFRQAPGKEREGVSCIRSKDGSTYYADSVKGRFTI
SRDNAKNTVYLQMNSLKPEDTAVYYCAACDDDDEGGIIKMRSSSTVVWYYYYWGK
GTQVTVSSGSGC

Trop2$^{Nb108}$-EGFP

MGQVQLQESGGGLVQPGGSLRLSCAASGRIFSSFAMAWFRQAPGKEREFVTAITWSG
GSTYYADSVKGRFTISRDNAKNTVYLQMNSLKPEDTAVYYCAAADDEGLSTVVYYY
WGQGTQVTVSSGGGGSGGGGSGSGGGGSVSKGEELFTGVVPILVELDGDVNGHKFS
VSGEGEGDATYGKLTLKFICTTGKLPVPWPTLVTTLTYGVQCFSRYPDHMKQHDFFK
SAMPEGYVQERTIFFKDDGNYKTRAEVKFEGDTLVNRIELKGIDFKEDGNILGHKLEY
NYNSHNVYIMADKQKNGIKVNFKIRHNIEDGSVQLADHYQQNTPIGDGPVLLPDNHY
LSTQSALSKDPNEKRDHMVLLEFVTAAGITLGMDELYKLEHHHHHH

CLND18.2$^{Nb}$-EGFP

MGQVQLVESGGGLVQPGGSLRLSCAASGVDISSDVMAWYRQAPGKGLEFVSGLTRG
GSINYADSVKGRFTISRDNSKNTLYLQMNSLRAEDTAVYYCNAEIYTGTFYPRSYWG
QGTLVTVSGGGGSGGGGSGSGGGGSVSKGEELFTGVVPILVELDGDVNGHKFSVSGE
GEGDATYGKLTLKFICTTGKLPVPWPTLVTTLTYGVQCFSRYPDHMKQHDFFKSAMP
EGYVQERTIFFKDDGNYKTRAEVKFEGDTLVNRIELKGIDFKEDGNILGHKLEYNYNS
HNVYIMADKQKNGIKVNFKIRHNIEDGSVQLADHYQQNTPIGDGPVLLPDNHYLSTQ
SALSKDPNEKRDHMVLLEFVTAAGITLGMDELYKLEHHHHHH

C7(FRα)-EGFP

MGMHTAPGWGYRLSGGGGSGGGGSGGGGSMHTAPGWGYRLSGGGGSGGGGSGG
GGSMHTAPGWGYRLSGGGGSGGGGSGSGGGGSVSKGEELFTGVVPILVELDGDVNG
HKFSVSGEGEGDATYGKLTLKFICTTGKLPVPWPTLVTTLTYGVQCFSRYPDHMKQH
DFFKSAMPEGYVQERTIFFKDDGNYKTRAEVKFEGDTLVNRIELKGIDFKEDGNILGH
KLEYNYNSHNVYIMADKQKNGIKVNFKIRHNIEDGSVQLADHYQQNTPIGDGPVLLP
DNHYLSTQSALSKDPNEKRDHMVLLEFVTAAGITLGMDELYKLEHHHHHH

CD44$^{Nb}$-EGFP

MGMQVQLVESGGGSVQAGGSLRLSCTASGGSEYSYSTFSLGWFRQAPGQEREAVAA
IASMGGLTYYADSVKGRFTISRDNAKNTVTLQMNNLKPEDTAIYYCAARNAGGRFR

PSAAGGYNYWGQGTLVTVSSGGGGSGGGGSGSGGGGSVSKGEELFTGVVPILVELD
GDVNGHKFSVSGEGEGDATYGKLTLKFICTTGKLPVPWPTLVTTLTYGVQCFSRYPD
HMKQHDFFKSAMPEGYVQERTIFFKDDGNYKTRAEVKFEGDTLVNRIELKGIDFKED
GNILGHKLEYNYNSHNVYIMADKQKNGIKVNFKIRHNIEDGSVQLADHYQQNTPIGD
GPVLLPDNHYLSTQSALSKDPNEKRDHMVLLEFVTAAGITLGMDELYKLEHHHHHH

IGF2-11A4

MGLCGGELVDTLQFVCGDRGFYFSRPASRVSRRSRGIVEECCFRSCDLALLETYCATP
AKSEEFGSGGGGSGGGGSEVQLVESGGGLVQAGGSLRLSCATSGITFMRYALGWYR
QSPGKQREMVASINSGGTTNYADSVKGRFTISRDNAKNTVYLQMNSLKPEDTAVYY
CNARWVKPQFIDNNYWGQGTQVTVSSLELEHHHHHH

7D12-Trop2$^{Nb108}$

MAMQVKLEESGGGSVQTGGSLRLTCAASGRTSRSYGMGWFRQAPGKEREFVSGISW
RGDSTGYADSVKGRFTISRDNAKNTVDLQMNSLKPEDTAIYYCAAAAGSAWYGTLY
EYDYWGQGTQVTVSSGSGGGGSGGGGSQVQLQESGGGLVQPGGSLRLSCAASGRIF
SSFAMAWFRQAPGKEREFVTAITWSGGSTYYADSVKGRFTISRDNAKNTVYLQMNS
LKPEDTAVYYCAAADDEGLSTVVYYYWGQGTQVTVSSLEHHHHHH

7D12-CD44$^{Nb}$

MAMQVKLEESGGGSVQTGGSLRLTCAASGRTSRSYGMGWFRQAPGKEREFVSGISW
RGDSTGYADSVKGRFTISRDNAKNTVDLQMNSLKPEDTAIYYCAAAAGSAWYGTLY
EYDYWGQGTQVTVSSGSGGGGSGGGGSMQVQLVESGGGSVQAGGSLRLSCTASGG
SEYSYSTFSLGWFRQAPGQEREAVAAIASMGGLTYYADSVKGRFTISRDNAKNTVTL
QMNNLKPEDTAIYYCAARNAGGRFRPSAAGGYNYWGQGTLVTVSSLEHHHHHH

Trop2$^{Nb}$-cys

MGSSHHHHHHSSGLVPRGSHMQVQLQESGGGLVQPGGSLRLSCAASGFTLDSYAIA
WFRQAPGKEREGVSCIRSKDGSTYYADSVKGRFTISRDNAKNTVYLQMNSLKPEDTA
VYYCAACDDDDEGGIIKMRSSSTVVWYYYYWGKGTQVTVSSGSGC

11A4-Trop2$^{Nb}$

MGEVQLVESGGGLVQAGGSLRLSCATSGITFMRYALGWYRQSPGKQREMVASINSG
GTTNYADSVKGRFTISRDNAKNTVYLQMNSLKPEDTAVYYCNARWVKPQFIDNNY

WGQGTQVTVSSLEGSGGGGSGGGGSGGGGSQVQLQESGGGLVQPGGSLRLSCAASGFTLDSYAIAWFRQAPGKEREGVSCIRSKDGSTYYADSVKGRFTISRDNAKNTVYLQMNSLKPEDTAVYYCAACDDDDEGGIIKMRSSSTVVWYYYYWGKGTQVTVSSLEHHHHHH

7D12-Trop2$^{Nb}$-cys

MGSSHHHHHHSSGLVPRGSHMQVKLEESGGGSVQTGGSLRLTCAASGRTSRSYGMGWFRQAPGKEREFVSGISWRGDSTGYADSVKGRFTISRDNAKNTVDLQMNSLKPEDTAIYYCAAAAGSAWYGTLYEYDYWGQGTQVTVSSGSGGGGSGGGGSQVQLQESGGGLVQPGGSLRLSCAASGFTLDSYAIAWFRQAPGKEREGVSCIRSKDGSTYYADSVKGRFTISRDNAKNTVYLQMNSLKPEDTAVYYCAACDDDDEGGIIKMRSSSTVVWYYYYWGKGTQVTVSSGSGC

7D12-Trop2$^{Nb}$

MAMQVKLEESGGGSVQTGGSLRLTCAASGRTSRSYGMGWFRQAPGKEREFVSGISWRGDSTGYADSVKGRFTISRDNAKNTVDLQMNSLKPEDTAIYYCAAAAGSAWYGTLYEYDYWGQGTQVTVSSGSGGGGSGGGGSQVQLQESGGGLVQPGGSLRLSCAASGFTLDSYAIAWFRQAPGKEREGVSCIRSKDGSTYYADSVKGRFTISRDNAKNTVYLQMNSLKPEDTAVYYCAACDDDDEGGIIKMRSSSTVVWYYYYWGKGTQVTVSSLEHHHHHH

KN035-Trop2$^{Nb}$

MGTGQVQLQESGGGLVQPGGSLRLSCAASGKMSSRRCMAWFRQAPGKERERVAKLLTTSGSTYLADSVKGRFTISQNNAKSTVYLQMNSLKPEDTAMYYCAADSFEDPTCTLVTSSGAFQYWGQGTQVTVSSGSMDPGGSGSGGGGSGGGGSQVQLQESGGGLVQPGGSLRLSCAASGFTLDSYAIAWFRQAPGKEREGVSCIRSKDGSTYYADSVKGRFTISRDNAKNTVYLQMNSLKPEDTAVYYCAACDDDDEGGIIKMRSSSTVVWYYYYWGKGTQVTVSSLEHHHHHH

7D12-CLDN18.2$^{Nb}$

MAMQVKLEESGGGSVQTGGSLRLTCAASGRTSRSYGMGWFRQAPGKEREFVSGISWRGDSTGYADSVKGRFTISRDNAKNTVDLQMNSLKPEDTAIYYCAAAAGSAWYGTLYEYDYWGQGTQVTVSSGSGGGGSGGGGSQVQLVESGGGLVQPGGSLRLSCAASGVDISSDVMAWYRQAPGKGLEFVSGLTRGGSINYADSVKGRFTISRDNSKNTLYLQMNSLRAEDTAVYYCNAEIYTGTFYPRSYWGQGTLVTVSLEHHHHHH

7D12-EGFP

MAMQVKLEESGGGSVQTGGSLRLTCAASGRTSRSYGMGWFRQAPGKEREFVSGISW
RGDSTGYADSVKGRFTISRDNAKNTVDLQMNSLKPEDTAIYYCAAAAGSAWYGTLY
EYDYWGQGTQVTVSSGSGGGGSVSKGEELFTGVVPILVELDGDVNGHKFSVSGEGE
GDATYGKLTLKFICTTGKLPVPWPTLVTTLTYGVQCFSRYPDHMKQHDFFKSAMPEG
YVQERTIFFKDDGNYKTRAEVKFEGDTLVNRIELKGIDFKEDGNILGHKLEYNYNSH
NVYIMADKQKNGIKVNFKIRHNIEDGSVQLADHYQQNTPIGDGPVLLPDNHYLSTQS
ALSKDPNEKRDHMVLLEFVTAAGITLGMDELYKLEHHHHHH

7D12-CD44$^{binder}$

MAMQVKLEESGGGSVQTGGSLRLTCAASGRTSRSYGMGWFRQAPGKEREFVSGISW
RGDSTGYADSVKGRFTISRDNAKNTVDLQMNSLKPEDTAIYYCAAAAGSAWYGTLY
EYDYWGQGTQVTVSSGSGGGGSGGGGSGGGGSDTYCFDTYCFDTYCFLEHHHHHH
*

IGF2-7D12

MGLCGGELVDTLQFVCGDRGFYFSRPASRVSRRSRGIVEECCFRSCDLALLETYCATP
AKSEEFGSGGGGSGGGGSQVKLEESGGGSVQTGGSLRLTCAASGRTSRSYGMGWFR
QAPGKEREFVSGISWRGDSTGYADSVKGRFTISRDNAKNTVDLQMNSLKPEDTAIYY
CAAAAGSAWYGTLYEYDYWGQGTQVTVSSLEHHHHHH

IGF2-KN035

MGLCGGELVDTLQFVCGDRGFYFSRPASRVSRRSRGIVEECCFRSCDLALLETYCATP
AKSEEFGSGGGGSGGGGSGGGGSTGQVQLQESGGGLVQPGGSLRLSCAASGKMSSR
RCMAWFRQAPGKERERVAKLLTTSGSTYLADSVKGRFTISQNNAKSTVYLQMNSLK
PEDTAMYYCAADSFEDPTCTLVTSSGAFQYWGQGTQVTVSSGSMDPGGSLEHHHHH
H

7D12-Trop2$^{Nb108}$

MAMQVKLEESGGGSVQTGGSLRLTCAASGRTSRSYGMGWFRQAPGKEREFVSGISW
RGDSTGYADSVKGRFTISRDNAKNTVDLQMNSLKPEDTAIYYCAAAAGSAWYGTLY
EYDYWGQGTQVTVSSGSGGGGSGGGGSQVQLQESGGGLVQPGGSLRLSCAASGRIF

SSFAMAWFRQAPGKEREFVTAITWSGGSTYYADSVKGRFTISRDNAKNTVYLQMNS
LKPEDTAVYYCAAADDEGLSTVVYYYWGQGTQVTVSSLEHHHHHH

7D12-Trop2$^{\text{VNAR-5G8}}$

MAMQVKLEESGGGSVQTGGSLRLTCAASGRTSRSYGMGWFRQAPGKEREFVSGISW
RGDSTGYADSVKGRFTISRDNAKNTVDLQMNSLKPEDTAIYYCAAAAGSAWYGTLY
EYDYWGQGTQVTVSSGSGGGGSGGGGSQWVEQTPTTTTKEAGESLTINCVLRDSSC
PLASTYWYFTKKGATKKESLSNGGRYAETVNKASKSFSLRISDLRVEDSGTYHCKAV
NSWTNCAPLERYYEGGGTILSVKPAALEHHHHHH

Trop2$^{\text{VNAR-5G8}}$-EGFP

MGQWVEQTPTTTTKEAGESLTINCVLRDSSCPLASTYWYFTKKGATKKESLSNGGRY
AETVNKASKSFSLRISDLRVEDSGTYHCKAVNSWTNCAPLERYYEGGGTILSVKPAA
GGGGSGGGGSGSGGGGSVSKGEELFTGVVPILVELDGDVNGHKFSVSGEGEGDATY
GKLTLKFICTTGKLPVPWPTLVTTLTYGVQCFSRYPDHMKQHDFFKSAMPEGYVQER
TIFFKDDGNYKTRAEVKFEGDTLVNRIELKGIDFKEDGNILGHKLEYNYNSHNVYIM
ADKQKNGIKVNFKIRHNIEDGSVQLADHYQQNTPIGDGPVLLPDNHYLSTQSALSKD
PNEKRDHMVLLEFVTAAGITLGMDELYKLEHHHHHH

Recombinant human TROP2 protein

HTAAQDNCTCPTNKMTVCSPDGPGGRCQCRALGSGMAVDCSTLTSKCLLLKARMS
APKNARTLVRPSEHALVDNDGLYDPDCDPEGRFKARQCNQTSVCWCVNSVGVRRT
DKGDLSLRCDELVRTHHILIDLRHRPTAGAFNHSDLDAELRRLFRERYRLHPKFVAA
VHYEQPTIQIELRQNTSQKAAGDVDIGDAAYYFERDIKGESLFQGRGGLDLRVRGEPL
QVERTLIYYLDEIPPKFSMKRLT

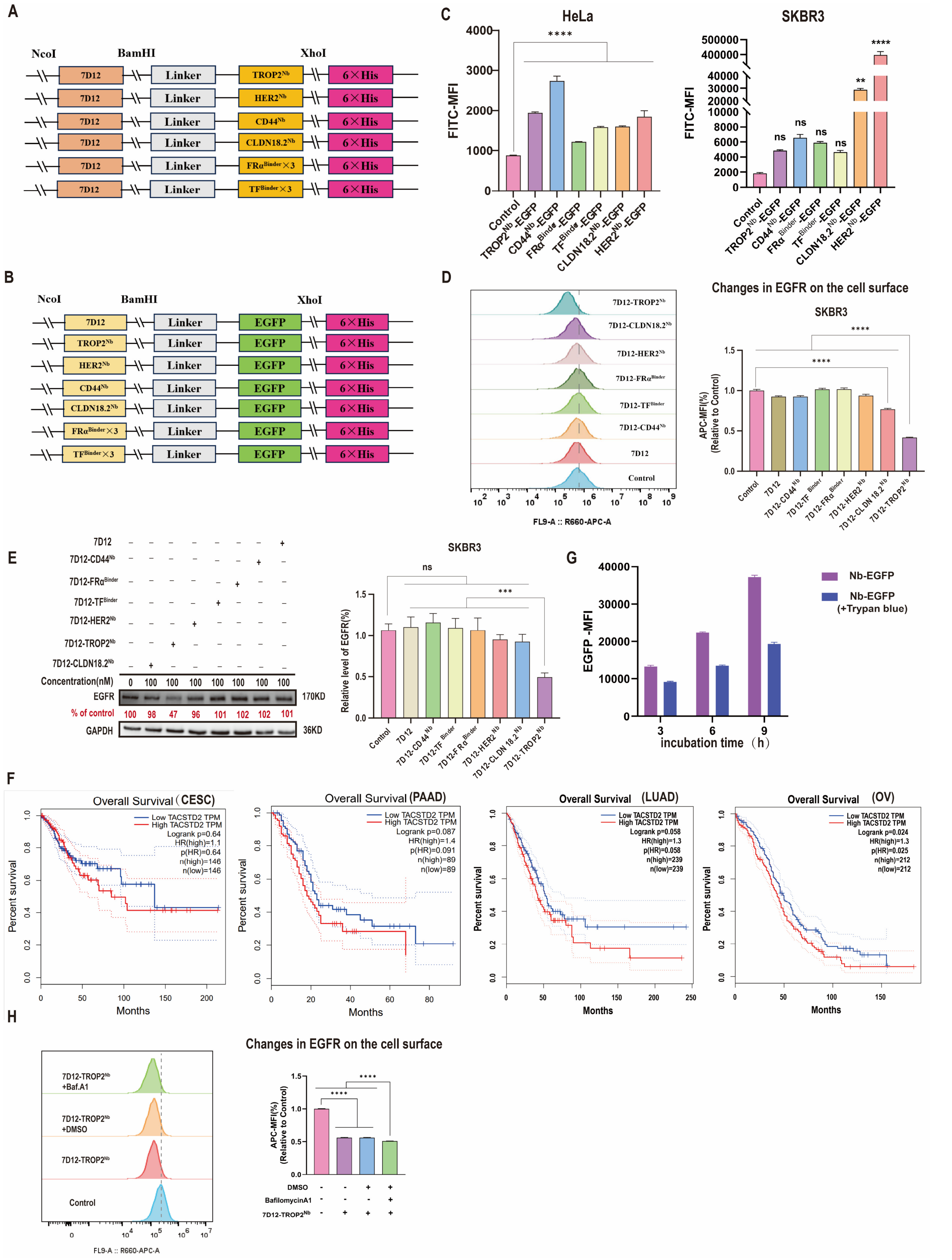


**Figure S1. Identification of TROP2 as an effective LTR with high clinical relevance.** (**A**) Constructs of nanobody-based heterobifunctional chimeras designed to recruit TROP2, HER2, CD44, CLDN18.2, FRα, or TF to EGFR. (**B**) Constructs of EGFP-fused nanobodies or binding peptides as probes against EGFR, TROP2, HER2, CD44, CLDN18.2, FRα, or TF. (**C**) Flow cytometry analysis of TROP2, CD44, FRα, TF, CLDN18.2, or HER2 expression on the surface of HeLa or SKBR3 cells following incubation with indicated protein probes. (**D**) Flow cytometry analysis of cell-surface EGFR levels in SKBR3 cells treated with 500 nM of 7D12, 7D12-CD44$^{Nb}$, 7D12-TF$^{Binder}$, 7D12-FRα$^{Binder}$, 7D12-HER2$^{Nb}$, 7D12-CLDN18.2$^{Nb}$, or 7D12-TROP2$^{Nb}$ for 24 h. (**E**) Western blot analysis of EGFR levels in SKBR3 cells treated with 100 nM of 7D12, 7D12-CD44$^{Nb}$, 7D12-TF$^{Binder}$, 7D12-FRα$^{Binder}$, 7D12-HER2$^{Nb}$, 7D12-CLDN18.2$^{Nb}$, or 7D12-TROP2$^{Nb}$ for 24 h. (**F**) Kaplan–Meier analysis of correlation between TACSTD2 expression and overall survival in patients with CESC (cervical squamous cell carcinoma), PAAD (pancreatic adenocarcinoma), LUAD (lung adenocarcinoma), or OV (ovarian serous cystadenocarcinoma). (**G**) Flow cytometry analysis of cells incubated with 1 μM of TROP2$^{Nb}$-EGFP for 3, 6, or 9 h, then treated with or without trypan blue. (**H**) Flow cytometry analysis and quantification of cell-surface EGFR levels in HeLa cells treated with 100 nM of 7D12-TROP2$^{Nb}$ for 24 h in the presence or absence of bafilomycin A1. Data are presented as mean ± standard deviation (SD) (n = 3 biologically independent replicates) when relevant. *P* values were determined by one-way analysis of variance (ANOVA) with Turkey's *post hoc* test. **P* < 0.05, ***P* < 0.01, ****P* < 0.001, *****P* < 0.0001.

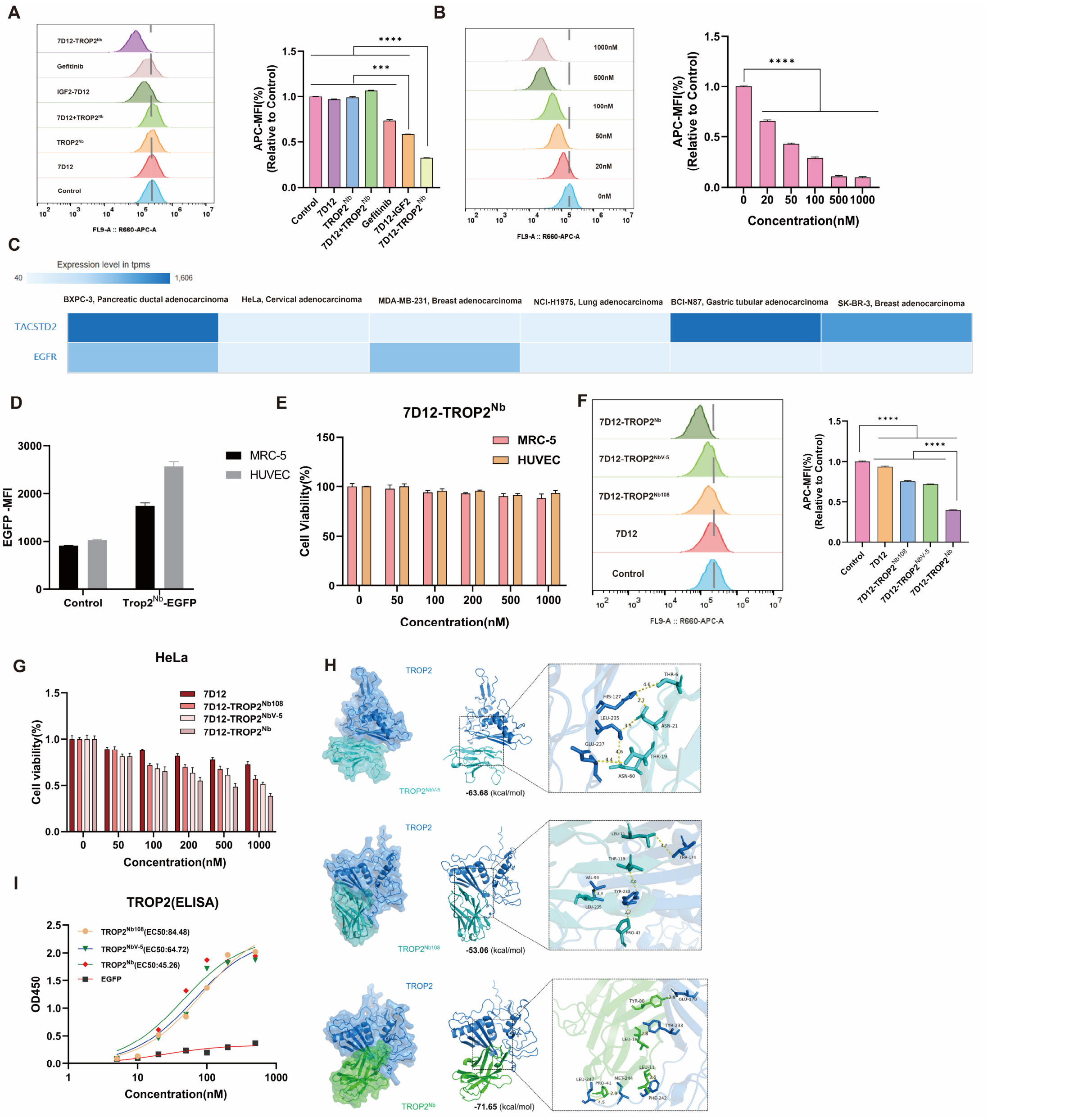


**Figure S2. 7D12-TROP2$^{Nb}$-induced EGFR internalization and degradation.** (**A** and **B**) Flow cytometry analysis of cell-surface EGFR levels in HeLa cells treated for 24 h with 100 nM of 7D12, TROP2$^{Nb}$, 7D12 + TROP2$^{Nb}$, 7D12-IGF2, gefitinib, or 7D12-TROP2$^{Nb}$ (**A**) or with serial concentrations of 7D12-TROP2$^{Nb}$ (**B**). (**C**) Heat map of TACSTD2 and EGFR expression in BxPC-3, HeLa, MDA-MB-231, NCI-H1975, NCI-N87, and SK-BR-3 cells. Data were obtained from the Cancer Cell Line Encyclopedia. (**D**) Flow cytometry analysis of TROP2 expression in MRC-5 and HUVEC cells. (**E**) MTT assay of MRC-5 and HUVEC cells treated with serial concentrations of 7D12-TROP2$^{Nb}$. (**F**) Flow cytometry analysis of cell-surface EGFR levels in HeLa cells treated with 500 nM of 7D12, 7D12-TROP2$^{NbV-5}$, 7D12-TROP2$^{Nb108}$, or 7D12-TROP2$^{Nb}$ for 24 h. (**G**) MTT assay of HeLa cells treated with serial concentrations of 7D12, 7D12-TROP2$^{NbV-5}$, 7D12-TROP2$^{Nb108}$, or 7D12-TROP2$^{Nb}$ for 7 d. (**H**)

Molecular docking of TROP2 with TROP2$^{Nb}$, TROP2$^{NbV\text{-}5}$, or TROP2$^{Nb108}$. (**I**) ELISA analysis of TROP2$^{Nb}$-EGFP, TROP2$^{NbV\text{-}5}$-EGFP, TROP2$^{Nb108}$-EGFP, or EGFP binding to TROP2. Data are presented as mean ± SD (n = 3 biologically independent replicates) when relevant. *P* values were determined by one-way ANOVA with Turkey's *post hoc* test. ***$P < 0.001$, ****$P < 0.0001$.

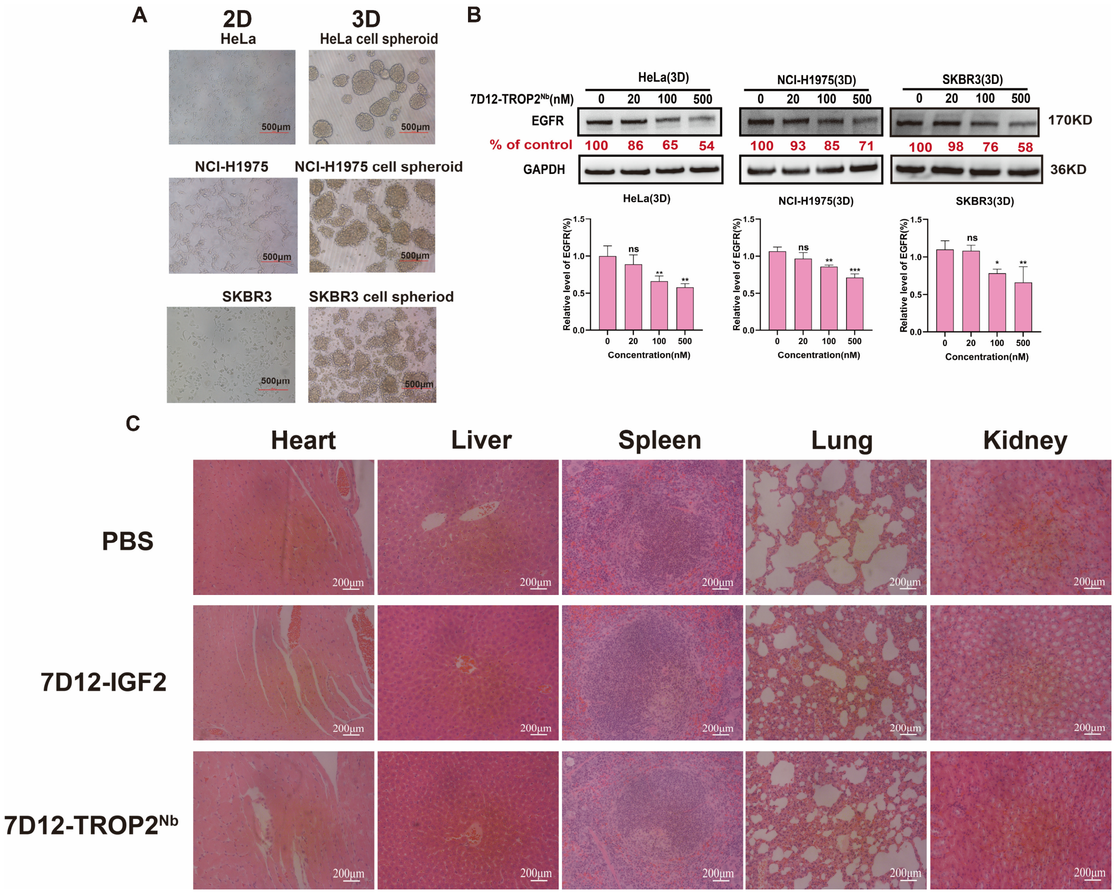


**Figure 3. 7D12-TROP2$^{Nb}$-mediated EGFR degradation in 3D tumor spheroids and assessment of *in vivo* safety.** (**A**) Bright field images of HeLa, H1975, and SKBR3 cells in 2D and 3D culture. Scale bar: 500 μm. (**B**) Western blot analysis and quantification of EGFR levels in HeLa, H1975, and SKBR3 spheroids treated with different concentrations of 7D12-TROP2$^{Nb}$ for 24 h (n = 3 biologically independent replicates). (**C**) Haematoxylin and eosin (H&E) staining of major organs, including heart, liver, spleen, lung, and kidney, collected from mice treated with PBS, 7D12-IGF2, or 7D12-TROP2$^{Nb}$. Scale bar: 200 μm. Data are presented as mean ± SD when relevant. *P* values were determined by one-way ANOVA with Turkey's *post hoc* test. ns, no significance; **P* < 0.05; ***P* < 0.01; ****P* < 0.001.

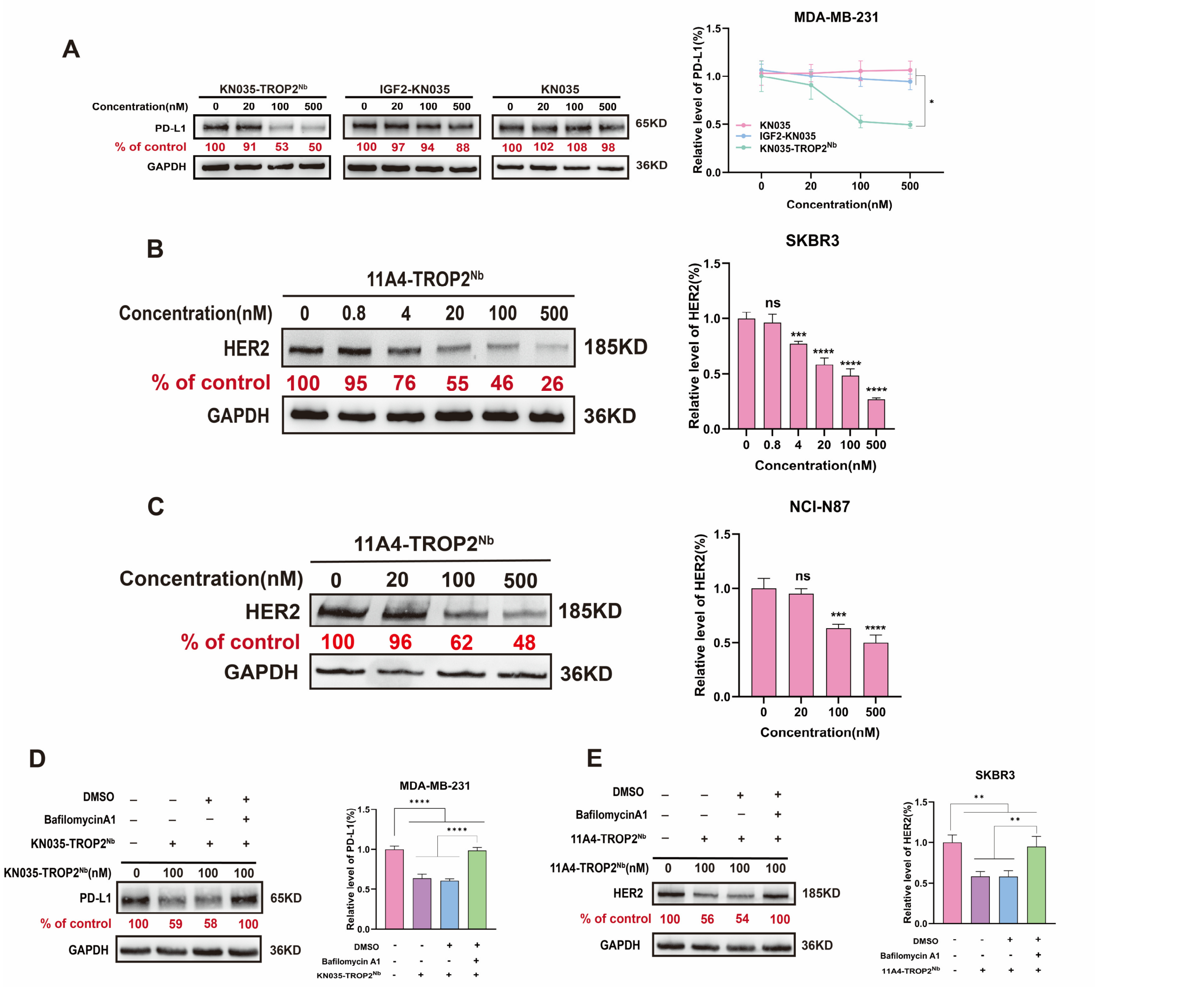


**Figure S4. TRTAC-mediated degradation of PD-L1 and HER2.** (**A**) Western blot analysis and quantification of PD-L1 levels in MDA-MB-231 cells treated for 24 h with serial concentrations of KN035-TROP2$^{Nb}$, IGF2-KN035, or KN035. (**B** and **C**) Western blot analysis and quantification of HER2 levels in SKBR3 cells (**B**) or NCI-N87 cells (**C**) treated for 24 h with serial concentrations of 11A4-TROP2$^{Nb}$. (**D**) Western blot analysis and quantification of PD-L1 levels in MDA-MB-231 cells treated with 100 nM of KN035-TROP2$^{Nb}$ for 24 h in the presence or absence of bafilomycin A1 (50 nM). (**E**) Western blot analysis and quantification of HER2 levels in SKBR3 cells treated with 100 nM of 11A4-TROP2$^{Nb}$ for 24 h in the presence or absence of bafilomycin A1. Data are presented as mean ± SD (n = 3 biologically independent replicates) when relevant. *P* values were determined by one-way ANOVA with Turkey's *post hoc* test. ns, no significance; **P* < 0.05; ***P* < 0.01; ****P* < 0.001; *****P* < 0.0001.

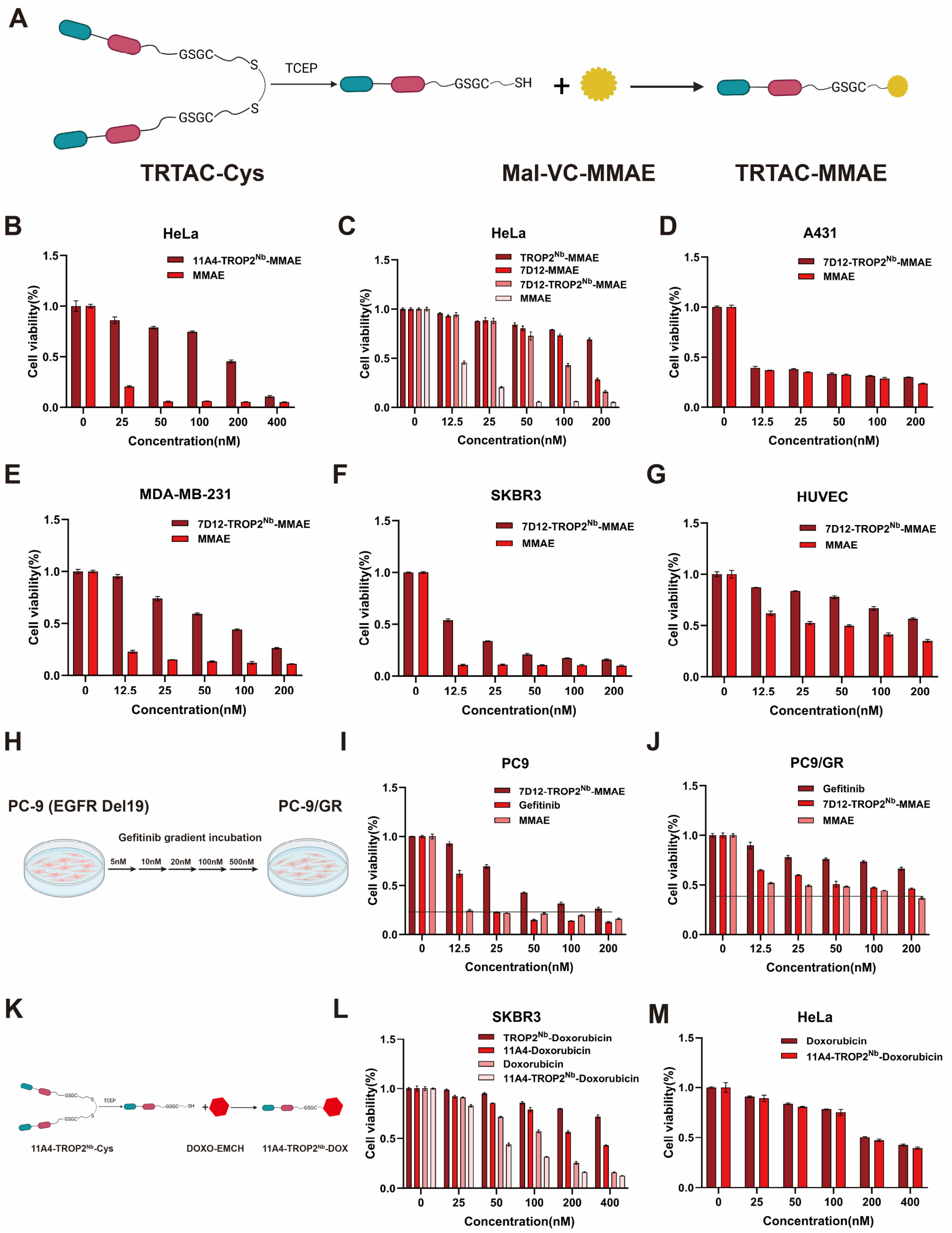


**Figure S5. Characterization of TRTAC-DCs.** (**A**) Synthesis of 11A4-TROP2$^{Nb}$-MMAE. (**B** and **C**) MTT assay of HeLa cells treated for 72 h with serial concentrations of MMAE or 11A4-TROP2$^{Nb}$-MMAE (**B**) or with serial concentrations of MMAE, 7D12-MMAE, or 7D12-TROP2$^{Nb}$-MMAE (**C**). (**D-G**) MTT assay of A431 (**D**), MDA-MB-231 (**E**), SKBR3 (**F**), or HUVEC (**G**) cells treated with serial concentrations of MMAE or 7D12-TROP2$^{Nb}$-MMAE for 72 h. (**H**) Schematic illustration of establishment of gefitinib-resistant PC-9/GR cell strains. Gefitinib-sensitive PC-9 cells were cultured with stepwise increasing concentrations of gefitinib for generating resistant PC-9/GR cells. (**I** and **J**) MTT assay of PC-9 (**I**) or PC-9/GR (**J**) cells treated with serial concentrations of gefitinib, 7D12-TROP2$^{Nb}$-MMAE, or free MMAE for 72 h. (**K**) Synthesis of 11A4-TROP2$^{Nb}$-DOX. (**L** and **M**) MTT assay of SKBR3 (**L**) or HeLa

(**M**) cells treated with serial concentrations of 11A4-TROP2$^{Nb}$-DOX or indicated controls for 72 h. Data are presented as mean ± SD (n = 3 biologically independent replicates).

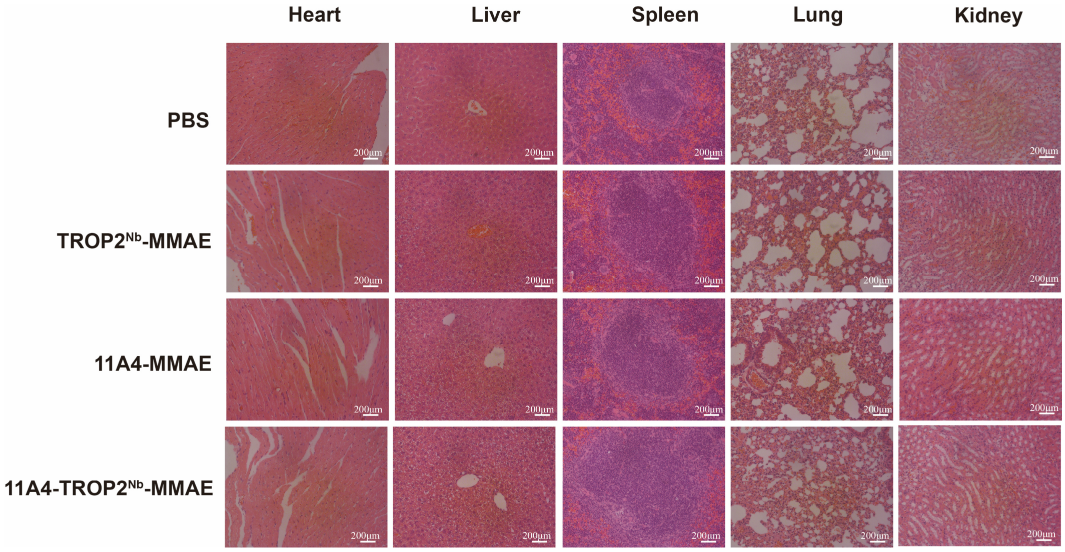


**Figure S6. Assessment of *in vivo* safety of 11A4- TROP2$^{Nb}$-MMAE.** H&E staining of major organs, including heart, liver, spleen, lung and kidney, collected from mice treated with PBS, TROP2$^{Nb}$-MMAE, 11A4-MMAE or 11A4- TROP2$^{Nb}$-MMAE. Scale bar: 200 µm.

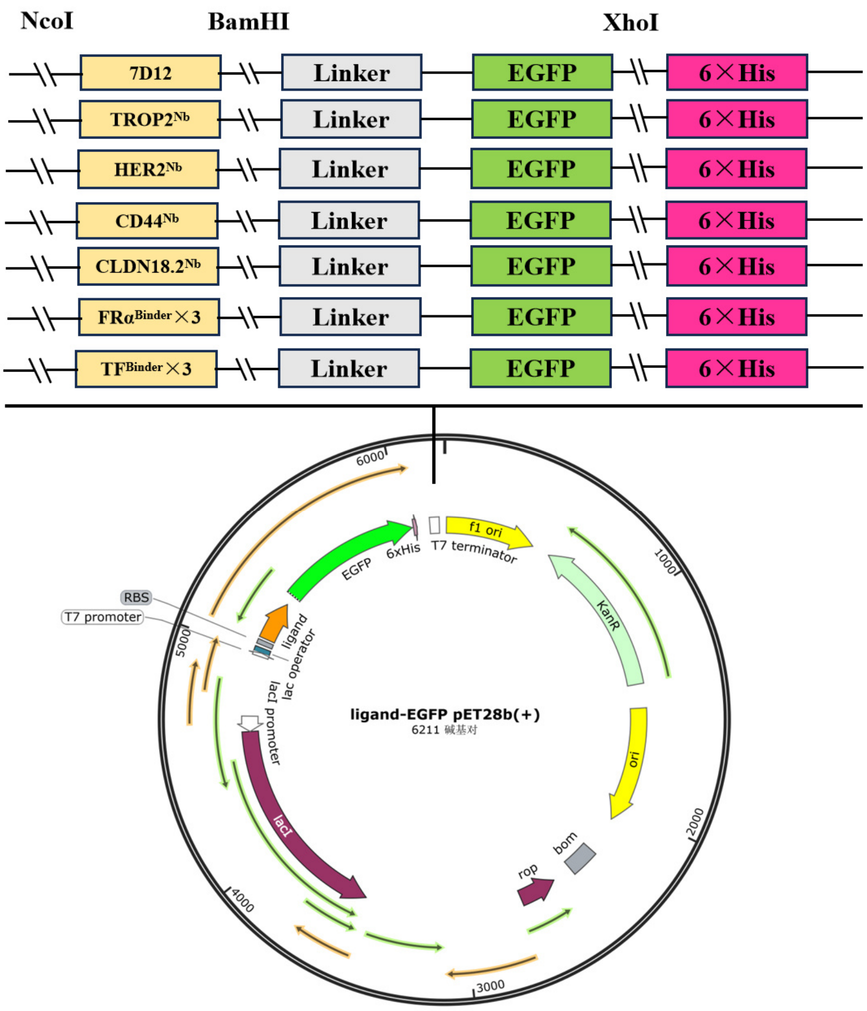


**Figure S7. Plasmid design for expression of EGFP-fused nanobodies or binding peptides against membrane receptors.**

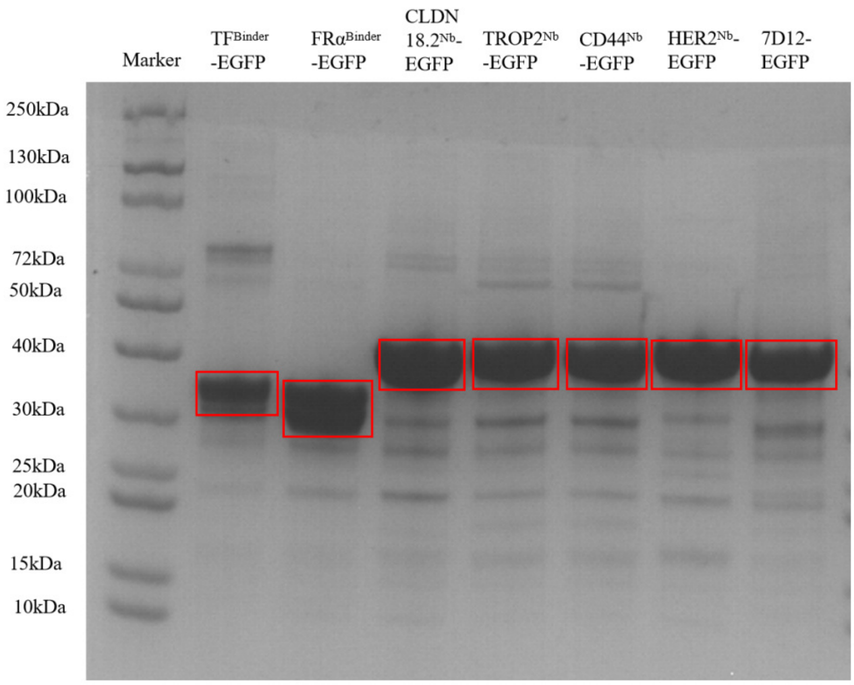


**Figure S8. SDS-PAGE of purified proteins.** Lane 1: Marker; Lane 2: TF$^{Binder}$-EGFP; Lane 3: FRα$^{Binder}$-EGFP; Lane 4: CLDN$^{18.2Nb}$-EGFP; Lane 5: TROP2$^{Nb}$-EGFP; Lane 6: CD44$^{Nb}$-EGFP; Lane 7: HER2$^{Nb}$-EGFP; Lane 8: 7D12-EGFP.

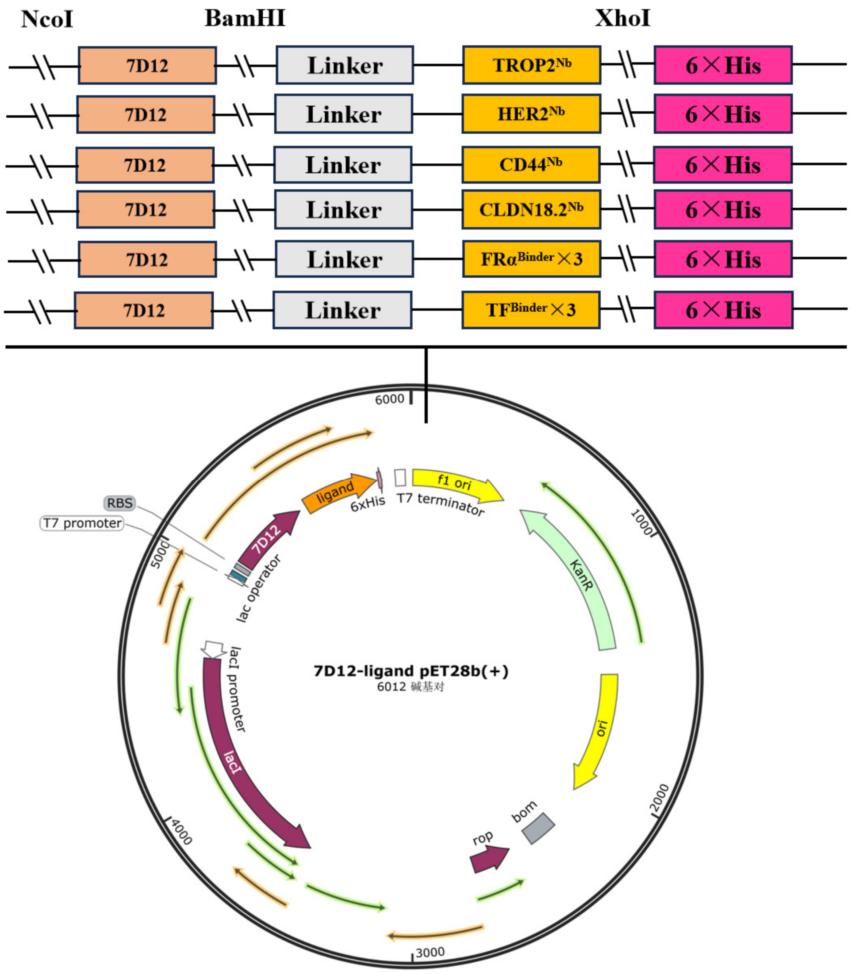


**Figure S9. Plasmid design for expression of 7D12–ligand fusion proteins**

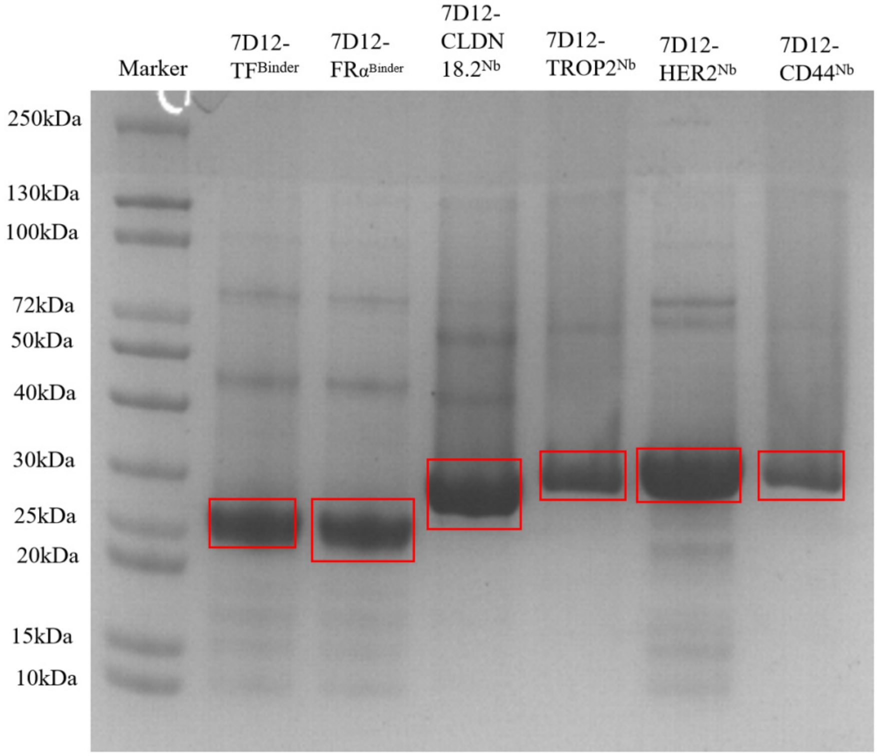


**Figure S10. SDS-PAGE of purified proteins.** Lane 1: Marker; Lane 2: 7D12-TF$^{Binder}$; Lane 3: 7D12-FRα$^{Binder}$; Lane 4: 7D12-CLDN$^{18.2Nb}$; Lane 5: 7D12-TROP2$^{Nb}$; Lane 6: 7D12-CD44$^{Nb}$; Lane 7: 7D12-HER2$^{Nb}$; Lane 8: 7D12- CD44$^{Nb}$.

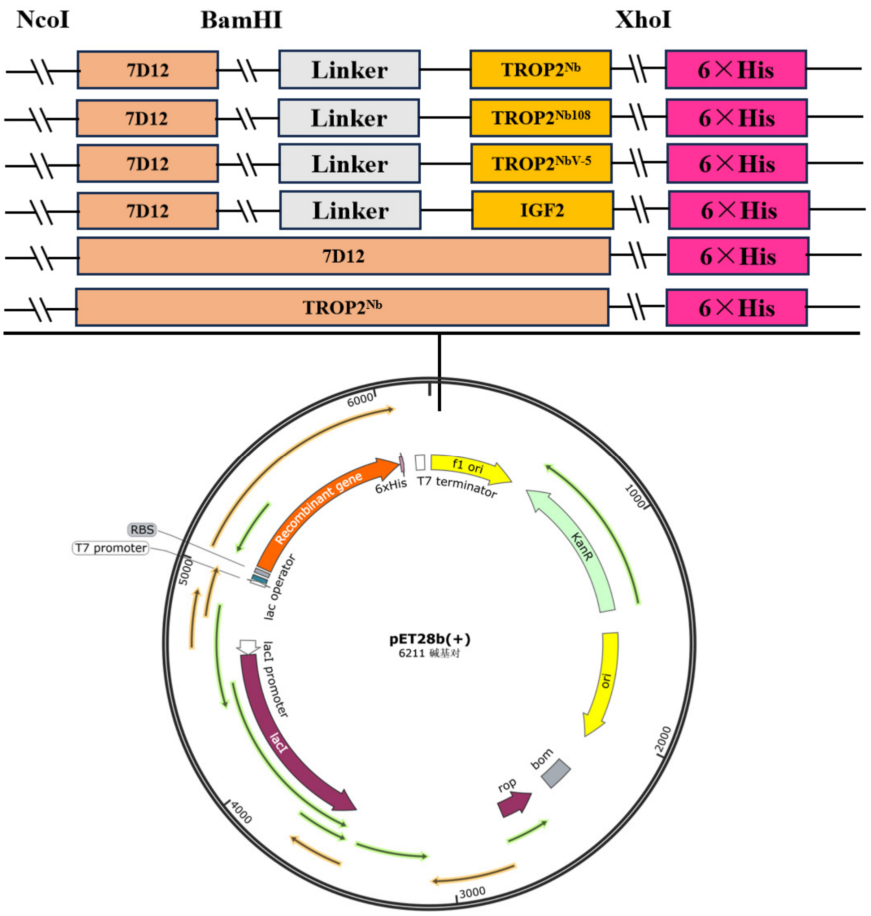


**Figure S11. Plasmid design for expression of 7D12-TROP2$^{Nb}$, 7D12-TROP2$^{Nb108}$, 7D12-TROP2$^{NbV-5}$, and IGF2-7D12.**

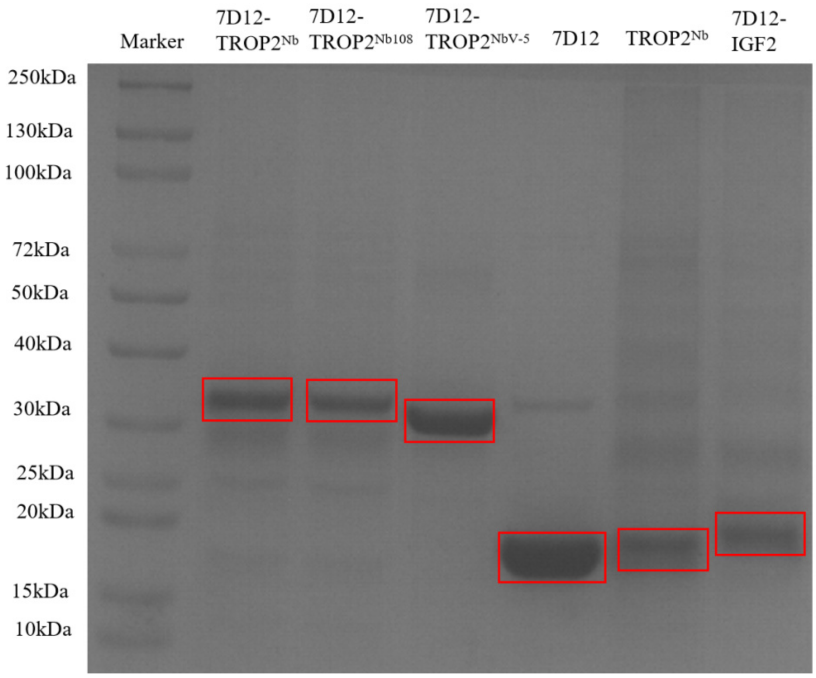


**Figure S12. SDS-PAGE of purified proteins.** Lane 1: Marker; Lane 2: 7D12-TROP2$^{Nb}$; Lane 3: 7D12-TROP2$^{Nb108}$; Lane 4: 7D12-TROP2$^{NbV\text{-}5}$; Lane 5: 7D12; Lane 6: TROP2$^{Nb}$; Lane 8: 7D12-IGF2.

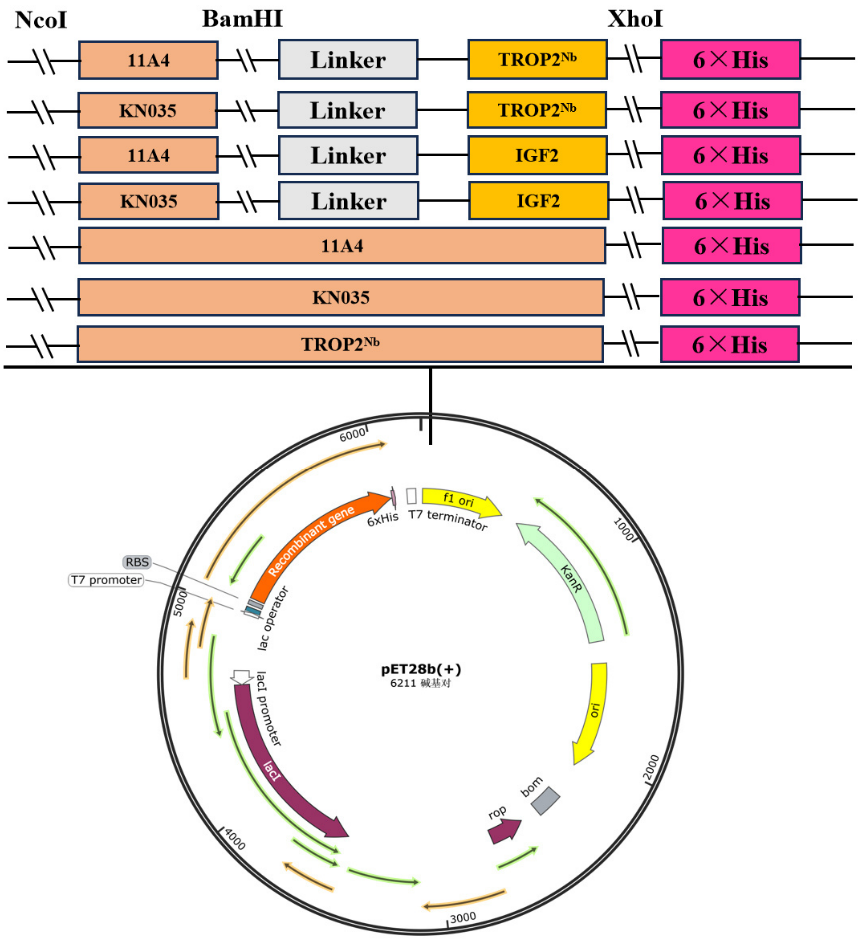


**Figure S13. Plasmid design for expression of HER2- and PD-L1-targeted TRTACs and corresponding controls.**

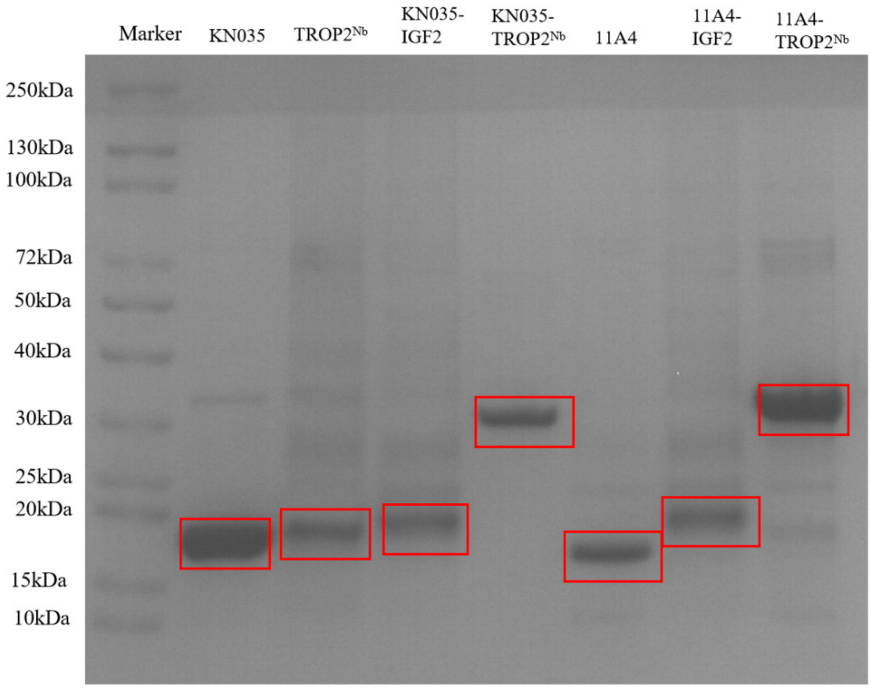


**Figure S14. SDS-PAGE of purified proteins.** Lane 1: Marker; Lane 2: KN035; Lane 3: TROP2$^{Nb}$; Lane 4: KN035-IGF2; Lane 5: KN035-TROP2$^{Nb}$; Lane 6: 11A4; Lane 7: 11A4-IGF2; Lane 8: 11A4-TROP2$^{Nb}$.

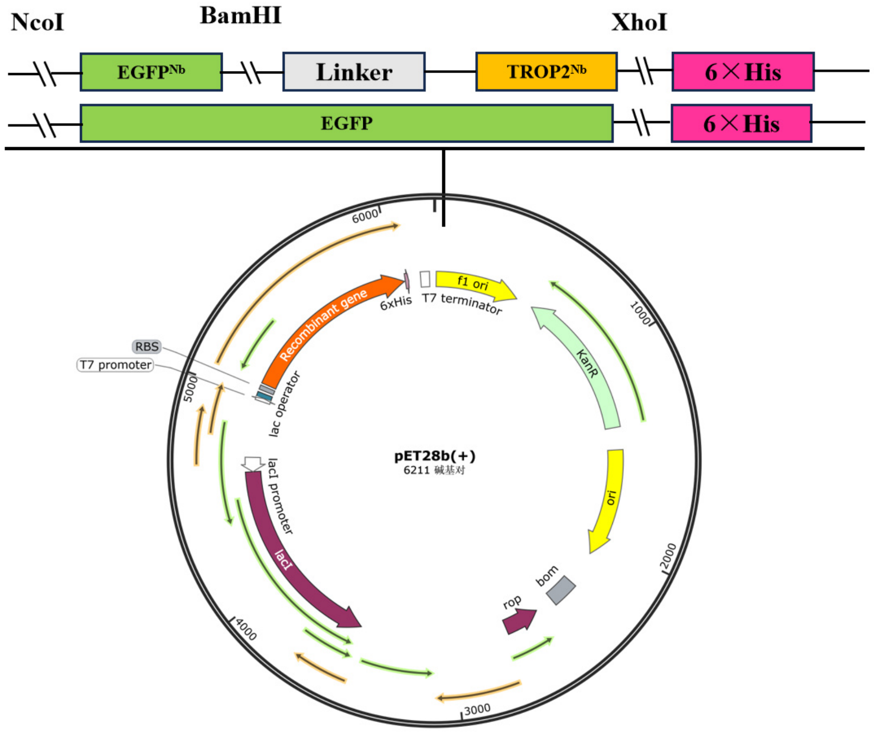


**Figure S15. Plasmid design for expression of EGFP-targeted TRTAC and soluble EGFP.**

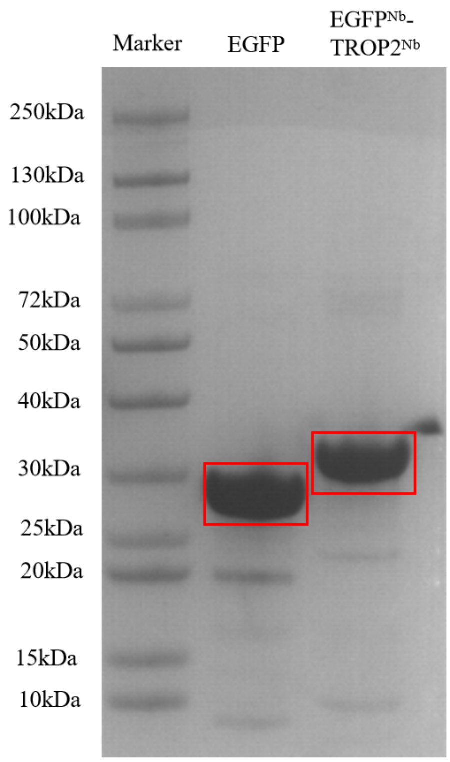


**Figure S16. SDS-PAGE of purified proteins.** Lane 1: Marker; Lane 2: EGFP; Lane 3: EGFP$^{Nb}$-TROP2$^{Nb}$.

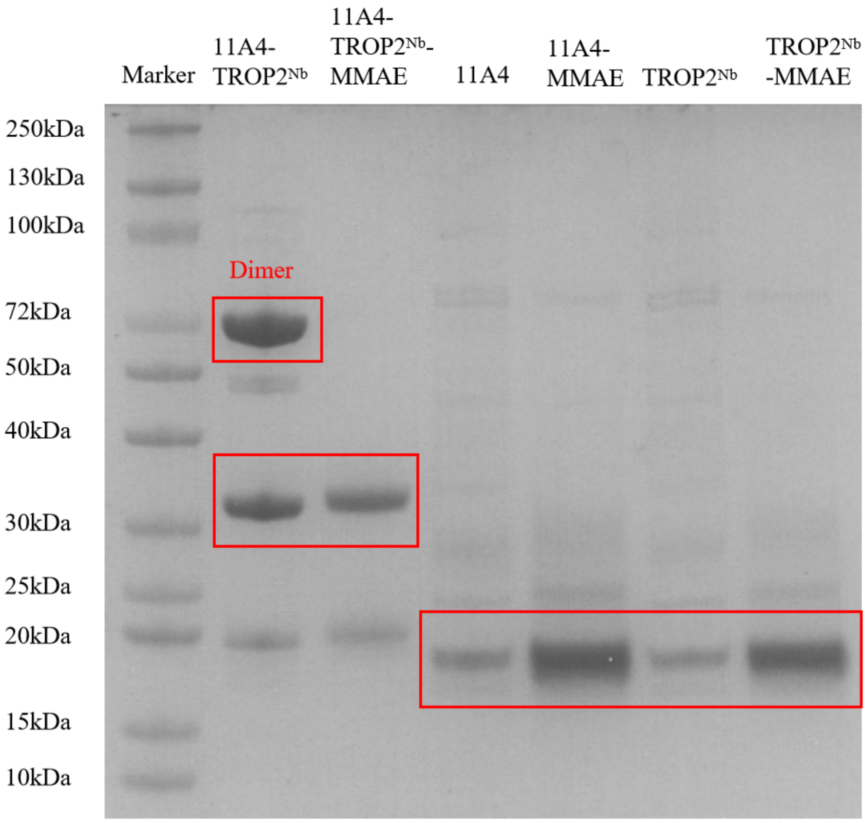


**Figure S17. SDS-PAGE of purified proteins.** Lane 1: Marker; Lane 2: 11A4-TROP2$^{Nb}$; Lane 3: 11A4-TROP2$^{Nb}$-MMAE; Lane 4: 11A4; Lane 5: 11A4-MMAE; Lane 6: TROP2$^{Nb}$; Lane 7: TROP2$^{Nb}$-MMAE.

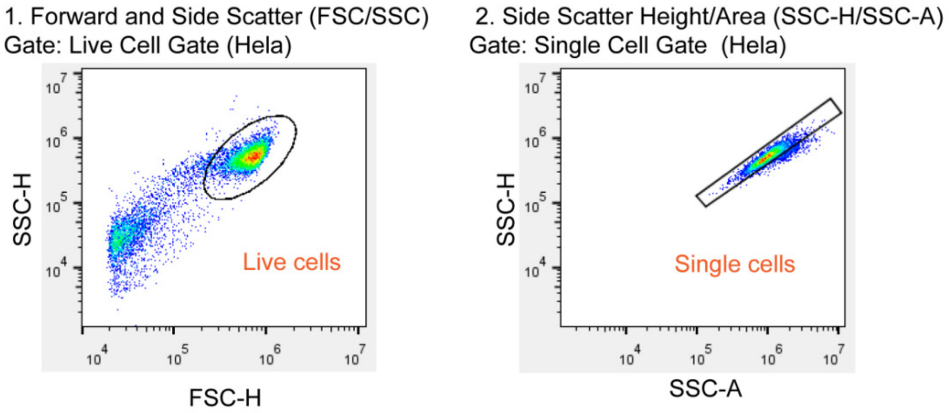


**Figure S18. Gating strategy for flow cytometry.**

**WB raw data**

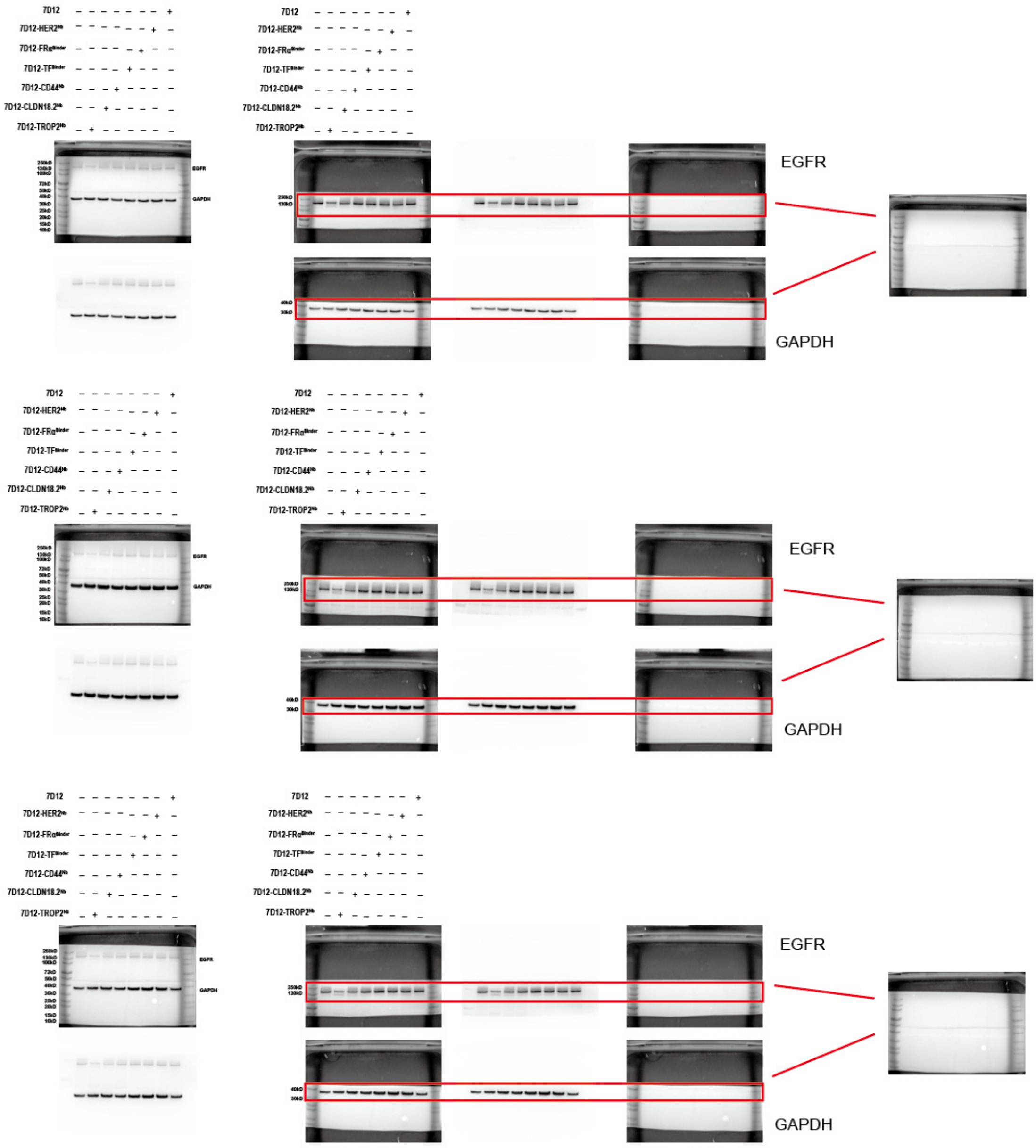


**Figure 1D**

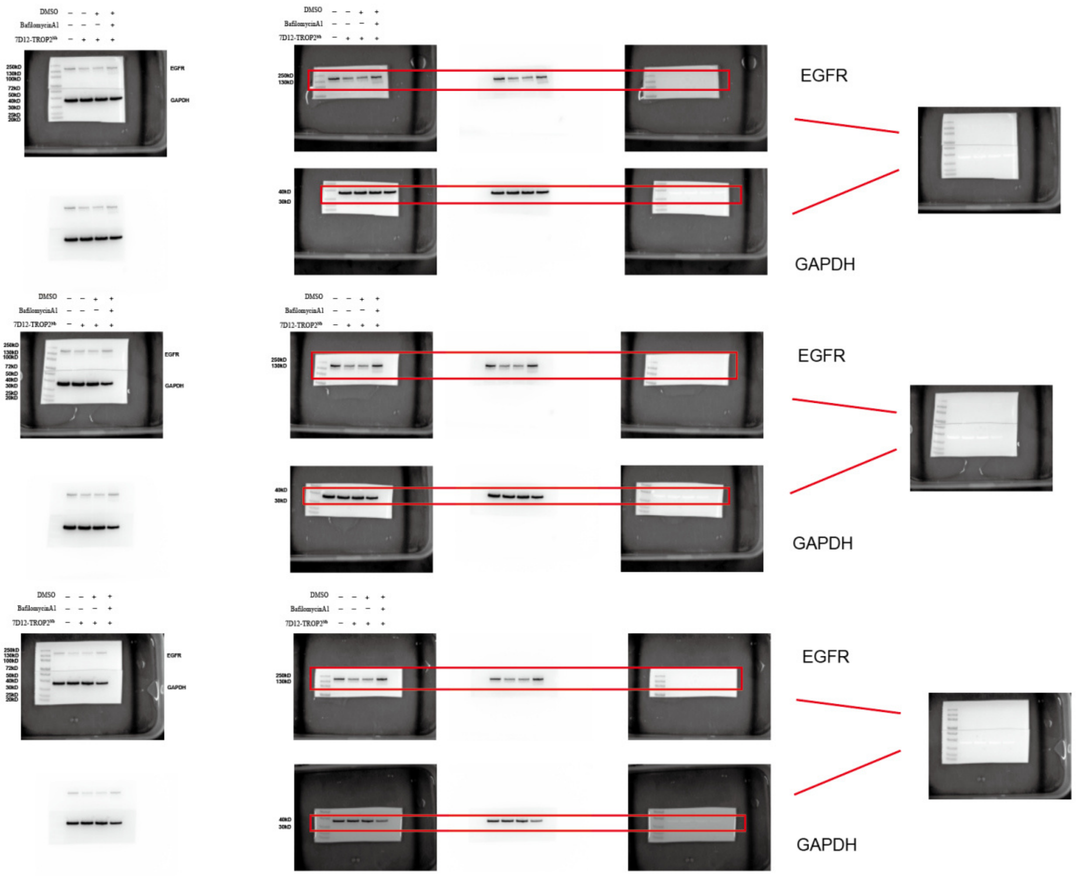


**Figure 1H**

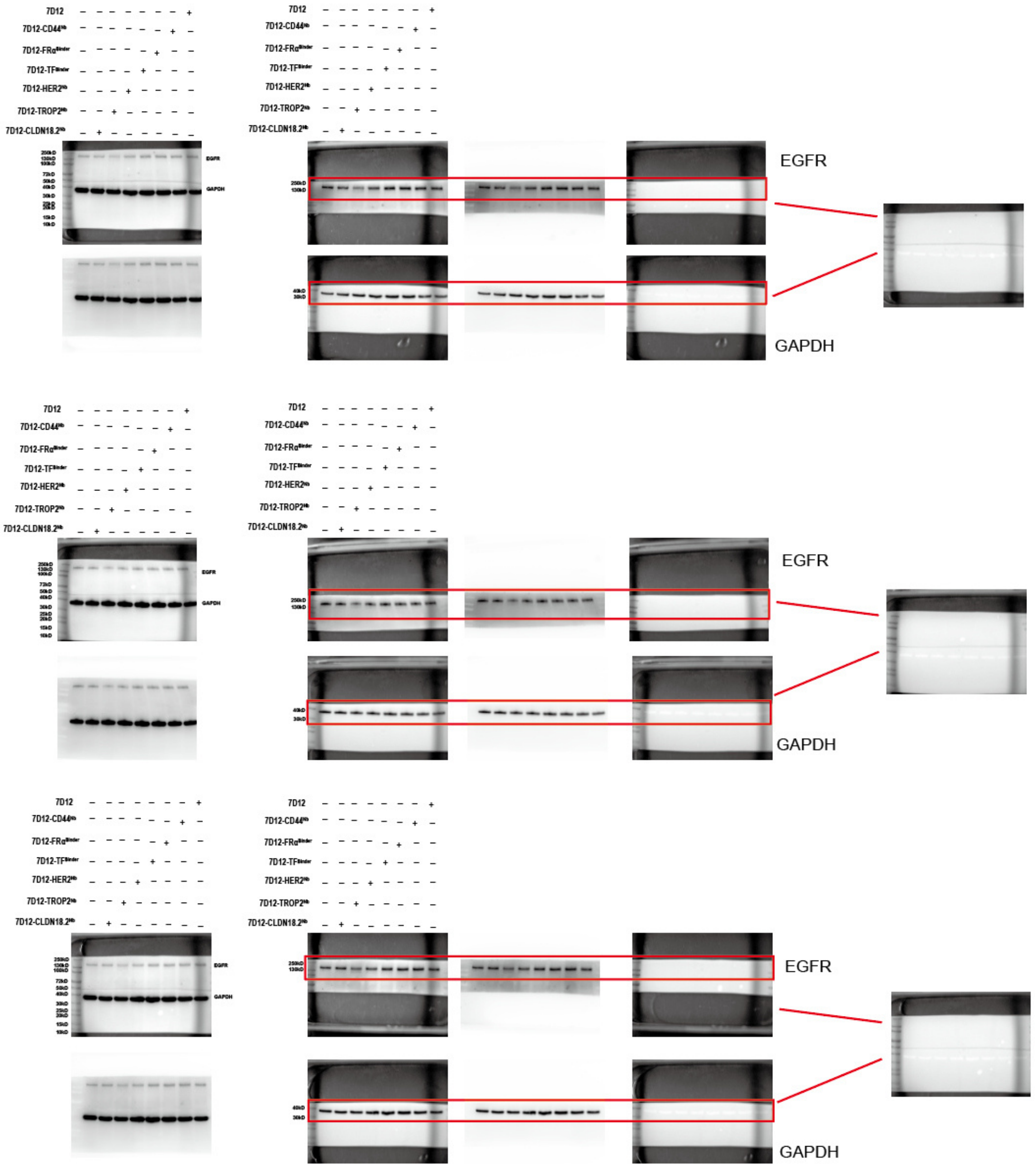
7D12
7D12-CD44
7D12-FRα
7D12-TF
7D12-HER2
7D12-TROP2
7D12-CLDN18.2
EGFR
GAPDH

**Figure S1E**

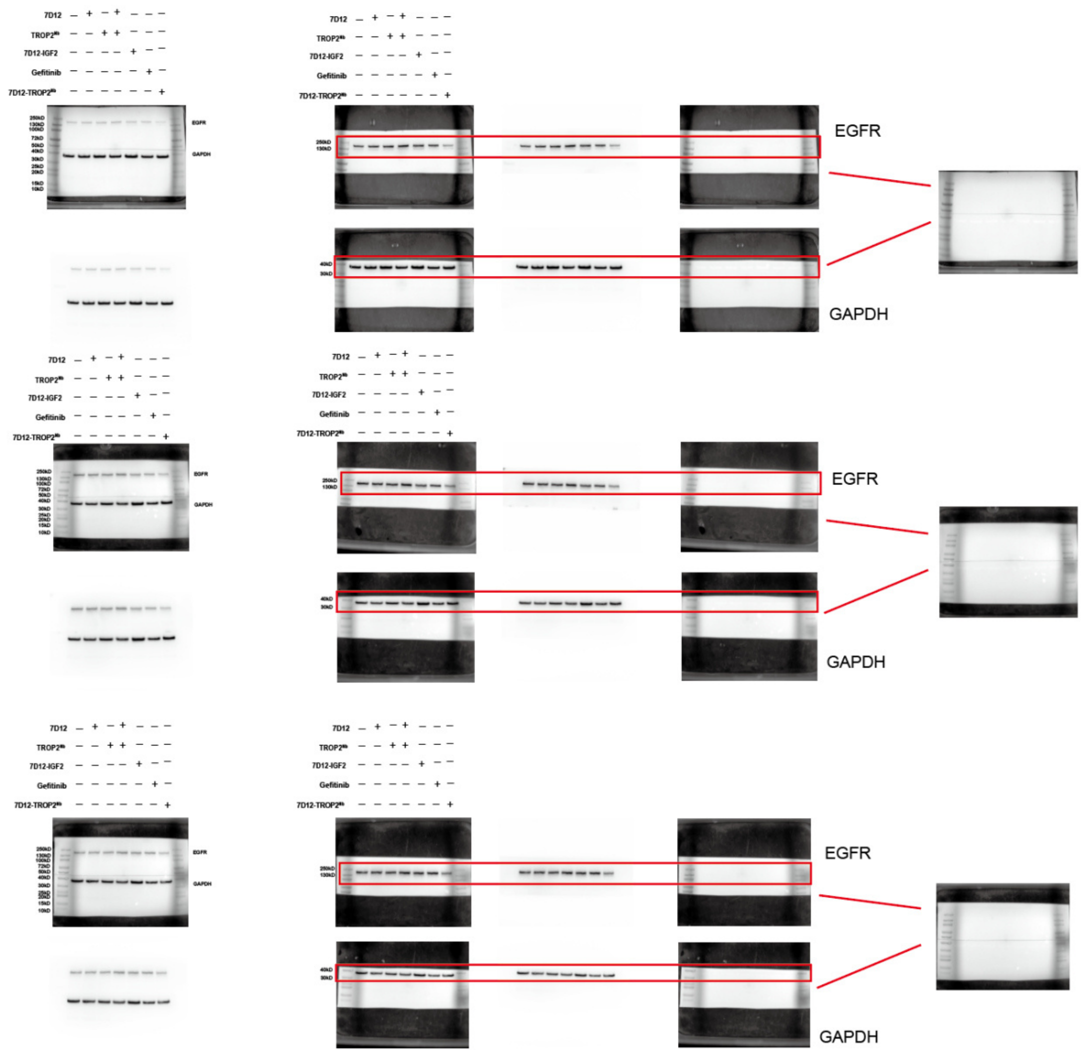


**Figure 2A**

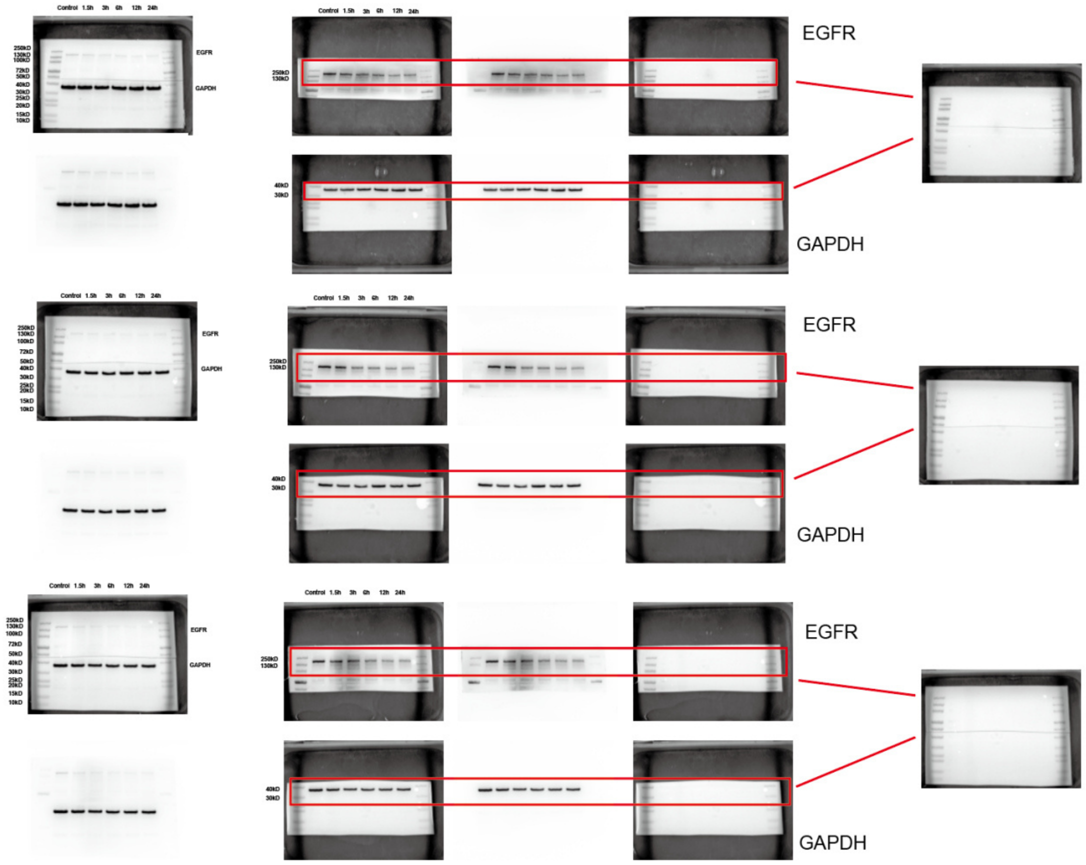


Figure 2B

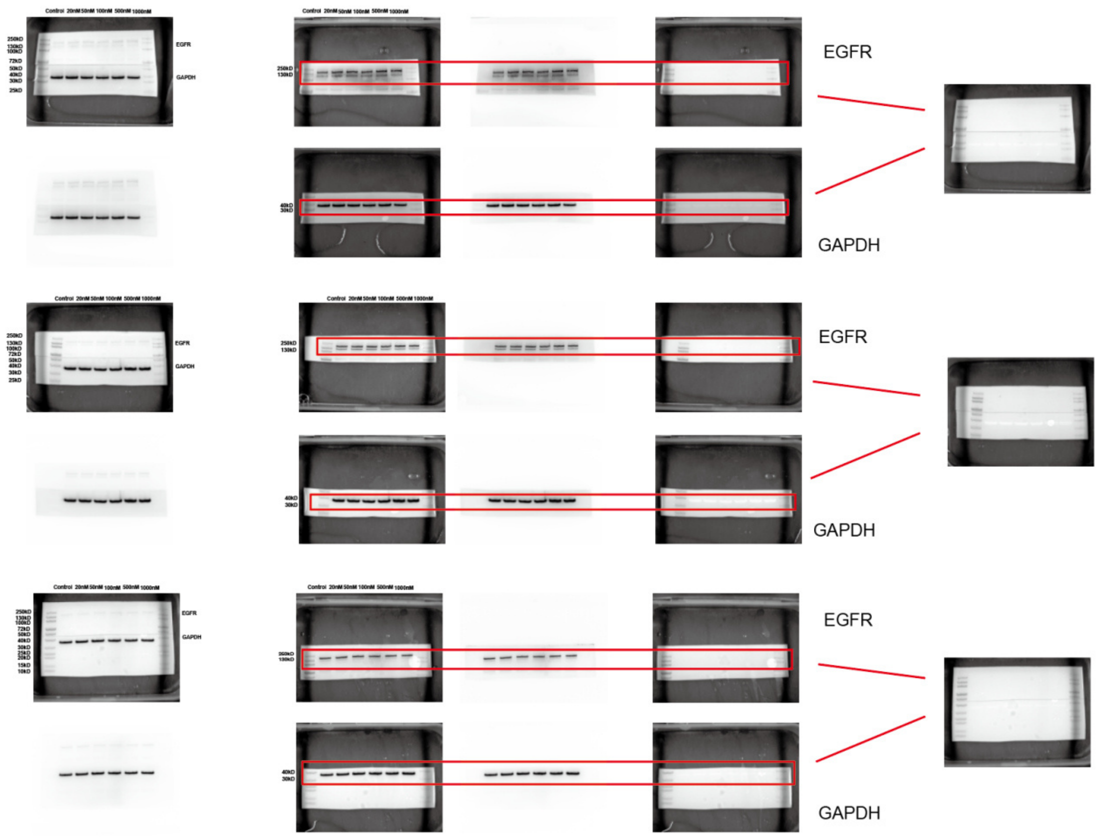


**Figure 2C 7D12**

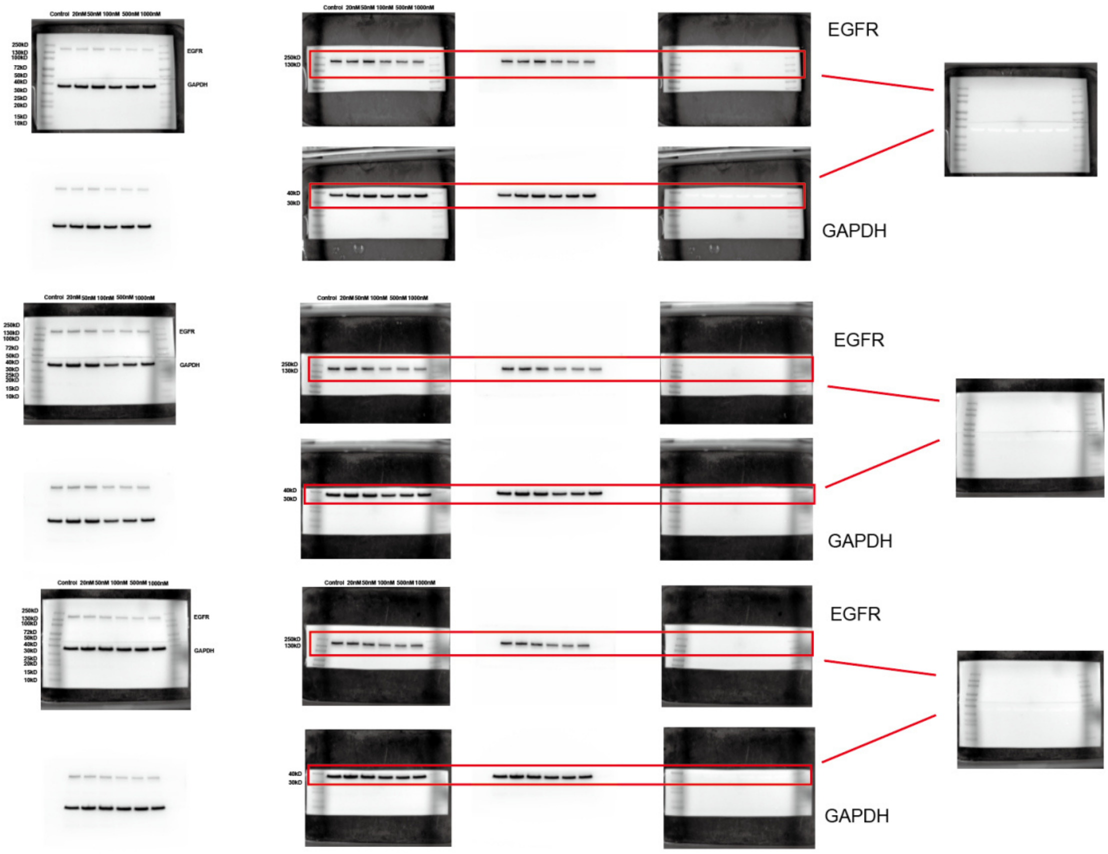


**Figure 2C 7D12-IGF2**

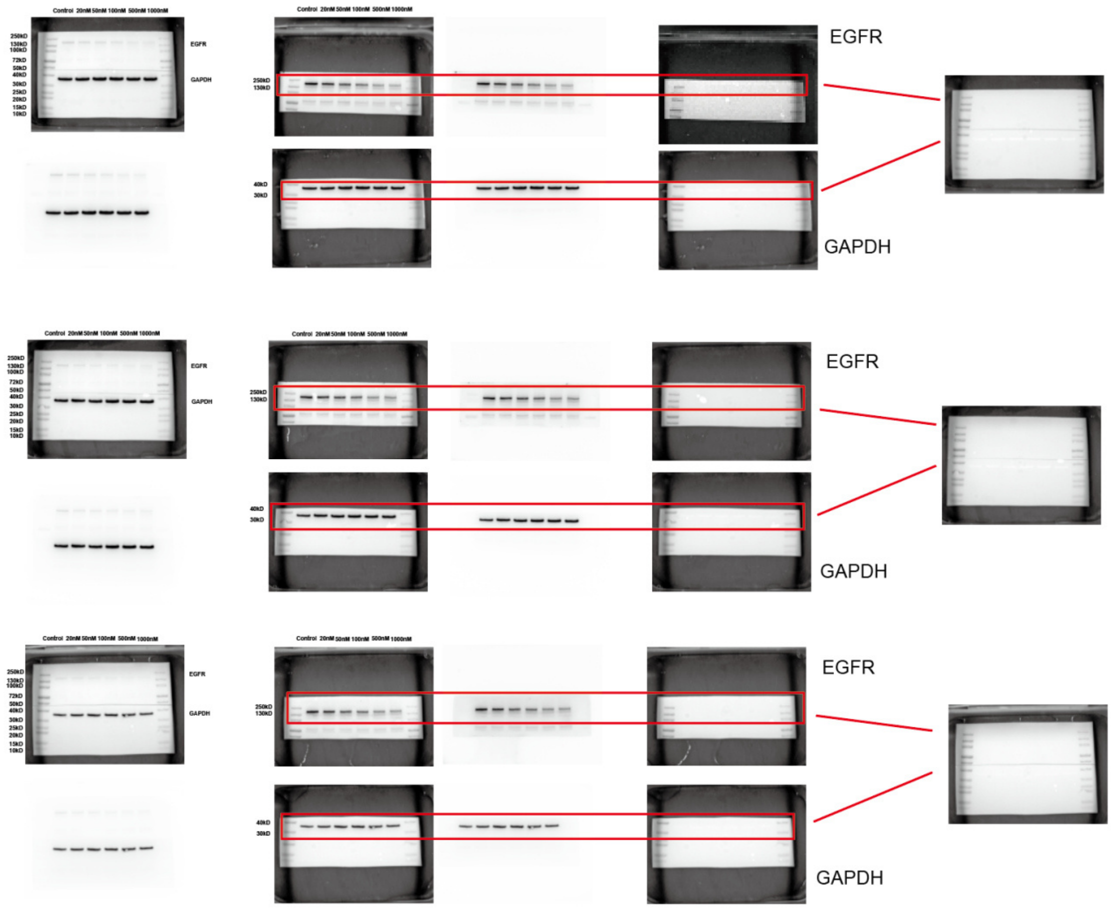


**Figure 2C 7D12-TROP2$^{Nb}$**

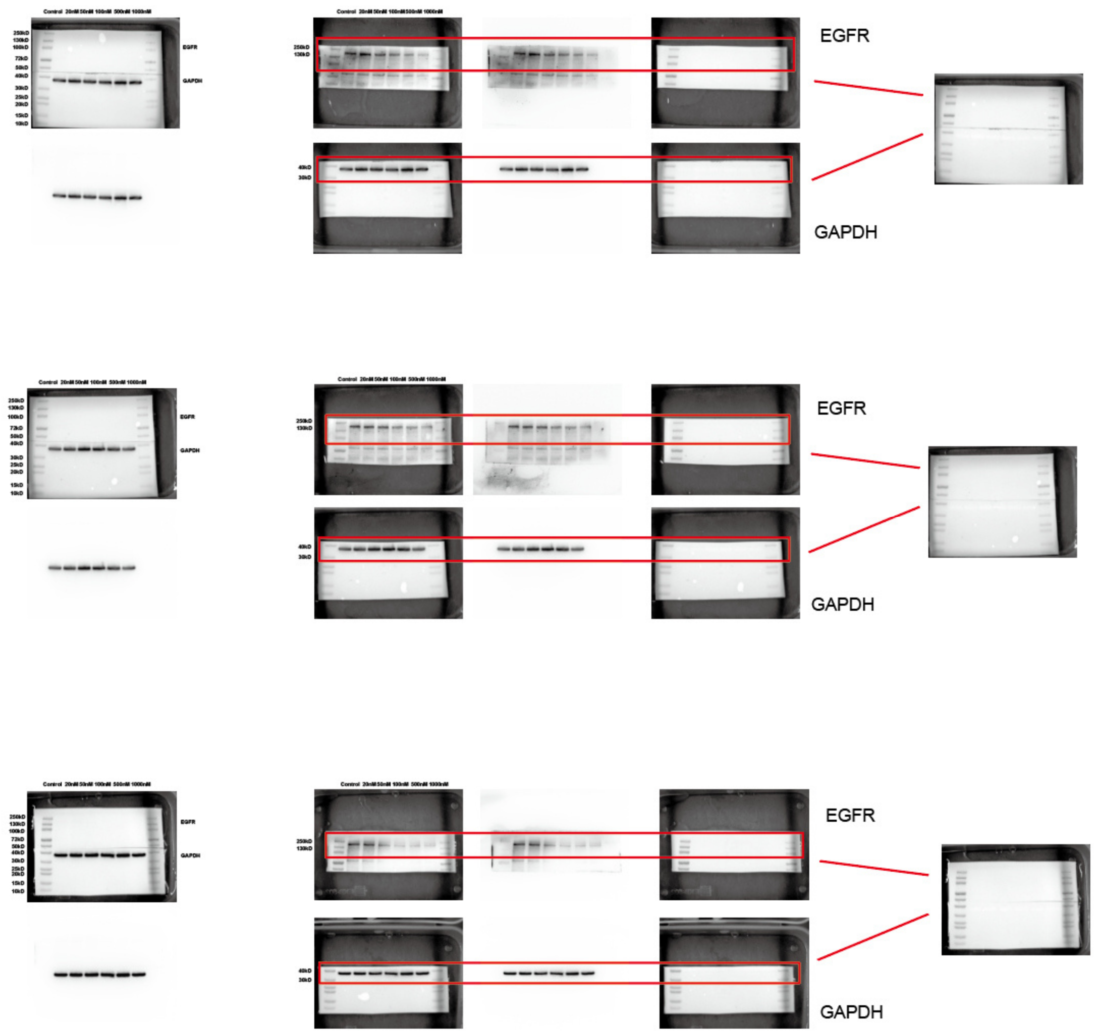


**Figure 2D SKBR3 cells**

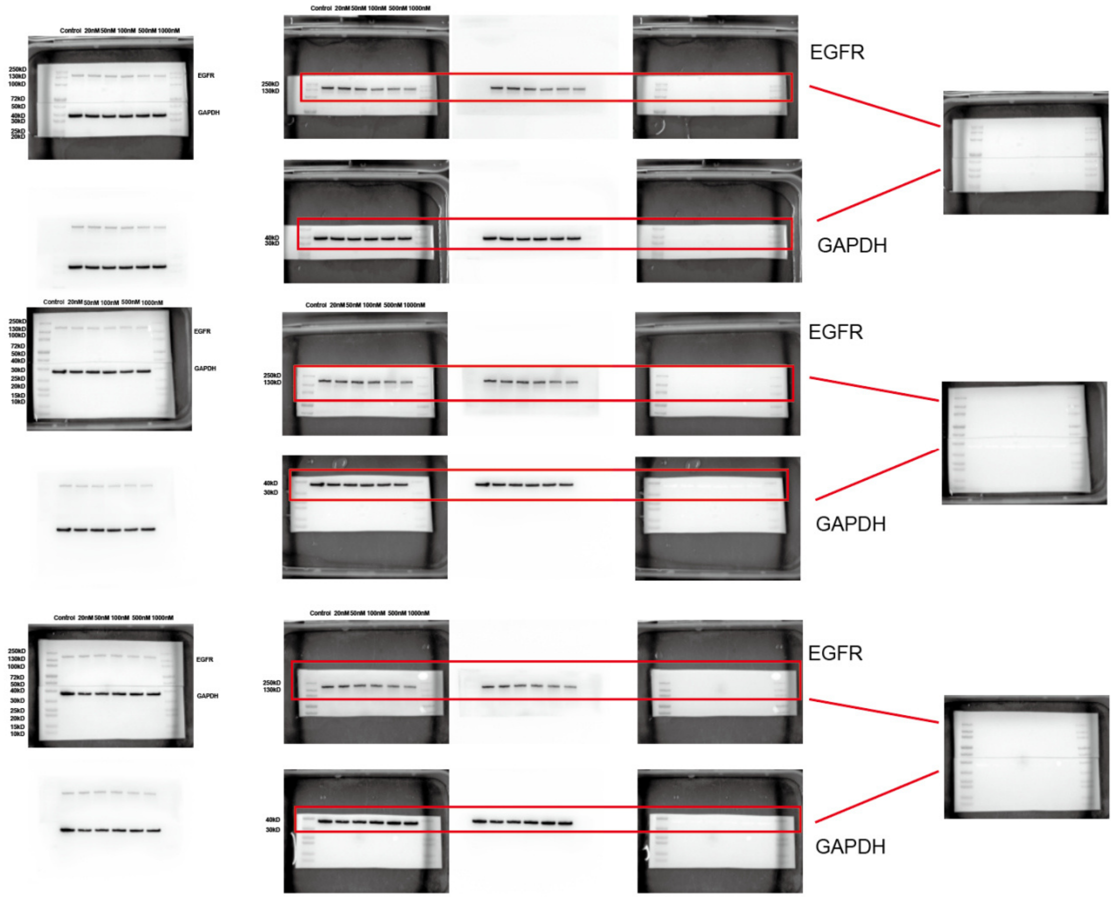


**Figure 2D MDA-MB-231 cells**

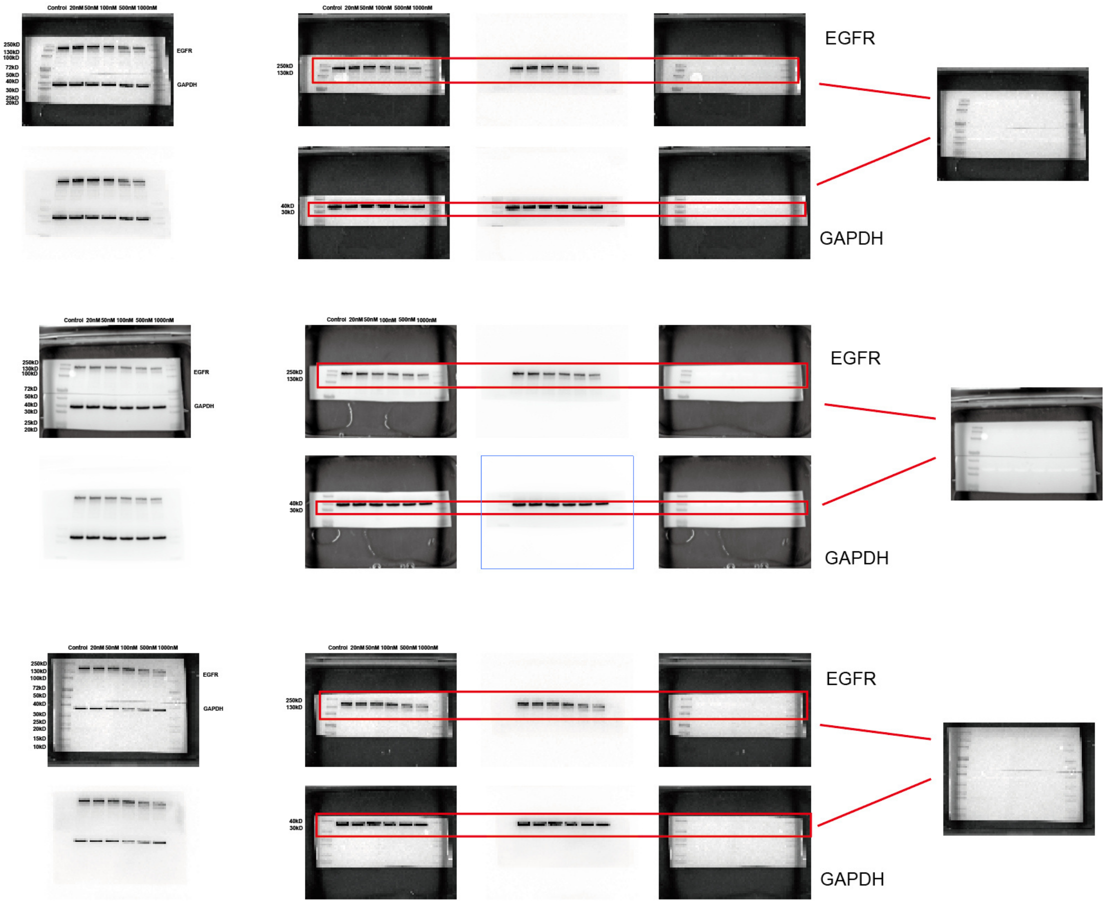


**Figure 2D NCI-N87 cells**

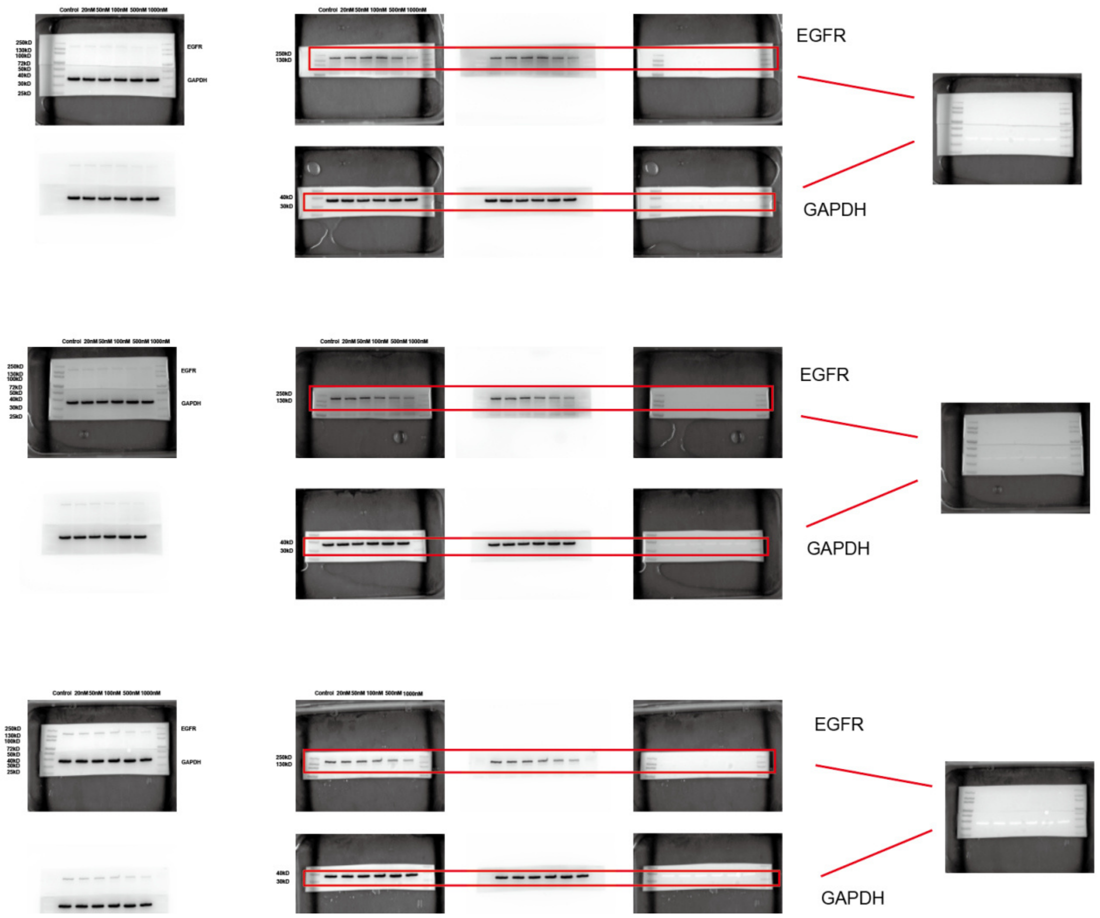


**Figure 2D BxPC-3 cells**

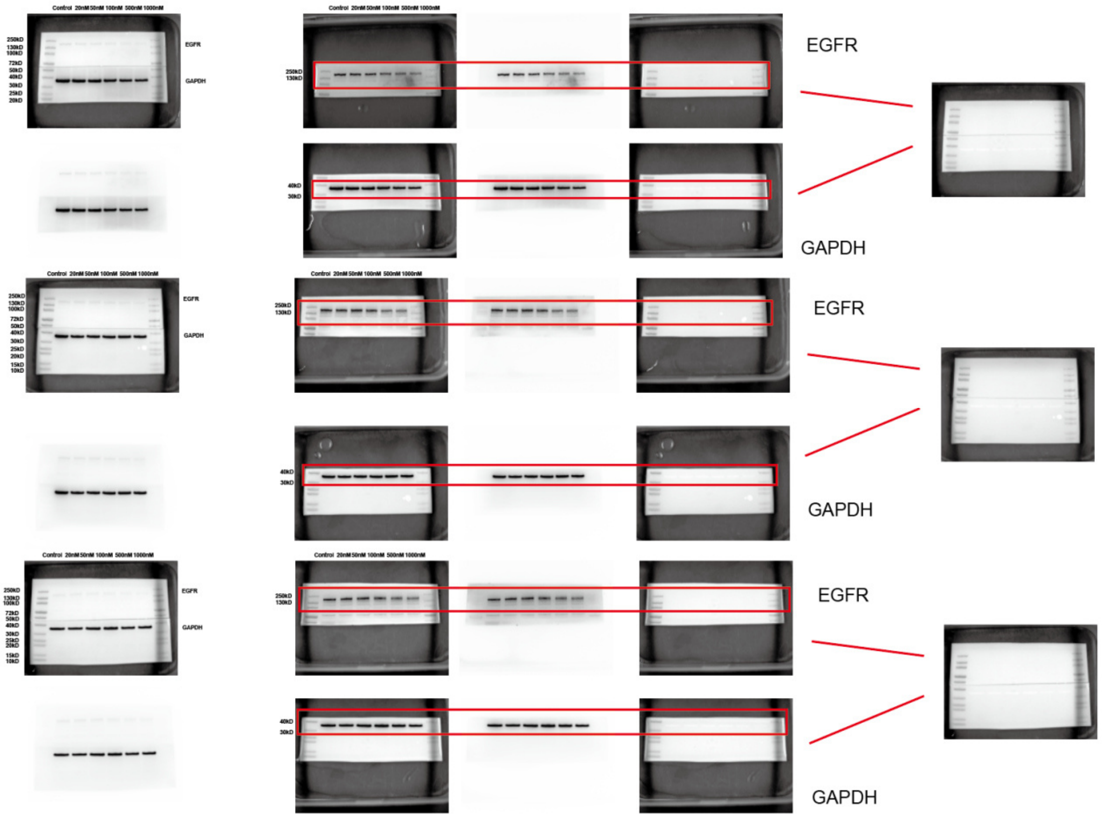


**Figure 2D NCI-H1975 cells**

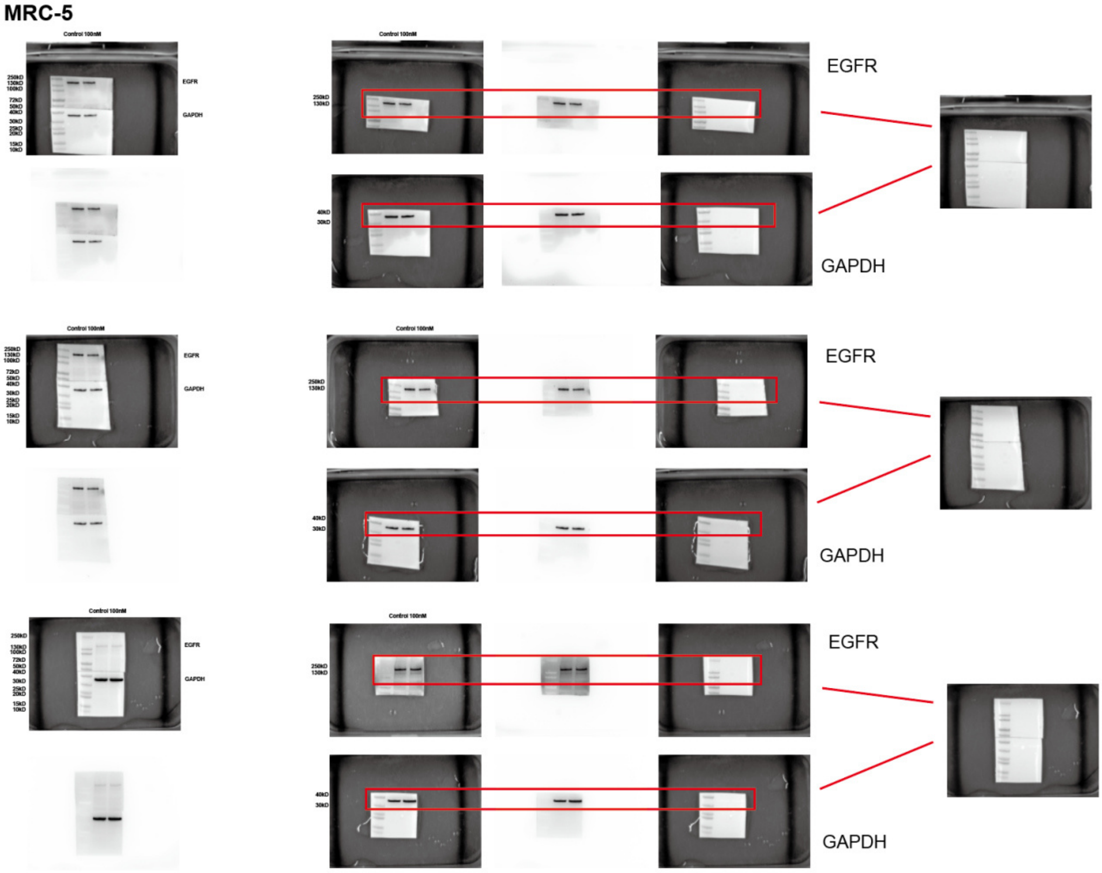


**Figure 2E MRC-5 cells**

HUVEC

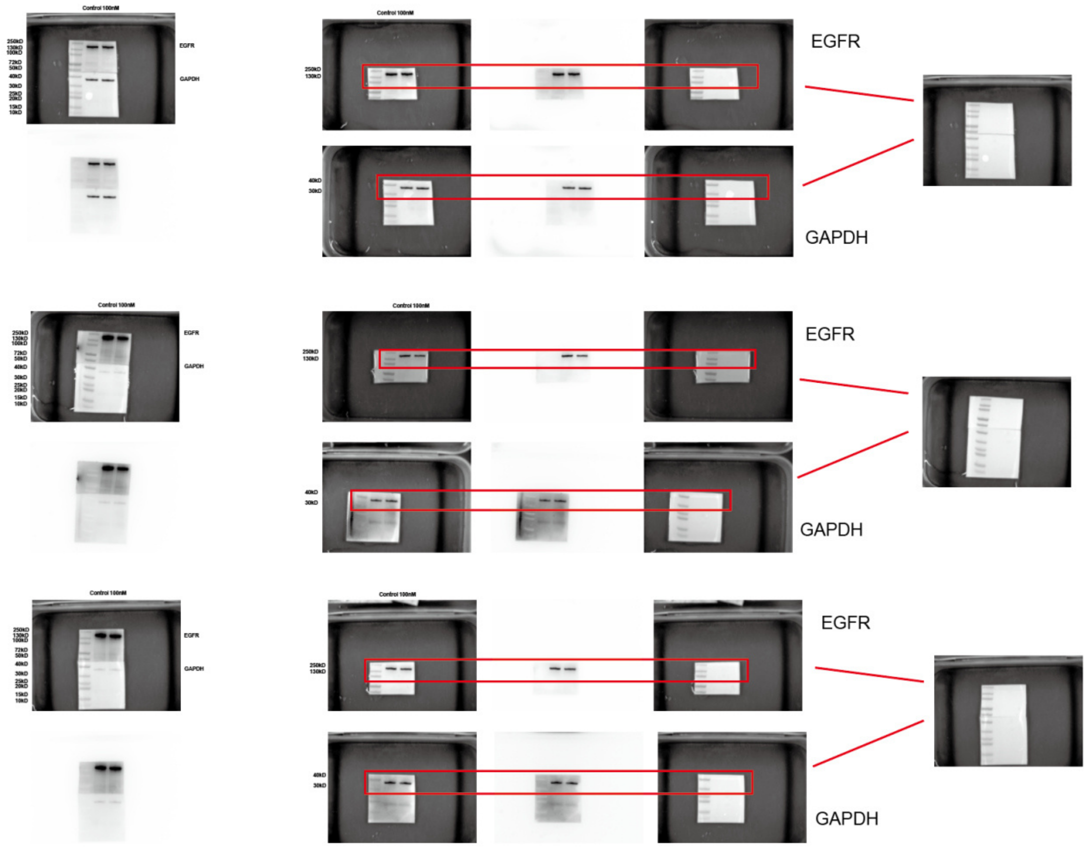


**Figure 2E HUVEC cells**

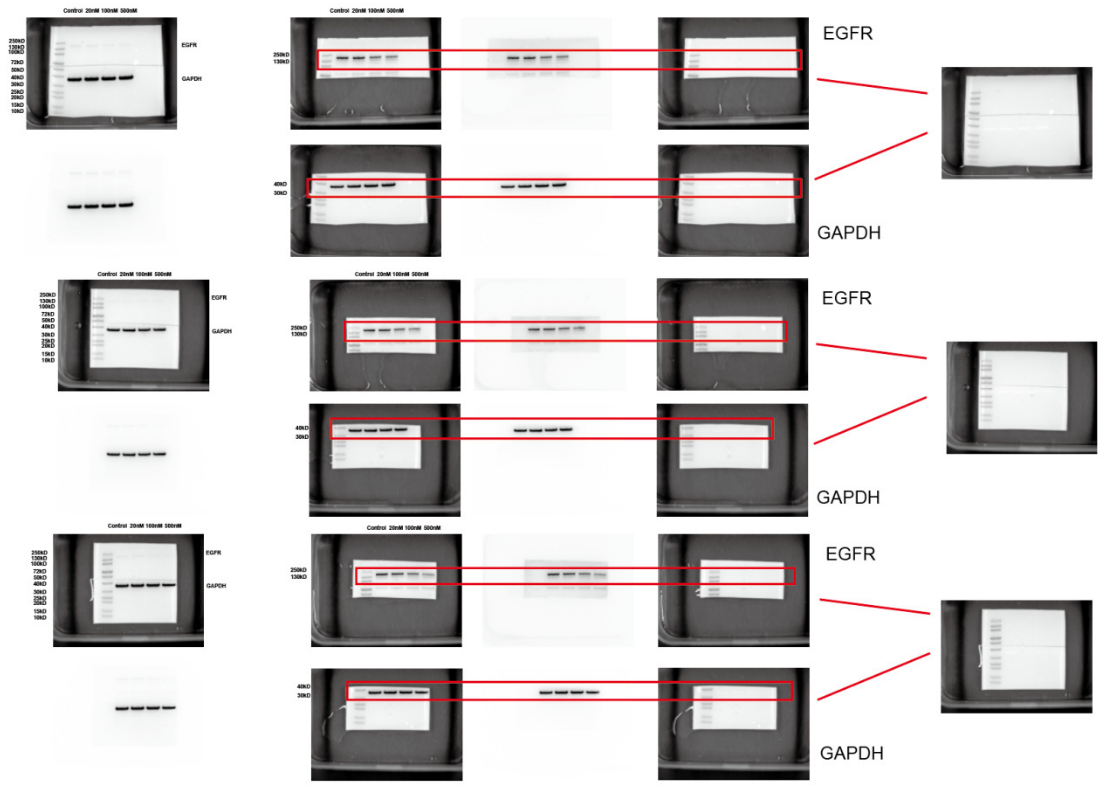


**Figure S3B HeLa cells**

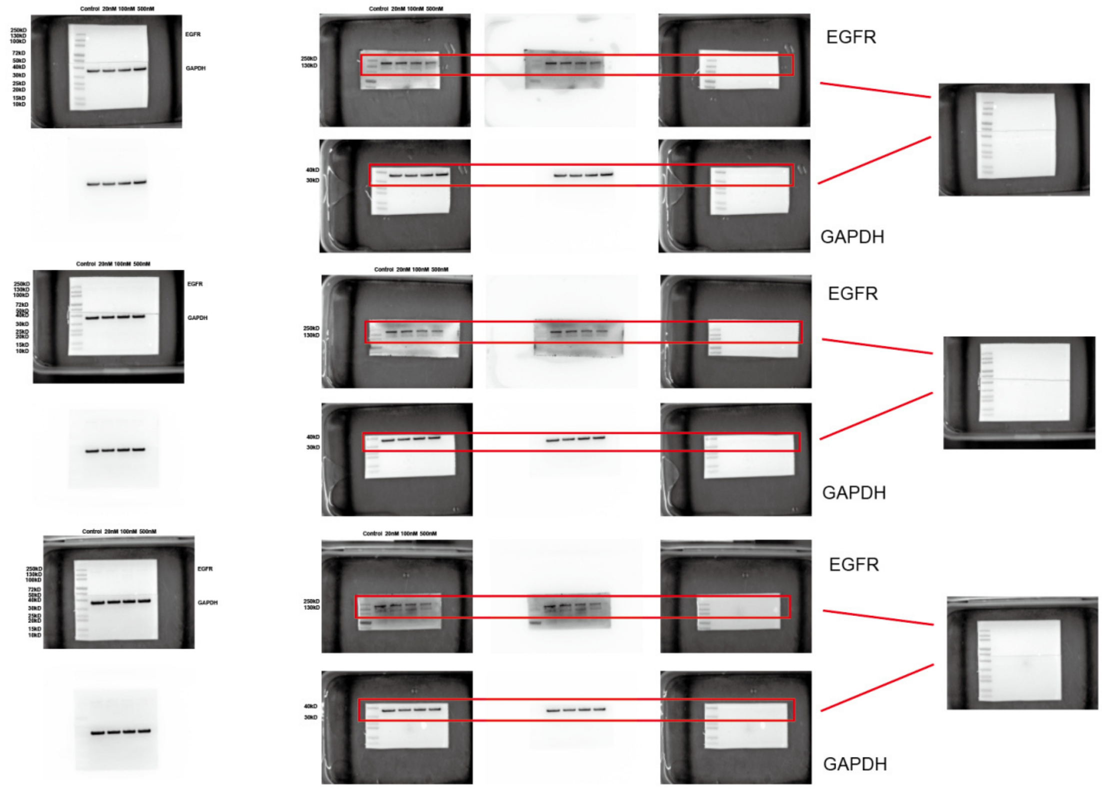


**Figure S3B H1975 cells**

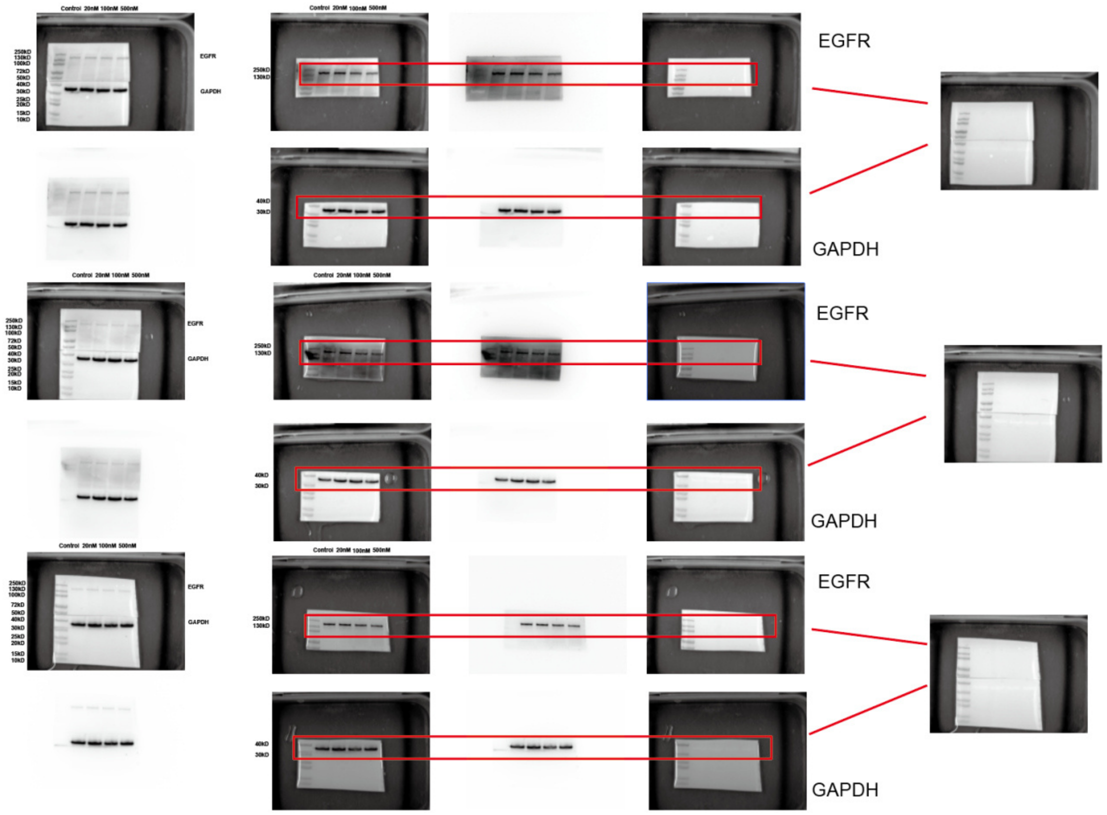


**Figure S3B SKBR3 cells**

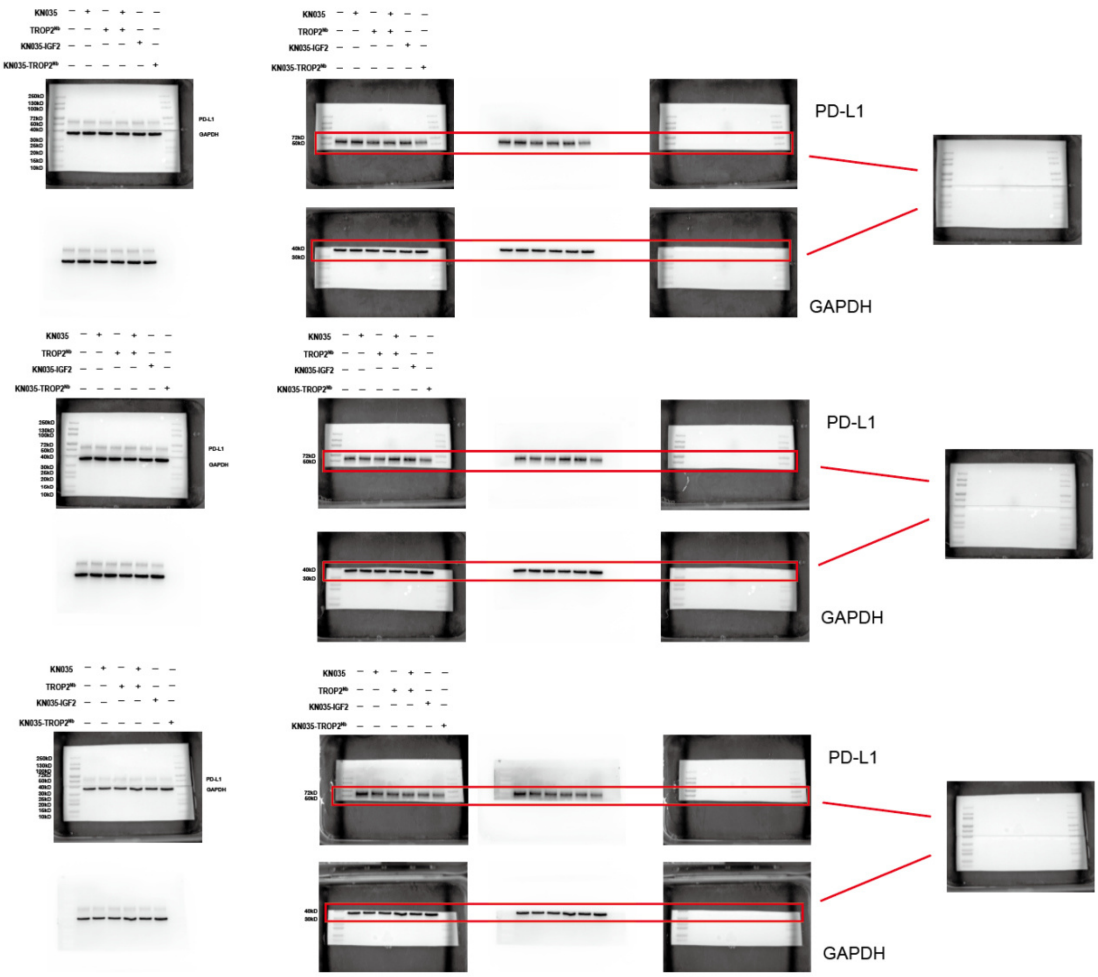
KN035
TROP2
KN035-IGF2
KN035-TROP2
PD-L1
GAPDH
PD-L1
GAPDH
PD-L1
GAPDH

**Figure 4C**

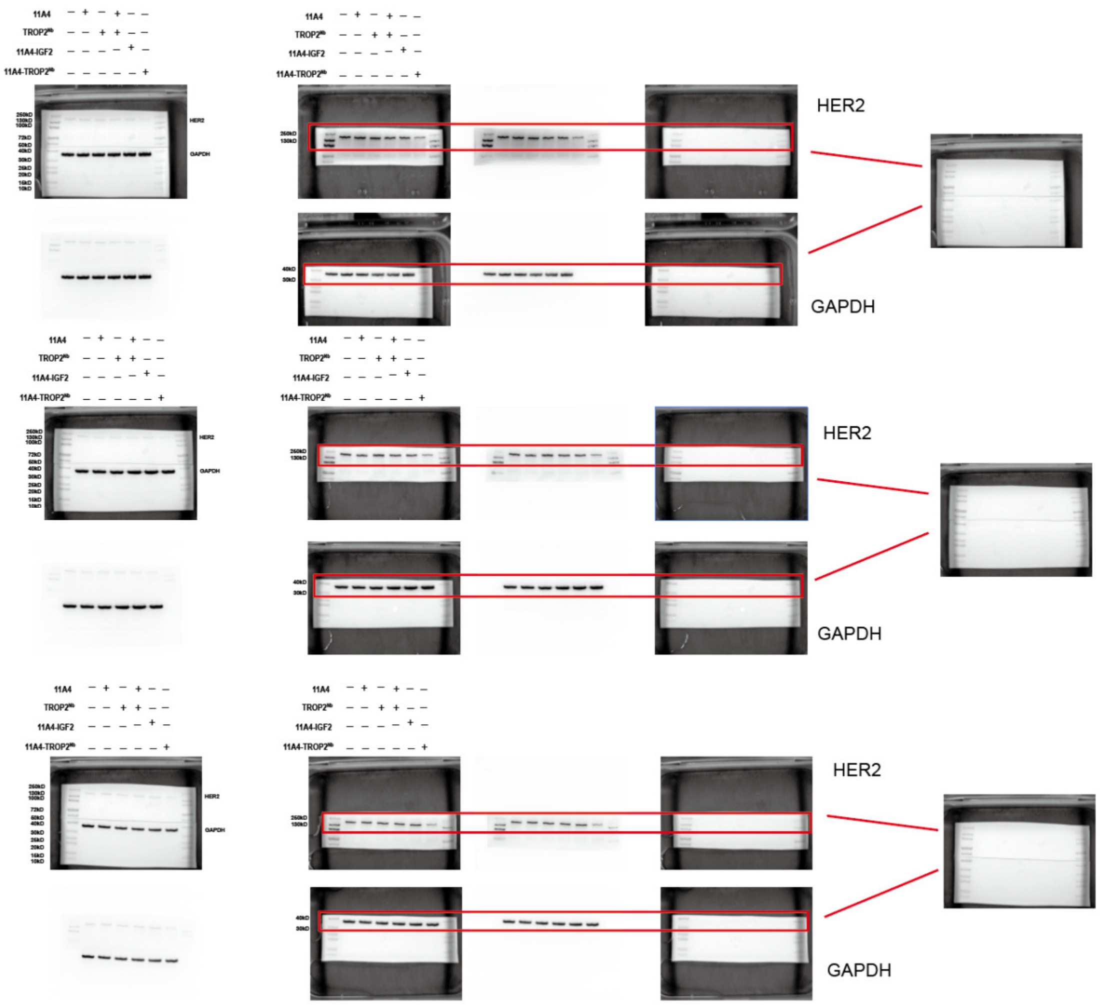
11A4
TROP2Ab
11A4-IGF2
11A4-TROP2Ab
HER2
GAPDH
HER2
GAPDH
HER2
GAPDH

**Figure 4E**

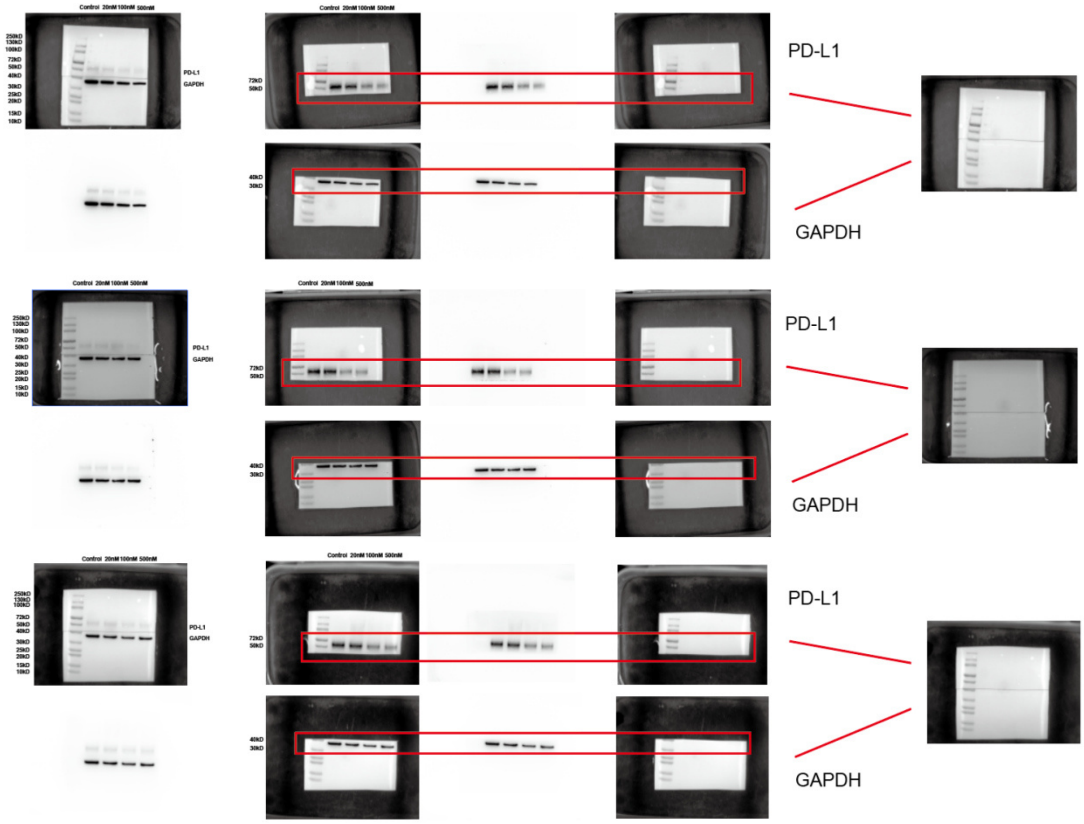


**Figure S4A**

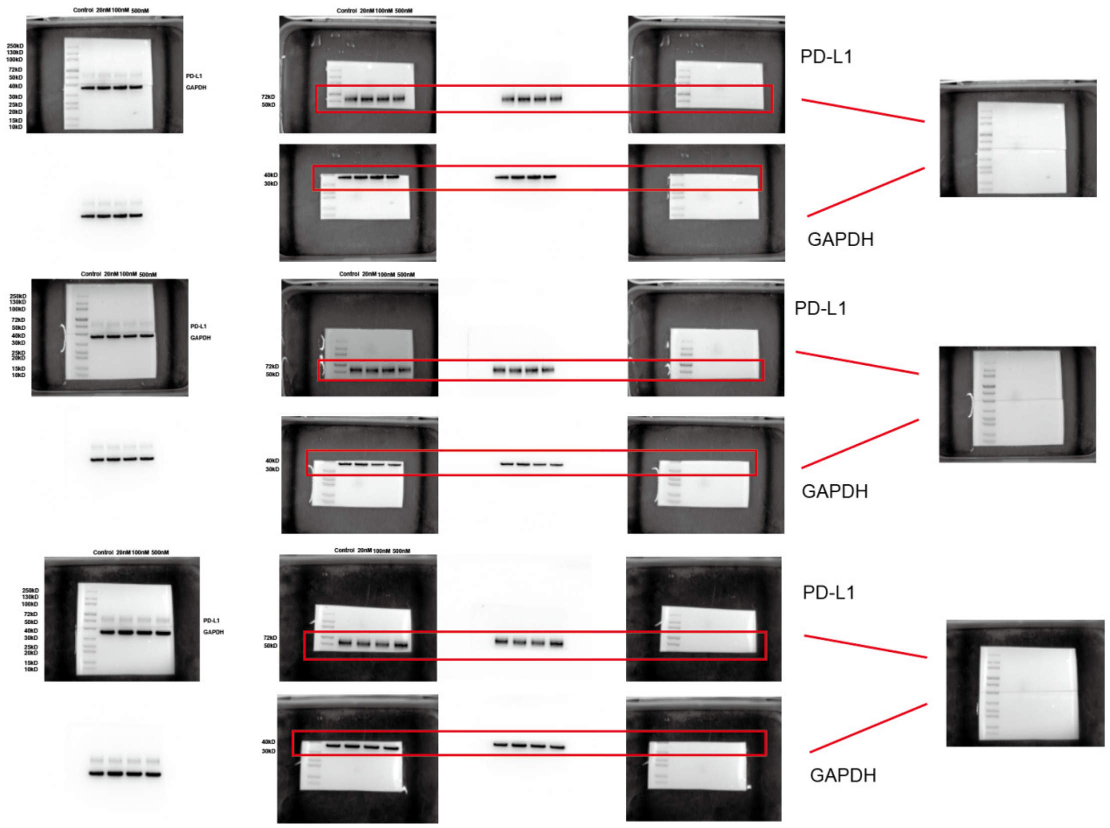


**Figure S4A**

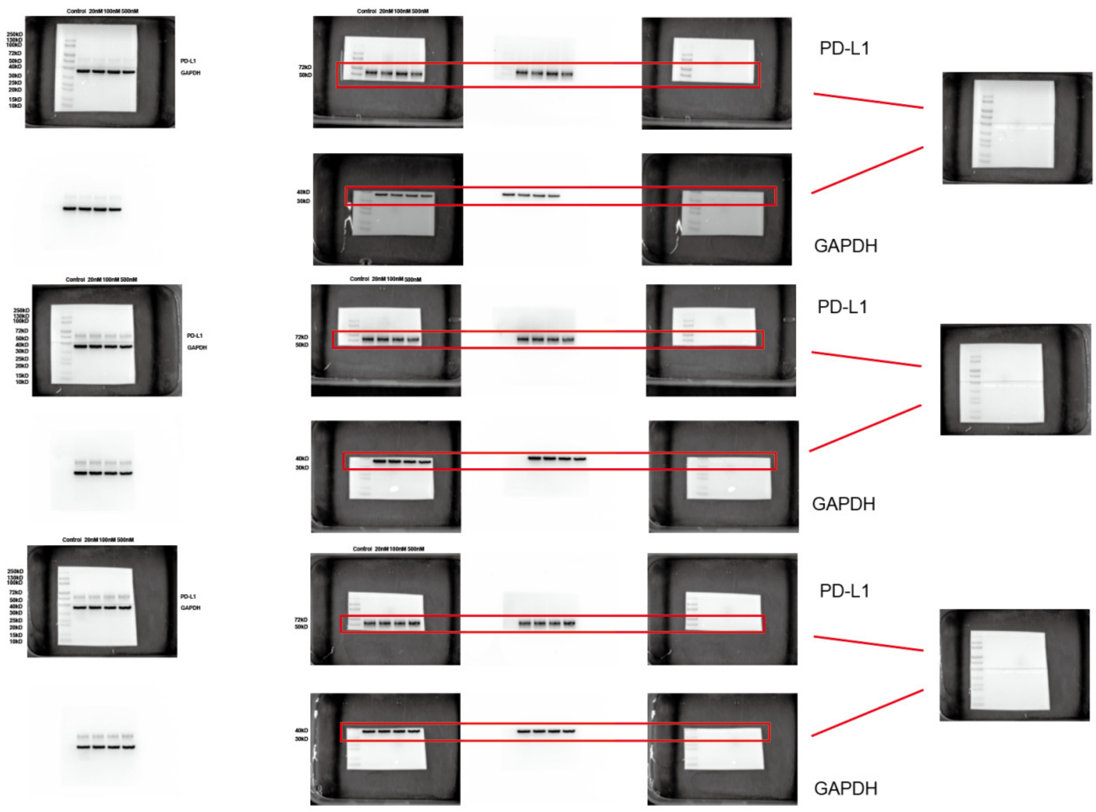


**Figure S4A**

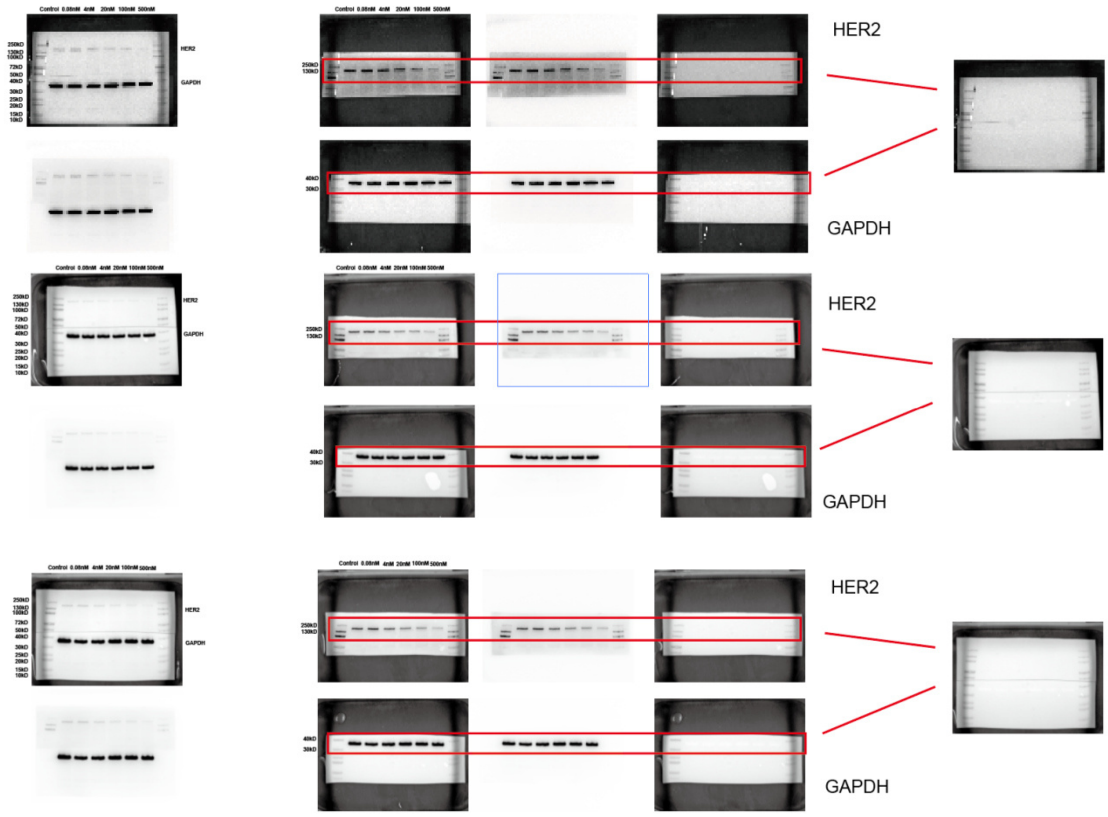


**Figure S4B**

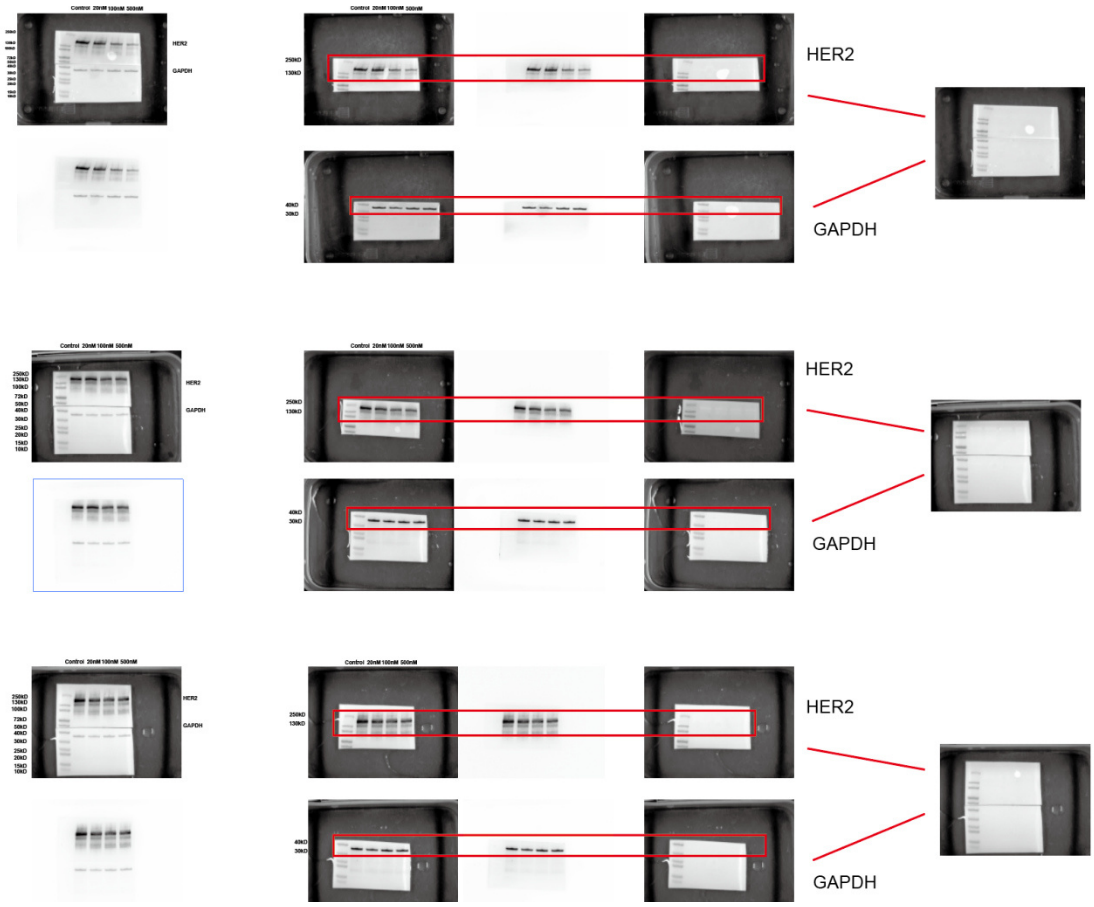

Control 20nM 100nM 500nM
HER2
GAPDH
250kD
130kD
40kD
30kD
HER2
GAPDH
HER2
GAPDH
HER2
GAPDH


**Figure S4C**

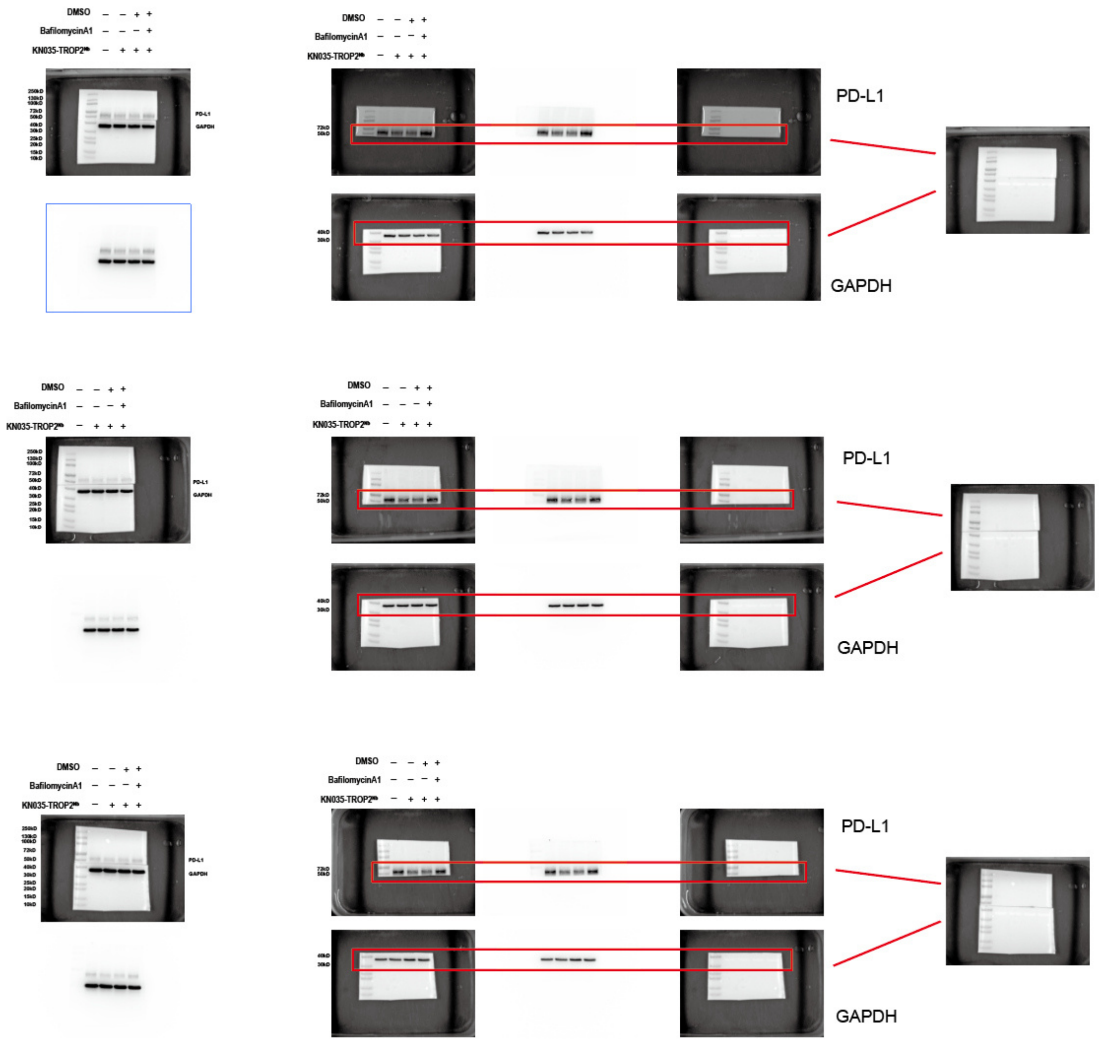


**Figure S4D**

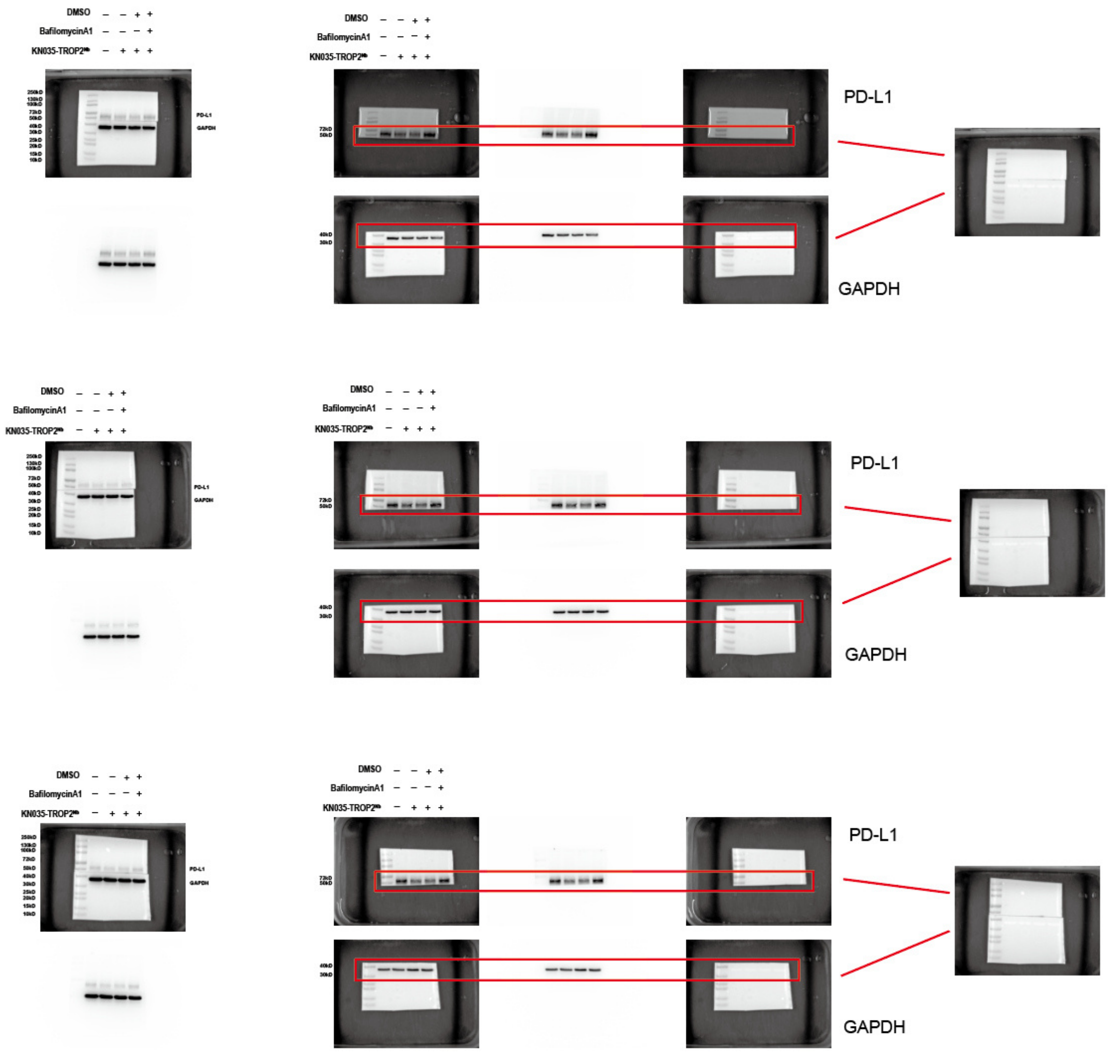


**Figure S4E**
